# An exhaustive map of binary memory-two direct reciprocity over symmetric two-player games

**DOI:** 10.64898/2026.09.05.749606

**Authors:** Martin A. Nowak

## Abstract

Direct reciprocity, a mechanism for evolution of cooperation, rests on a promise of the future: that the cost of cooperation will be returned in subsequent encounters. Whether reciprocity works is often asked as a question about evolutionary stability: can a population of cooperators resist invasion by mutants that defect? Here we answer this question exhaustively for a large, but finite strategy space projected onto an uncountable infinity of evolutionary games. We take the binary memory-two strategies, in which each of the sixteen possible two-round histories is answered by cooperation or defection up to a small error rate. We work out what every one of them achieves on its own and against every rival, in every symmetric two-player game. Alone, they realise 475 distinct patterns of play carrying 229 distinct cooperation rates. Against each other, a game supports between 299 and 22069 Nash equilibria, that is, resident strategies that no rare mutant can outperform. Efficient strategies, which are those that reach maximum payoff, are present as equilibria at every game, and for 3/8 of games efficient strategies constitute the only equilibria. Those results belong to the limit of a vanishingly small error rate. For any positive error rate the map changes: under 1/4 of all games support no equilibrium, over 3/8 support equilibria but none are efficient, over 1/8 support equilibria and all are efficient, and under 1/4 of games support equilibria of both kinds. Every share quoted here is an exact natural density over the plane of games. The map is the landscape over which any evolutionary dynamics in this strategy space moves.

## Introduction

Cooperation, in which one individual pays a cost so that another may benefit, is not something natural selection favours on its own; it needs a mechanism (1–4). Direct reciprocity is the mechanism built on repetition. Trivers introduced reciprocal altruism with examples from animal behaviour, cleaning symbioses in fish among them (1); it has since been documented across the animal kingdom, from blood sharing in vampire bats to predator inspection in sticklebacks (5, 6), and it pervades human life. When the same two individuals meet again, a strategy for the repeated game can make what it does now depend on what was done before, so that a cooperative move can be answered in kind and a defection can be punished (1, 7–14). How much of the past such a strategy may consult is the single most consequential modelling choice available (15–19).

The literature has answered the question of whether reciprocity sustains cooperation often in the language of equilibria (14, 17, 20–25). One asks whether a cooperative arrangement can resist invasion by a rare mutant — the evolutionary stability question of Maynard Smith and Price (26, 27) — and the folk theorem guarantees that in a sufficiently long game very many arrangements can (28–32). The fully cooperative arrangements that resist invasion are the partner strategies of the literature (12, 20, 21). This paper asks the usual question completely rather than by example, and asks alongside it what is available at all. We fix a strategy space large enough to be interesting and small enough to be enumerated in full. Then we compute, for every game and every strategy at once, what each strategy achieves, what it sustains against every rival, whether it is a Nash equilibrium somewhere on the map and whether that property survives trembling hands.

### Why binary memory-two strategies

We restrict attention to strategies whose cooperation probabilities take only two values, *ɛ* and 1 *− ɛ*, for a small error rate *ɛ*. In this setting (8, 9), the parameter *ɛ* can be read as Selten’s trembling hand (33). It is also where the interesting strategies live: when allowing the full interval [*ɛ,* 1 *− ɛ*], evolutionarily stable strategies exist at its boundary (34, 35). In a recent study of evolutionary dynamics, binary memory-two strategies resolve every social dilemma examined (36). Restricting to two values per component turns an uncountable space into the 2^16^ = 65536 objects of Lindgren’s binary genome (37), and an exhaustive answer becomes possible where before only sampling was. The space itself is not new and has been swept before (17, 19); what is new here is the map computed over the space of symmetric two-player games. Memory two is not arbitrary either. It is the shortest memory at which a strategy can tell an isolated error from a second defection, and the shortest at which two players can alternate perfectly, which is the social optimum above the switch line introduced below; binary memory-one strategies can do neither, and in the Snowdrift game evolution with memory one fails where memory two succeeds (36). Memory three has 2^64^binary strategies, and an exhaustive census of every strategy against every rival is out of reach. Memory two is therefore the one level at which the space is both rich enough to contain the behaviour that matters and small enough to be surveyed completely.

### Strategies

A memory-*n* strategy consults the outcomes of the last *n* rounds. Since each round ends in one of the four outcomes *CC*, *CD*, *DC*, *DD*, a memory-*n* rule is a list of 4*^n^* cooperation probabilities. For *n* = 2 that is sixteen numbers, one for each pair (last round, round before). Two players using such rules generate a Markov chain on sixteen states. We study infinitely repeated games with no discounting of the future. The payoff is the long-run average per round. When every probability lies strictly between zero and one the chain has a unique stationary distribution, and payoffs and cooperation rates are read from it (**Methods**). We write a strategy as the sixteen-bit string of its components, ordered by (last round, round before), so that *ALLC* is sixteen ones, *ALLD* sixteen zeros, and *TFT*, *WSLS* and *Grim* are particular strings among the 65536.

### Games

Every symmetric two-player, two-action game in which mutual cooperation beats mutual defection can be brought by a positive affine change of scale to the payoff matrix

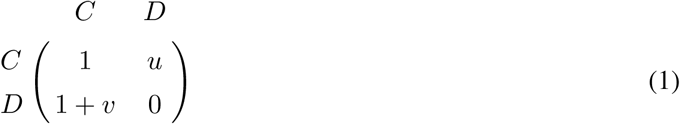

Mutual cooperation earns 1, mutual defection earns 0, and the two parameters *u* and *v* locate the game in a plane (25, 38, 39), over which the strategic properties of the *one-shot* game have themselves been mapped (40). The four sign quadrants are the four classical dilemmas: Prisoner’s Dilemma (*u <* 0, *v >* 0), Snowdrift (*u >* 0, *v >* 0), Stag Hunt (*u <* 0, *v <* 0) and Harmony (*u >* 0, *v <* 0), meeting at the origin (41, 42). Working in this plane rather than at a handful of chosen games is what makes the results below statements about all games rather than about examples.

The best that two players can jointly achieve per round is *E*_max_ = max*{*1, (1 + *u* + *v*)*/*2*}*. The two branches matter. Below the line *u* + *v* = 1 the optimum is mutual cooperation. Above it, the optimum is alternation: the pair takes turns at *CD* and *DC*, and the average of the two asymmetric payoffs beats mutual cooperation (43–45). That line crosses three of the four quadrants, and almost everything below turns on which side of it a game lies. We call a strategy *efficient* at a given game when a population of it attains *E*_max_ there as the error rate vanishes.

### Drawing all games at once

The plane (1) is unbounded and every result below is a statement about the whole of it, so any window *|u|, |v| ≤ M* would leave part of the answer outside the picture — and not an inert part, since at a positive error rate the games carrying the most equilibria are in the far field. Figures 2 to 5 therefore draw the plane compactified rather than cropped, by the map

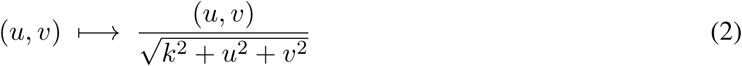

The parameter *k* fixes how the page is divided between the near field and the far: the games with *|*(*u, v*)*| ≤ k* occupy exactly half the disk’s area, so a smaller *k* magnifies the neighbourhood of the origin and a larger one gives more of the picture to the far field. Figures 2 to 5 are drawn at *k* = 4, and the two single-game panels of Figure 2 at *k* = 1.5. The map carries R^2^ onto the open unit disk and the directions at infinity onto its rim. It fixes the origin and keeps every line through the origin straight, so the two axes stay diameters, the four dilemmas stay four quadrants of equal area, and each family can be named on the rim of its own sector; every other line becomes a conic arc. Nothing is cut off. What the map costs is uniformity of scale — area far out is compressed, which is why a raster drawn on the disk falls a little short of maxima that are attained only in the far field. A count of equilibria at a game is unaffected by the map. A share of the plane is a subtler matter, and the distinction decides how the shares below are to be read: a share of the drawn *area* depends on *k* exactly as it would depend on the half-width of a window, whereas a share of the *directions*, read at the rim, does not. Every share reported below is of the second kind (**Methods**).

## Results

### What a strategy achieves on its own

Before asking what is stable, we ask what is available at all. A population in which everyone uses the same binary memory-two strategy settles, as the error rate vanishes, on a distribution over the four round outcomes, and from it on a cooperation rate. Enumerating all 65536 strategies exactly gives 475 distinct distributions carrying 229 distinct cooperation rates (Figure 1; **SI** §2). The two numbers are different and the difference is the point: a cooperation rate is a single number extracted from a pattern of play, and thirteen behaviourally distinct residents share the rate 2*/*3 while fifteen share 1*/*2. Knowing how much a population cooperates does not tell you how it cooperates.

**Figure 1:**
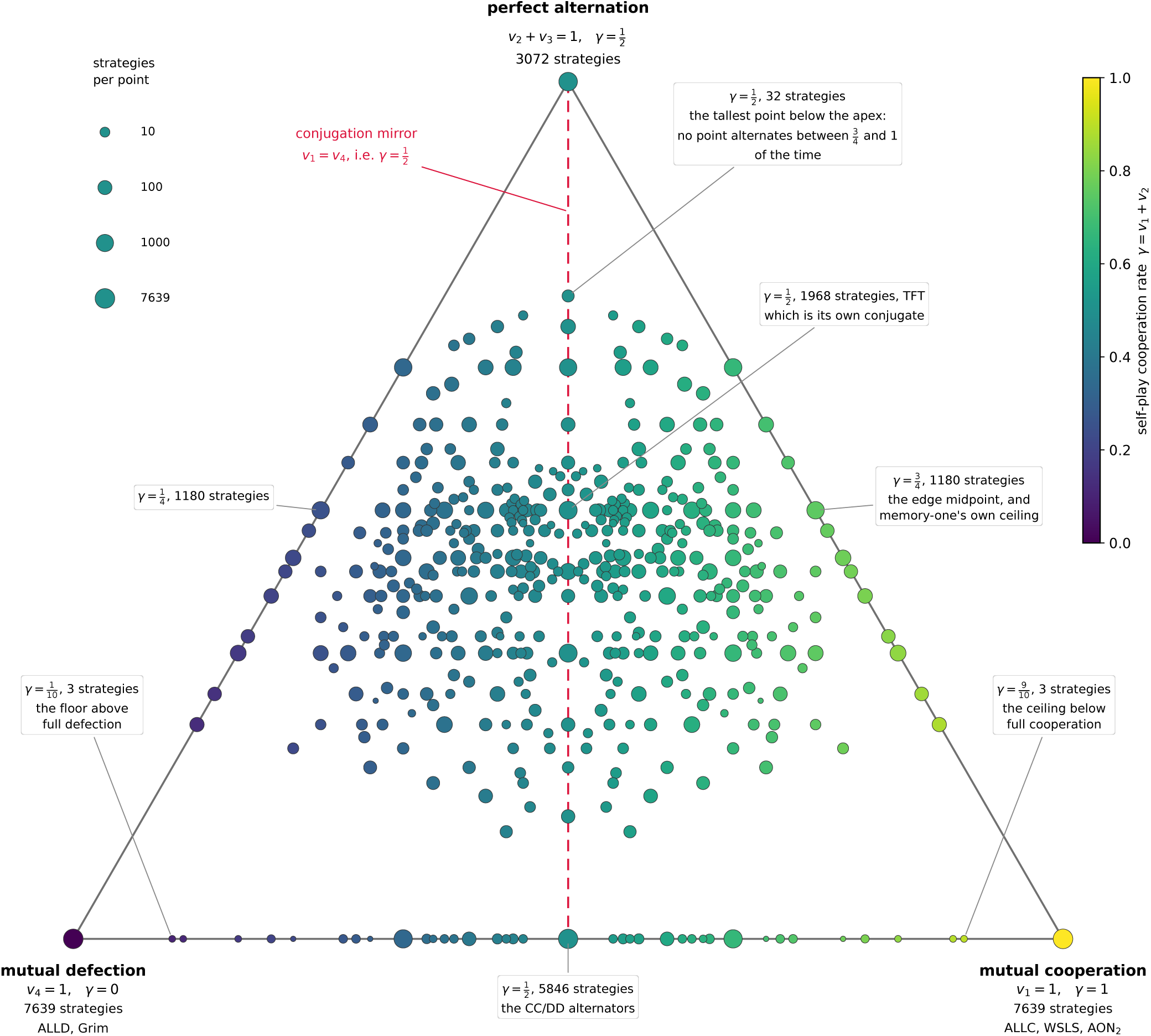
The 475 self-play points of the 65536 binary memory-two strategies. Each strategy, played against a population of itself, settles as the error rate vanishes on a distribution (*v*_1_*, v*_2_*, v*_3_*, v*_4_) over the round outcomes *CC*, *CD*, *DC*, *DD*; because the two players are interchangeable, *v*_2_ = *v*_3_, so the distribution has two free coordinates and is drawn in a triangle. Mutual defection is the bottom left corner, mutual cooperation the bottom right and perfect alternation the apex, so height measures how much of the time the two players are doing different things, and the self-play cooperation rate *γ* = *v*_1_ + *v*_2_ increases from left to right along the base. Marker area is proportional to log_10_ of the number of strategies realising a point, and colour is *γ*. Seven points are called out with their rate and the number of strategies at them. The dashed line is the conjugation mirror, the reflection induced by relabelling *C* and *D*: it fixes the apex, exchanges the two bottom corners — which is why both carry the same 7639 strategies — and sends *γ* to 1 *− γ*. The axis it fixes is *v*_1_ = *v*_4_, which is the same condition as *γ* = 1*/*2.

Two features of this catalogue recur throughout. First, it is exactly symmetric: exchanging the roles of *C* and *D* maps the strategy set onto itself and reverses the cooperation rate. The 475 points fall into 230 mirror pairs and 15 self-conjugate points. Second, there is a gap. Walking down from full cooperation the achievable rates run 1, 9*/*10, 8*/*9, 7*/*8*, …*, and so no population of a single binary memory-two strategy cooperates at a rate strictly between 9*/*10 and 1. Memory-one has the same feature with a lower ceiling, 3*/*4, which can be read off published tables of its limiting self-play distributions (46). Adding an extra round of memory raises the best sub-perfect performance but does not close the gap.

### The equilibria of every game at once

A strategy is a Nash equilibrium of a given game when no deviation earns more against a population using it. In evolutionary terms the population is a monomorphic population of residents, a deviation is a rare mutant, and the condition says that natural selection does not favour the mutant: it is the first of Maynard Smith’s two conditions for an evolutionarily stable strategy (27), the one that decides whether a resident can be invaded at all. The second, which decides the fate of a mutant that ties, is a matter for the ties, and we shall see that in the limit every equilibrium has them (**SI** §4). Because payoffs are read from a stationary distribution that does not depend on *u* and *v*, and because those parameters enter the payoff linearly, the gain from any deviation is an affine function of the game. Each deviation therefore rules out a half-plane of games, and the set of games at which a given strategy is an equilibrium is a convex polygon in the (*u, v*) plane (47, 48) (**SI** §3). One computation per strategy settles every game simultaneously — including every game off any lattice one might have chosen to examine.

What has to be tested depends on whether we consider a fixed error rate, *ɛ*, or the limit of *ɛ →* 0. The difference is not a technicality (**SI** §§3–4). At a fixed error rate every history occurs, so a deviator faces an average-reward Markov decision process on the sixteen states in which every feasible transition is bounded below by *ɛ*^2^ and every state is reached from every other in two rounds; unimprovability is then equivalent to optimality, and the sixteen single-component deviations decide the test. As the error rate vanishes that reduction fails: a component governing a state that is never visited contributes nothing, its constraint becomes vacuous, and the test admits strategies it should not. The limit computation therefore compares every resident against all 65535 rivals, with exact ties admitted (14, 19). Both cases, a fixed error rate *ɛ* and the limit *ɛ →* 0, are of interest. We compute both.

Figure 2 shows the map of all Nash equilibria. For the limit *ɛ →* 0, we find that 23861 strategies are equilibria on a two-dimensional set of games, 3774 along a line of games, 4875 at a single game, and 33026 nowhere. The full-dimensional regions are less varied than that count suggests: the 23861 collapse to 944 distinct polygons bounded by 254 distinct lines. Every game in the plane supports at least 299 equilibria. The most any game supports is 22069, attained not at some exceptional game but on an unbounded region of positive area, the Stag Hunts with *u ≤ −*4 and *v ≤ −*2. The lower strata are more concentrated still: the 3774 strategies that are equilibria only along a line share just 165 intervals on 44 distinct lines, so a tie is almost never a strategy’s own, and the 4875 that are equilibria at a single game sit at only 52 games, every one of them within *|*(*u, v*)*|* = √5 of the origin (**SI** §5).

**Figure 2:**
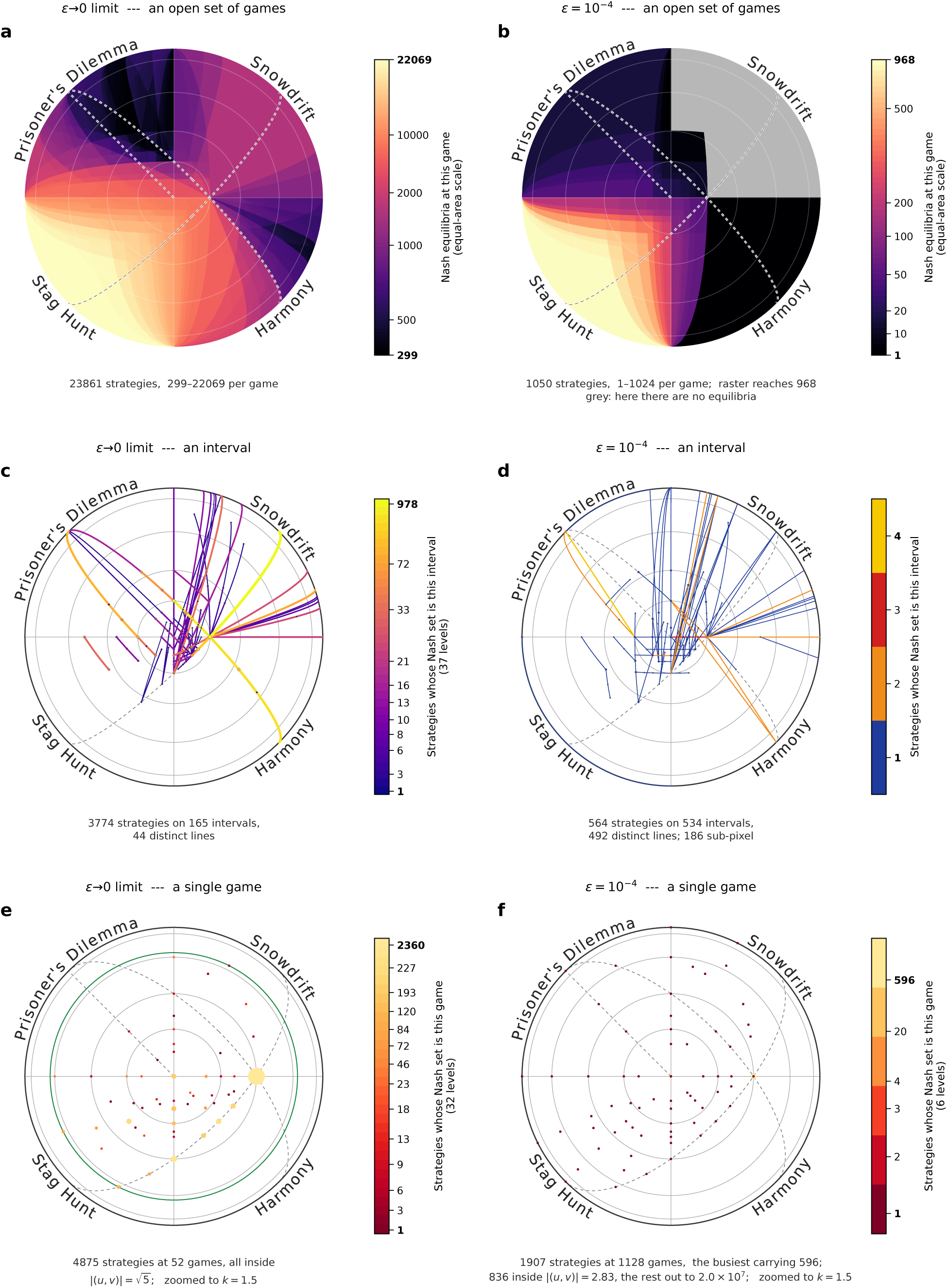
Where a strategy is an equilibrium, and how much of that survives an error rate. Columns are the two notions of equilibrium — left, the vanishing-error limit; right, *ɛ* = 10*^−^*^4^ — and rows are the dimension of the set of games at which a strategy is an equilibrium: an open set, only a line, only a single game. Each disk is the whole plane, its rim infinity and each point of it a direction, the four quadrants the four dilemmas. Every panel has its own colour scale, and the top row counts a different thing from the two below it: **a** and **b** count how many strategies are an equilibrium *at* each game, while **c** to **f** count how many have that interval or that game as their whole equilibrium set. **a**, 23861 strategies hold an open set; **b**, 1050 still do at *ɛ* = 10*^−^*^4^, and grey marks the games where no strategy is an equilibrium at all. **c**, 3774 hold only a line, sharing just 44 of them; **d**, 564 still do, scattered over 492. **e**, 4875 hold only a single game, and those games are only 52, every one within *|*(*u, v*)*|* = √5 of the origin — exactly √5, which three of them attain and the green circle marks; **f**, 1907 still do, at 1128 games, 596 of them at the single game (*u, v*) = (9999*/*9998, 1*/*9998). Those 1128 are bounded too, but far more widely, and in three shells: 836 of them lie within *|*(*u, v*)*|* = 2.83, none at all between there and 1.6 *×* 10^3^, and the farthest is at 2.0 *×* 10^7^, so the outer 292 land against the rim. Both single-game panels are drawn at the closer scale *k* = 1.5 and the other four at *k* = 4. **f**’s bar has six entries because those 1128 games carry only six distinct counts — 1, 2, 3, 4, 20 and 596 — and **d**’s has four for the same reason.

No strategy is an equilibrium at every game, and none commands more than half the plane. Every full-dimensional region is unbounded, and the largest are the 566 whose entire region is a single half-plane; two non-redundant constraints already make a region smaller than that. Exactly one of those half-planes has the origin strictly inside it, and it belongs to just two strategies, given by *CCCC DDDC DCCC ∗DCD*, where the star is the single component in which the two differ (**SI** Table 1). Their region is exactly *u* + *v ≤* 1: they are equilibria at every game at which mutual cooperation is the social optimum, and stop being equilibria exactly where full cooperation stops being the best thing to want. Both cooperate fully, and the component they differ in governs a state that only two simultaneous errors can reach, which is why the difference costs neither of them anything. *ALLC* and *ALLD* hold half-planes too: their regions are the half-planes *v ≤* 0 and *u ≤* 0, each passing through the origin, and shared with 294 and 72 other strategies respectively. *WSLS* holds the quadrant *u ≤* 1, *v ≤* 1, and *AON*_2_ — which cooperates only when both of the last two rounds were unanimous — the slightly larger *u ≤* 1, *v ≤* 2. *TFT* holds nothing at all: it is one of the 33026 strategies that are an equilibrium at no game whatever, exactly as for memory-one (**SI** §5).

**Table 1:**
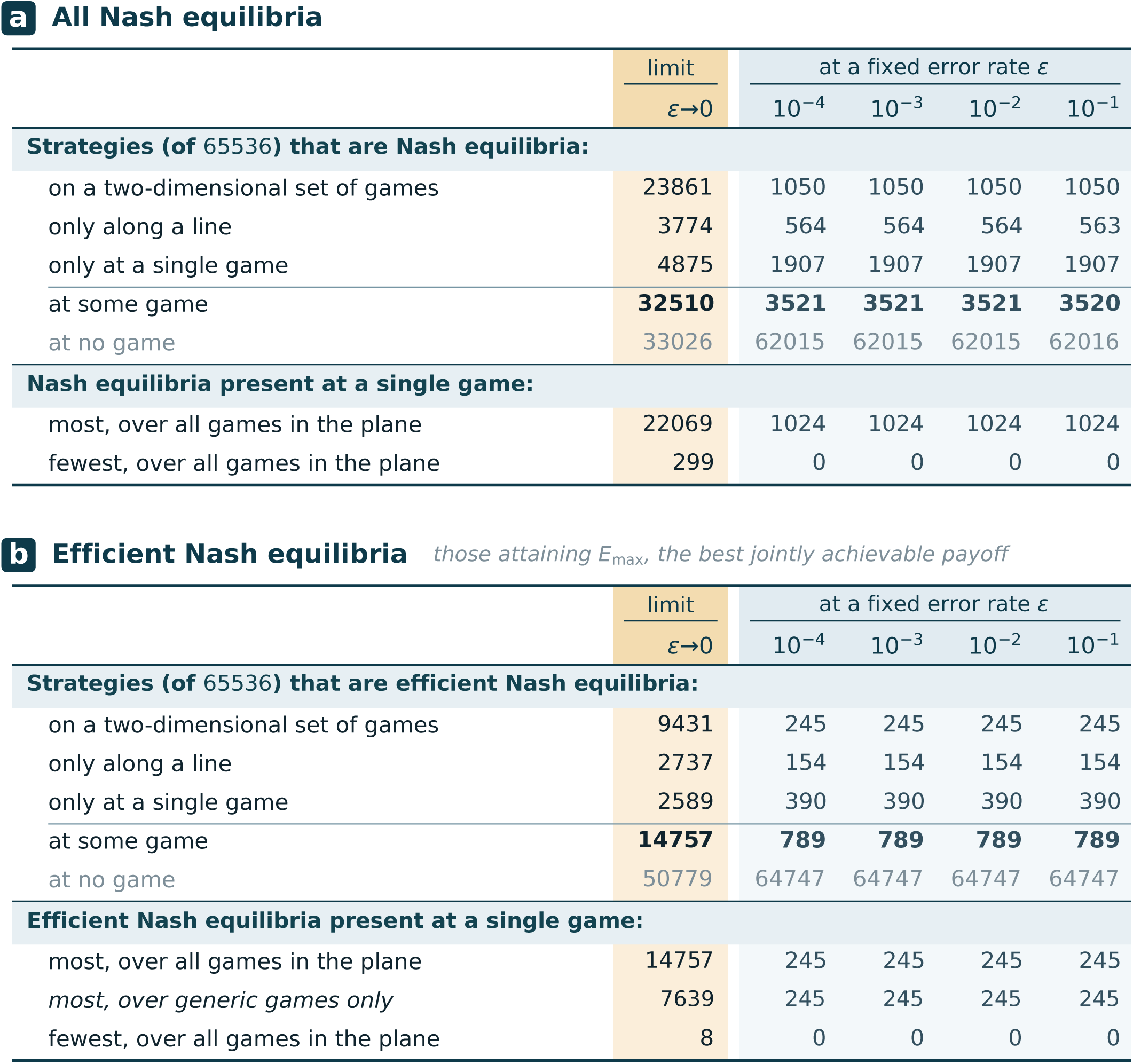
How many equilibria there are, in the limit and at four error rates. **a**, all Nash equilibria; **b**, only those that attain *E*_max_, the best jointly achievable payoff. Columns are the vanishing-error limit and then *ɛ* = 10*^−^*^4^, 10*^−^*^3^, 10*^−^*^2^, 10*^−^*^1^. The upper block of each panel counts *strategies*, of all 65536, by the set of games at which the strategy is an equilibrium — a two-dimensional set, only a line, only a single game, at some game, at no game — the last two summing to 65536 in every column. The lower block counts *equilibria at one game*, maximised and minimised over the whole plane. In **b** the *at some game* entry is also that panel’s limit maximum: every strategy that can ever be efficient is an efficient equilibrium somewhere. At a positive error rate that subtotal collapses to 789, the same at all four. And **b** carries one row **a** does not, *most, over generic games only*, because **b**’s limit maximum is attained at a single game and is therefore a null set, whereas **a**’s is attained on a region of positive area. Every entry is exact.

| | limit | at a fixed error rate $\varepsilon$ | | | |
| --- | --- | --- | --- | --- | --- |
| | $\varepsilon \rightarrow 0$ | $10^{-4}$ | $10^{-3}$ | $10^{-2}$ | $10^{-1}$ |
| **Strategies (of 65536) that are Nash equilibria:** |  |  |  |  |  |
| on a two-dimensional set of games | 23861 | 1050 | 1050 | 1050 | 1050 |
| only along a line | 3774 | 564 | 564 | 564 | 563 |
| only at a single game | 4875 | 1907 | 1907 | 1907 | 1907 |
| at some game | **32510** | **3521** | **3521** | **3521** | **3520** |
| at no game | 33026 | 62015 | 62015 | 62015 | 62016 |
| **Nash equilibria present at a single game:** |  |  |  |  |  |
| most, over all games in the plane | 22069 | 1024 | 1024 | 1024 | 1024 |
| fewest, over all games in the plane | 299 | 0 | 0 | 0 | 0 |

**b Efficient Nash equilibria** *those attaining $E_{\max}$ , the best jointly achievable payoff*
| | limit | at a fixed error rate $\varepsilon$ | | | |
| --- | --- | --- | --- | --- | --- |
| | $\varepsilon \rightarrow 0$ | $10^{-4}$ | $10^{-3}$ | $10^{-2}$ | $10^{-1}$ |
| **Strategies (of 65536) that are efficient Nash equilibria:** |  |  |  |  |  |
| on a two-dimensional set of games | 9431 | 245 | 245 | 245 | 245 |
| only along a line | 2737 | 154 | 154 | 154 | 154 |
| only at a single game | 2589 | 390 | 390 | 390 | 390 |
| at some game | **14757** | **789** | **789** | **789** | **789** |
| at no game | 50779 | 64747 | 64747 | 64747 | 64747 |
| **Efficient Nash equilibria present at a single game:** |  |  |  |  |  |
| most, over all games in the plane | 14757 | 245 | 245 | 245 | 245 |
| most, over generic games only | 7639 | 245 | 245 | 245 | 245 |
| fewest, over all games in the plane | 8 | 0 | 0 | 0 | 0 |

For *ɛ* = 10*^−^*^4^ we find a different map. Only 1050 strategies hold an open set of games, and the games that support no equilibrium at all are confined to Snowdrift, where they are nearly everything: a neighbourhood of the origin still carries equilibria, and beyond it, away from finitely many lines, only four thin pieces do, and far out in their directions what survives is never efficient. What confines them is a pair of half-planes that can be written down by hand: at a positive error rate *ALLC* and *ALLD* are equilibria on exactly (1 *−ɛ*)*v ≤ ɛu* and (1 *− ɛ*)*u ≤ ɛv*, each the limit’s half-plane tilted off its own axis by *τ* = arctan[*ɛ/*(1 *− ɛ*)]. Together the two cover the whole plane but for a cone inside the Snowdrift quadrant, and in the limit that same quadrant had carried at least 824 equilibria at every game. But no game of Prisoner’s Dilemma, Stag Hunt or Harmony ever loses its last equilibrium. Over most of Harmony a single equilibrium survives, and it is *ALLC* alone; the counts rise into three figures only in a strip along the *v* axis. The line set stratum shatters: its 564 survivors are scattered over 492 separate lines instead of sharing 44. And the single-game stratum, far from emptying, holds 1907 strategies at 1128 distinct games — 596 of them at the one game (*u, v*) = (9999*/*9998, 1*/*9998), which is where a positive error rate carries the limit’s (1, 0). The maximum number of equilibria per game falls from 22069 to 1024, and the corresponding set moves. In the limit *ɛ →* 0, the richest games are close to the origin. At *ɛ* = 10*^−^*^4^ they are in the far field, filling all but 2*τ* = 0.0115*^◦^* of the Stag Hunt’s ninety.

A positive error rate acts in two ways and at different scales. First, it removes equilibria everywhere, including near the origin. For *ɛ →* 0 every equilibrium is weak, but at fixed *ɛ >* 0 those ties resolve. Then nine of ten strategies that hold a two-dimensional equilibrium region in the limit are equilibria at no game anymore. Second, a positive error rate tilts the boundaries of regions that survive by order *ɛ*, which reshapes the far field, beyond *|*(*u, v*)*| ∼* 1*/ɛ* (**SI** §5).

### Efficient equilibria, and where they exist

A strategy is efficient at a game when a population of it attains *E*_max_ in the limit *ɛ →* 0. Below the line *u* + *v* = 1 a strategy is efficient when it cooperates fully, which is attained by 7639 strategies. Above the line a strategy is efficient when it alternates perfectly, which is done by 3072 strategies. Nothing else reaches the optimum on either side. On the line itself the two coincide, and efficiency then asks only that mutual defection carry no weight in self play, which 14757 strategies achieve (**SI** §5). Below the switch line an efficient equilibrium is therefore a strategy that cooperates fully with itself and that no deviation can beat, which is precisely the *partner* strategy of the literature (12, 20), Akin’s good strategy (21), up to a clause about exact ties in those definitions that can bind only on a set of games of measure zero (**SI** §1). Figure 3**a** below *u*+*v* = 1 is thus the complete map of the partners of binary memory-two: at every game, how many of the 7639 fully cooperating strategies are equilibria there, from 8 in a wedge of Prisoner’s Dilemmas to all of them across the Stag Hunt. Above the line the same definition, with alternation as the optimum, carries the notion to games the classical one does not reach. A *rival* never earns less than its co-player (12, 20); writing *w_CD_*and *w_DC_* for the long-run frequencies of the two asymmetric outcomes, *π*(*σ, τ*) *− π*(*τ, σ*) = (*T − S*)(*w_DC_ − w_CD_*), so rivalry depends on the game only through the sign of *T − S*, the side of the line *u − v* = 1, just as efficiency depends only on the side of *u* + *v* = 1. The *friendly rivals* of Murase and Baek (19), strategies that are efficient and rivals at once, are partners automatically — a co-player’s payoff is at most the pair’s average, which is at most *E*_max_ — and their set is constant on each of the four quarter-planes into which the two lines cut the plane: 8 where mutual cooperation is the optimum and *T > S*, which contains every Prisoner’s Dilemma below the switch line and every donation game; 1519 where mutual cooperation is the optimum and *T < S*; and 80 alternators on either side above the switch line (**SI** §5). The eight are exactly the eight partners of the wedge in which Figure 3**a** attains its minimum: in the hardest Prisoner’s Dilemmas the only partners left are the friendly rivals, *TFT-ATFT* (18) among them, and the two strategies whose equilibrium region is exactly *u* + *v ≤* 1 are the two of the eight that are rivals on both sides of *T* = *S*.

**Figure 3:**
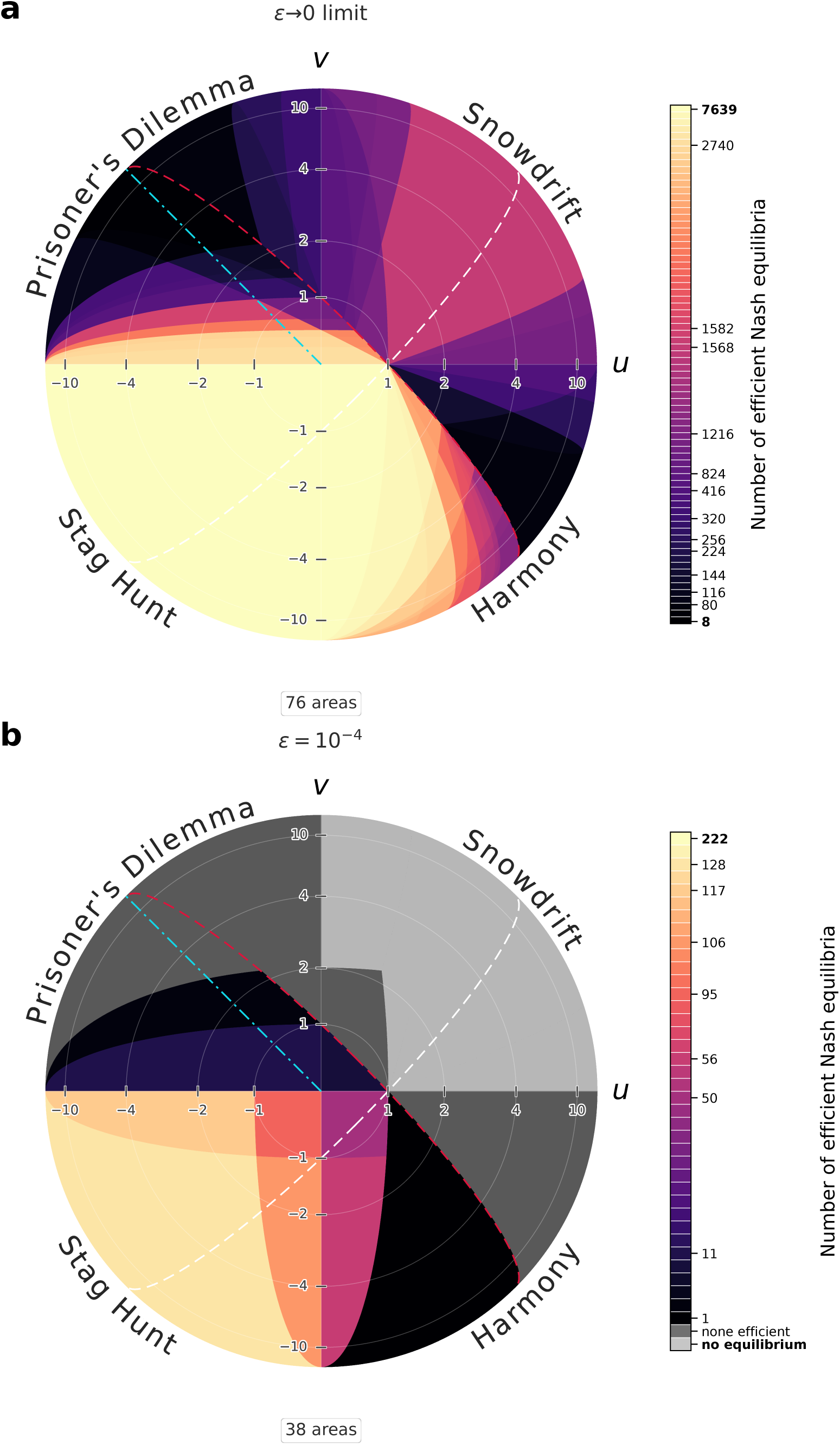
What a positive error rate costs the efficient equilibria. The same disk of games as Figure 2a and the same two notions of equilibrium, now counting at each game only the equilibria that attain the best jointly achievable payoff: **a**, the vanishing-error limit; **b**, *ɛ* = 10*^−^*^4^. Three families of games are marked on both. The crimson line *u* + *v* = 1 is where the social optimum switches from mutual cooperation to alternation; the white line *u − v* = 1 is where the two asymmetric outcomes pay equally, the symmetric anti-coordination game; and the dash-dotted cyan ray is the donation game with benefit 1 and cost *c*, at *u* = *−c/*(1 *− c*) and *v* = *c/*(1 *− c*), the half-line *u* + *v* = 0, *v >* 0 — a half-line and not a line because a cost must be positive. Each panel carries its own colour scale, and **b**’s has two categories off the spectrum: light grey marks the games with no equilibrium of any kind, dark grey those that have equilibria but not one of them efficient. Neither grey occurs in **a**. Where the Prisoner’s Dilemma keeps efficient equilibria at all, the smallest such set at a generic game is four variants of *AON*_2_. The box beneath each disk counts the field’s distinct areas, and the faint white circles are the images of *|*(*u, v*)*|* = 1, 2, 4, 10, crossing the axes at the ticks of those values.

Figure 3 counts for each game the equilibria that are efficient. In the limit *ɛ →* 0, the count runs from 8 to 7639 at a generic game. Everywhere on the map there is an efficient equilibrium. The maximum number of 7639 is reached across the whole Stag Hunt quadrant. The minimum of 8 is reached in a wedge of Prisoner’s Dilemmas below the line *u* + *v* = 1, between *v* = 2 and *u* + 2*v* = 2.

At *ɛ* = 10*^−^*^4^ the abundance of efficient equilibria declines. The count runs from 0 to 245. Above the line *u* + *v* = 1 the efficient equilibria form a set of measure zero: the alternators that survive hold lines and isolated games, but never a region. Below the line, there is a large set of Prisoner’s Dilemmas that have no efficient equilibrium. Below the line, only 245 fully cooperating strategies hold on to a two-dimensional region of games where they are equilibria. The union of those regions is given by the wedge

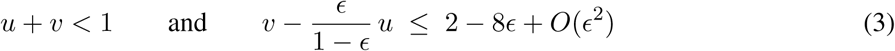

The second inequality derives from a line contributed by *AON*_2_. The smallest count of efficient equilibria in that wedge is one, and that equilibrium is *ALLC*. The corresponding region lies mostly in Harmony, but far out it also gains a sliver of Stag Hunt. The Prisoner’s Dilemma keeps some efficient equilibria near the origin but none in the far field. If it keeps any at a generic game, it keeps at least four. The Stag Hunt has the largest count at any single game, 245, which is a maximum certified over the whole plane, but the games attaining it lie very far out: they claim an unbounded region whose nearest point is at *|u| ≈* 2 *×* 10^7^. The Stag Hunt also has a wedge near the border with Prisoner’s Dilemma where the count drops to zero, which follows from equation (3); that wedge is exactly *τ* wide.

### What fraction of a game’s equilibria is efficient

Figure 4 shows the fraction of Nash equilibria that are efficient. In the limit *ɛ →* 0, we find two regions where all Nash equilibria are efficient, the first of them away from the tie lines on which an inefficient strategy holds its whole region: (i) in Snowdrift outside the rectangle 0 *< u ≤* 1, 0 *< v ≤* 2 and (ii) in Harmony within the wedge *u >* 2*/*3, *u − v >* 1, *u* + *v <* 1 with corners at (2*/*3*, −*1*/*3) and (1, 0). We also note that the fraction of efficient equilibria is nowhere zero. Every game carries an efficient equilibrium, and the worst game manages 8 of 1091. For *ɛ* = 10*^−^*^4^, we still find some areas where all Nash equilibria are efficient. They occur in Harmony below the line *u* + *v* = 1 and to the right of the line 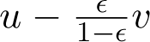 = 2*/*3 — the limit’s *u* = 2*/*3 tilted by *τ*, so that at every *u >* 0 the region is reached once *|v|* is large enough, and past the *v* axis it continues into the Stag Hunt as a wedge of width *τ* — and in two corners of the Snowdrift triangle *u, v >* 0, *u* + *v <* 1, where ten fully cooperating strategies constitute the entire equilibrium set. The fraction of efficient equilibria is zero (meaning there are equilibria, but not one of them is efficient) over Harmony above the line *u* + *v* = 1, over Prisoner’s Dilemma outside a bounded quadrilateral, and in a thin wedge of Stag Hunt along *v* = 0 — in each case away from the measure-zero lines and single games where a surviving alternator or fully cooperating strategy holds no region. The quadrilateral’s far vertex and the wedge’s apex both sit at *|u| ≈* 2 *×* 10^4^, which is 2*/ɛ*.

**Figure 4:**
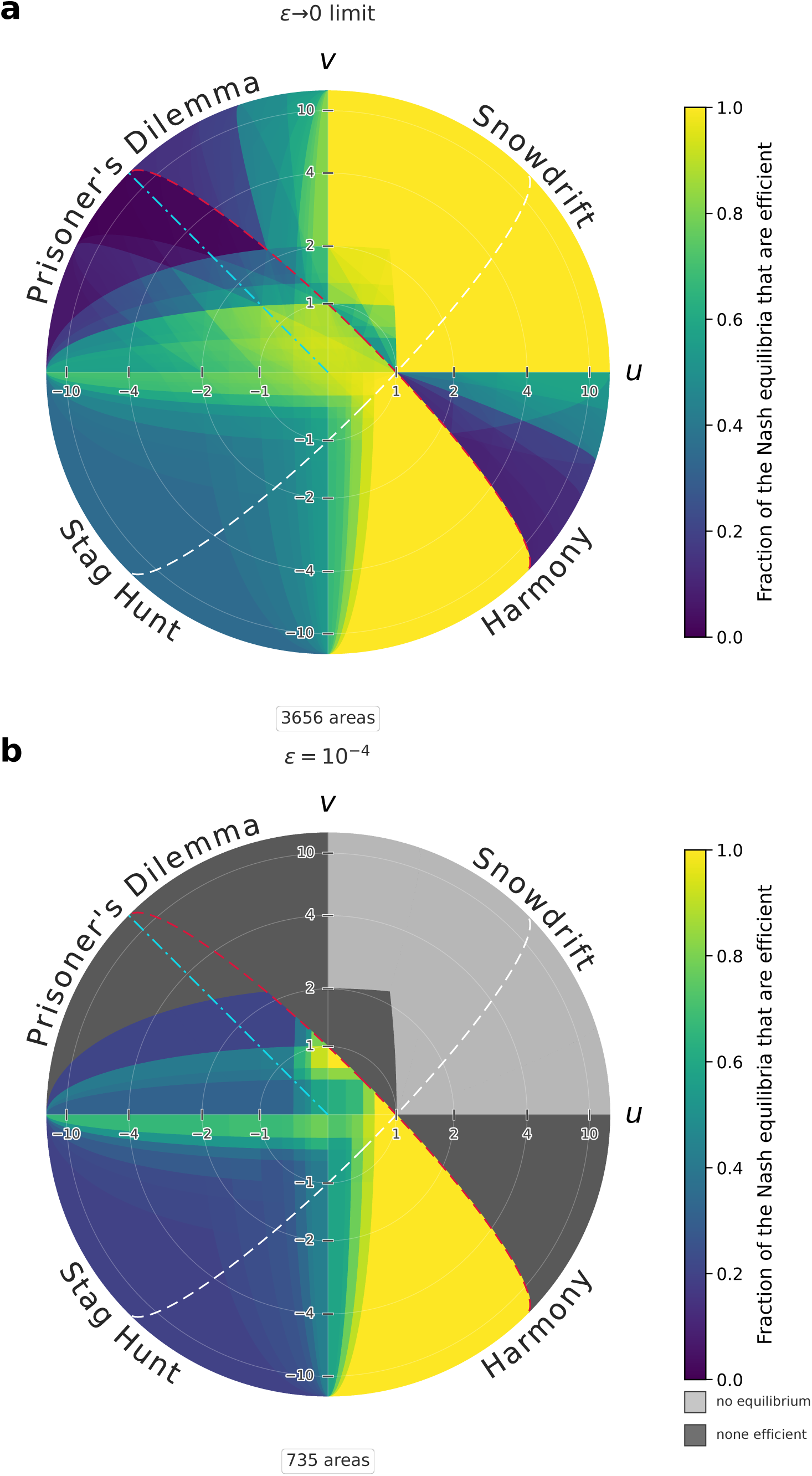
What share of a game’s equilibria is efficient. The same disk and the same two panels as Figure 3 — **a**, the vanishing-error limit; **b**, *ɛ* = 10*^−^*^4^ — now drawing not the count but the *share*: the field of Figure 3 divided, game by game, by the count of all equilibria there, the field of Figure 2a and b. The scale is linear from 0 to 1, and yellow is 1, every equilibrium of that game being efficient. A share is a ratio of two counts rather than a count, so it takes far more distinct levels than the field it divides — 3656 against Figure 3**a**’s 76, the totals in the boxes — and its bar is a continuous ramp where Figure 3’s enumerates. The two off-spectrum categories are Figure 3’s, in the same two greys and covering exactly the same games, so the pages can be compared by colour alone: light grey where a game carries no equilibrium at all and the share is undefined, dark grey where it carries equilibria but not one of them is efficient — an exact zero, and a statement about whole families of games rather than a small number: just over 3*/*8 of the directions at infinity, the largest of the four kinds of game. Both are confined to **b**. Figure 5 is this page read at infinity: its four fills are these four colours — the two greys, the body of the ramp, and the yellow at its top — and it measures each as a fraction of the directions at infinity rather than of the drawn disk.

### Measurement

How much of the plane does each kind of game cover? A share of an unbounded plane is not a number until a window is named. Therefore, we measure the natural density: the fraction of the *directions* at infinity that a set occupies, read at the rim of the disk of (2) (**Methods**). Three angles fix every share, the third asleep at realistic error rates. One is *τ* = arctan[*ɛ/*(1 *− ɛ*)], by which a positive error rate tilts *ALLC*’s and *ALLD*’s equilibrium boundaries off the *u* and *v* axes. The second is *σ*, the width of one of two thin slivers inside the Snowdrift quadrant along which an equilibrium survives arbitrarily far out, held by the same six strategies at every error rate below 1*/*4. The third, *ϱ*, is identically zero for *ɛ ≤* 1*/*4: above that threshold — and only there — four further strategies keep equilibria arbitrarily far out inside the Snowdrift quadrant, along two arcs adjoining the tilted axes. All three are explicit algebraic functions of *ɛ* given in **SI** §6; *σ* becomes two fifths of *τ* as *ɛ* vanishes, and all vanish with *ɛ*.

In the limit, Figure 5**a**, only two of the four kinds occur: every game carries an equilibrium and every game carries an efficient one, so the first two are empty. Of the directions, 5*/*8 carry equilibria of both kinds — the Prisoner’s Dilemma, the Stag Hunt and the half of Harmony above *u* + *v* = 1 — and 3*/*8 carry nothing but efficient ones, the whole Snowdrift quadrant together with the half of Harmony below that line.

**Figure 5:**
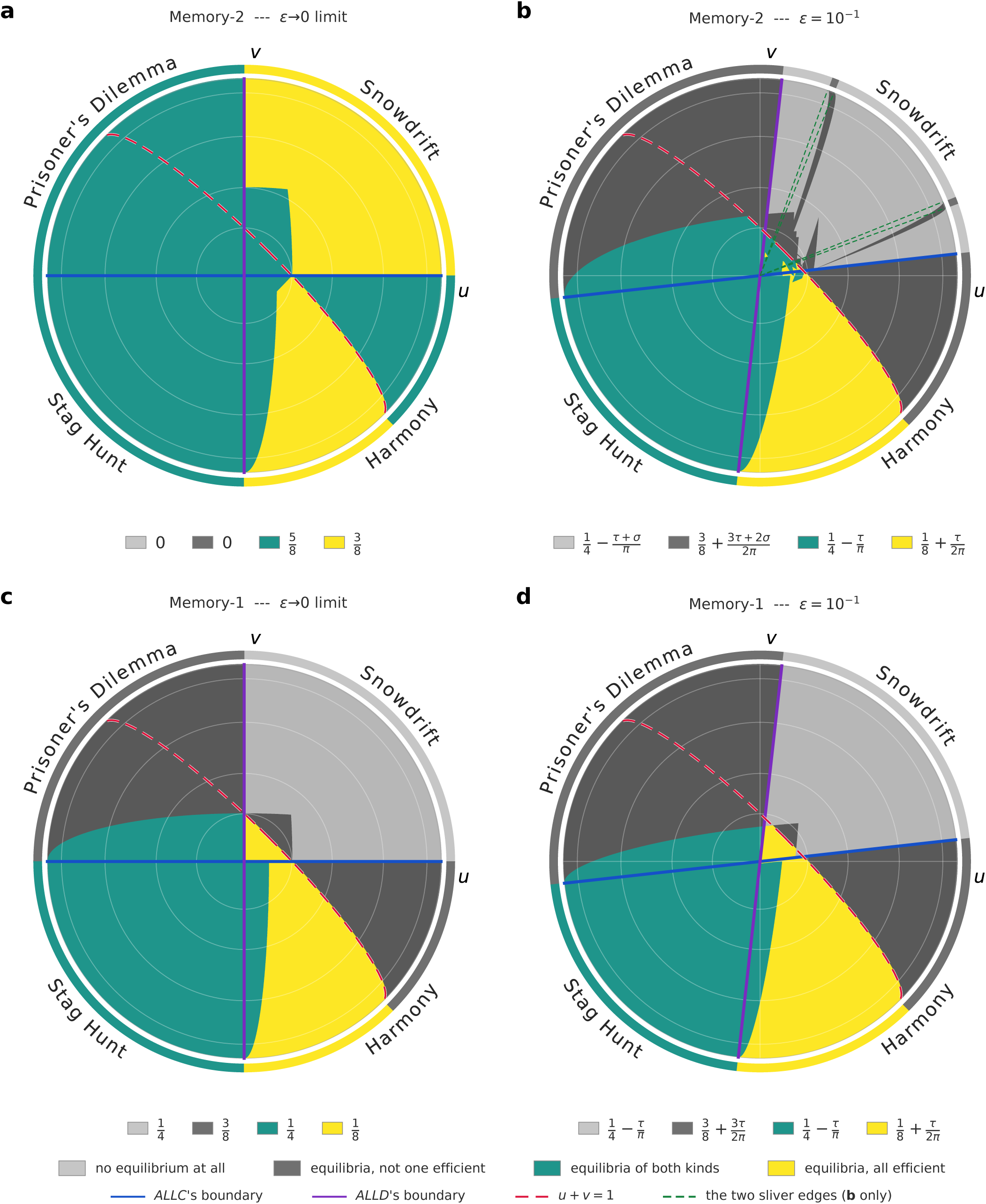
The four kinds of game, and how much of the plane each one is. Every game in the plane is of exactly one of four kinds, and this is the map of them. Each panel draws the disk of (2) at *k* = 4, shaded light grey where the game supports no Nash equilibrium at all, dark grey where it supports equilibria of which not one is maximally efficient, teal where both kinds occur at once, and yellow where every equilibrium is efficient — the two greys of Figures 3 and 4, the body of Figure 4’s ramp, and its fraction-one yellow, so that the three pages share one key. **a** and **b** are the 65536 binary memory-two strategies, **c** and **d** the 16 binary memory-one strategies; the left column is the limit *ɛ →* 0 and the right column *ɛ* = 10*^−^*^1^. Two things are drawn and they are different things. The field inside each rim is what happens at the games the disk reaches, half of which lie within *|*(*u, v*)*|* = 4; the faint white circles are the images of *|*(*u, v*)*|* = 1, 2, 4, 10, as in Figure 3. The ring outside each rim is the exact natural density — the share of the *directions* at infinity that each kind takes — and the four expressions under each disk are the ring’s, one to a colour and in the key’s order, not shares of the drawn area. The three lines are the same three in every panel and each is exact: *ALLC*’s boundary, *ALLD*’s boundary and the switch line *u* + *v* = 1. The first two are the *u* and *v* axes in the limit and tilt off them by *τ* once the error rate is positive; the third is the one that does not pass through the centre, which is why the kinds it bounds have a density and not a share of any window. The two green sliver edges exist only in **b**. Read along a row: memory-one’s four densities depend on *ɛ* through *τ* alone, so **c** and **d** differ in that tilt and in nothing else, while **a** and **b** have almost nothing in common — nearly the whole Snowdrift quadrant passes from all-efficient yellow to no-equilibrium grey, all but the two wedges of width *τ* along the axes and the two interior slivers, which turn dark grey, and the dark grey kind, exactly empty in the limit, takes 0.4388 of the directions. Read down a column: in the limit the extra round of memory buys the whole of Snowdrift, from no equilibrium at all in **c** to nothing but efficient ones in **a**, and it turns the Prisoner’s Dilemma and the half of Harmony above the switch line from inefficient-only into both kinds; at the drawn *ɛ* it buys the two slivers alone, 2*σ* = 3.96*^◦^* of the directions (above *ɛ* = 1*/*4 two further arcs join them; **SI** §6), and what they keep is an equilibrium and never an efficient one. Drawn at *ɛ* = 10*^−^*^1^, where *τ* = 6.34*^◦^* and *σ* = 1.98*^◦^*; at 10*^−^*^4^ they are 0.0057*^◦^* and 0.0023*^◦^*, so at the paper’s own error rate the four numbers under **b** would read 1/4, ⅜, 1/4 and ⅛ to three decimals — memory-one’s limit values, and neither column anything like **a**. **SI** §6 derives every entry and **SI** Table 2 gives all sixteen in closed form.

At every error rate 0 *< ɛ <* 1*/*2, Figure 5**b**, all four kinds occur, and all four are exact:

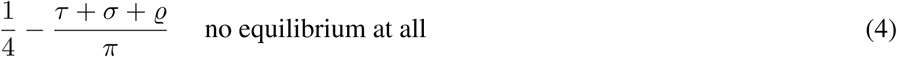

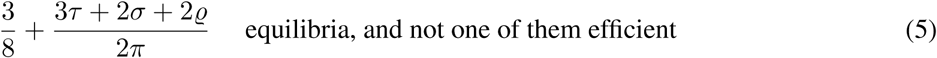

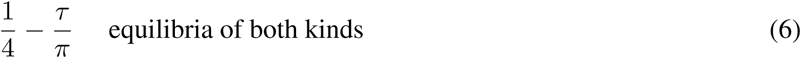

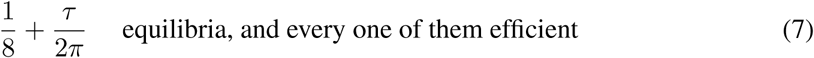

They sum to 1. The first share is the Snowdrift quadrant less a wedge of angle *τ* along each axis, less the two slivers and — above *ɛ* = 1*/*4, where *ϱ* wakes — less the two further arcs — that is the cone of *directions*, and not every game in it, because near the origin Snowdrift games keep equilibria, and at two corners of the triangle *u, v >* 0, *u* + *v <* 1 they keep none but efficient ones. The first two together are the games with no efficient equilibrium, 5*/*8 + *τ/*2*π*, and that share is *ALLC*’s alone: *ALLC* is efficient wherever *u* + *v ≤* 1 and is an equilibrium exactly on (1 *− ɛ*)*v ≤ ɛu*, so its own region is the inner wedge *u* + *v ≤* 1, (1 *− ɛ*)*v ≤ ɛu*, whose cone of directions is that of (3), and nothing else adds an arc of directions to it. Far out, the second kind — the dark grey of Figures 4**b** and 5**b** — is Harmony above the switch line, the whole Prisoner’s Dilemma, a wedge of Stag Hunt of width *τ* along *v* = 0, and Snowdrift’s two edge wedges with its two interior slivers (and, above *ɛ* = 1*/*4, the two *ϱ*-arcs). The third share is the Stag Hunt shaved by *τ* at each end, and the fourth share runs from the antipode of *ALLD*’s boundary to the switch line.

Every one of the four shares is discontinuous at *ɛ* = 0. Two rise from nothing, to 1*/*4 and to 3*/*8, while coexistence falls from 5*/*8 to 1*/*4 and the all-efficient share falls from 3*/*8 to 1*/*8. What a vanishing error rate reports is not what any positive one does. At *ɛ* = 10*^−^*^4^ the four shares stand at 0.2499554, 0.3750605, 0.2499682 and 0.1250159, and the first tends to a quarter as the rate vanishes without ever equalling it. All four are statements about 0 *< ɛ <* 1*/*2, the range in which the strategy space exists at all: at *ɛ* = 1*/*2 every strategy is the same coin toss and every one is an equilibrium of every game, so three of the kinds are empty and the whole plane carries equilibria of both kinds.

### Census

Table 1a collects counts for the limit *ɛ →* 0 and at four error rates from 10*^−^*^4^ to 10*^−^*^1^. It gives the numbers of strategies that are Nash equilibria on a two-dimensional set of games, only along a line and only at a single game. It shows the maximum and minimum numbers of Nash equilibria present at a single game. Table 1b does the same for efficient Nash equilibria. For *ɛ →* 0, the largest number of efficient Nash equilibria, 14757, is reached at the game (*u, v*) = (1, 0), while the largest number at a generic game is only 7639.

### Memory-one, for contrast

The same computations over the 16 binary memory-one strategies show what the extra round of memory provides (20, 21, 23–25) (**SI** §§2, 5). In the limit *ɛ →* 0, eight memory-one strategies hold a two-dimensional region of games, four are equilibria only along a line and three only at a single game, and *TFT* is an equilibrium at no game at all — while at every *ɛ >* 0 the single-game stratum is four, *TFT* holding exactly one game. In Snowdrift memory-one holds nothing but *WSLS*’s unit square at the origin and three lines, two of them running to infinity. Memory-one has no efficient equilibrium anywhere above the line *u* + *v* = 1, at any error rate. Attaining the optimum there requires perfect alternation, and no binary memory-one strategy alternates more than half the time. Measured over the directions of the plane, two of the four kinds of game are memory-two’s *exactly* at every positive error rate — the games carrying equilibria of both kinds, and those carrying only efficient ones. The other two differ by the two interior slivers alone and in opposite senses, so what the extra round of memory buys at infinity is those slivers, moved out of carrying nothing and into carrying nothing efficient. In the limit *ɛ →* 0 memory-one is continuous and memory-two is not. Figure 5 sets out all four for both families, and **SI** Table 2 gives them in closed form (**SI** §6).

## Conclusion

The space of binary memory-two strategies is highly relevant for the evolution of cooperation by direct reciprocity. It is large enough to contain the behaviours that matter and small enough to be analysed in full. In a recent study of evolutionary dynamics, binary memory-two strategies resolve every social dilemma, while binary memory-one strategies fail in many of them (36). The ability to enumerate the strategy space enables us to draw a map over the continuous space of games. Since each deviation rules out a half-plane of games, one computation per strategy settles the whole plane at once.

In the limit of a vanishingly small error rate, *ɛ →* 0, the 2^16^strategies realise 475 patterns of self play carrying 229 cooperation rates. The spectrum of cooperation rates has gaps at both ends: no cooperation rate lies strictly between 0 and 1*/*10 or strictly between 9*/*10 and 1. We find that 23861 strategies are equilibria for a two-dimensional region of games; every game carries between 299 and 22069 equilibria. No region is larger than a half-plane, and one of the half-planes is exactly *u* + *v ≤* 1, held by two fully cooperating strategies. *TFT* is an equilibrium at no game at all. Equilibria and efficient equilibria exist at every game in the plane and 3*/*8 of the plane’s directions carry only efficient equilibria. Below the switch line the efficient equilibria are precisely the partner strategies (12, 20, 21), so the map also lists every binary memory-two partner at every game. Among them are the friendly rivals: eight strategies in every Prisoner’s Dilemma below the switch line. In the hardest of those games they are the only partners that remain.

A positive error rate changes the map. For *ɛ →* 0 every equilibrium is weak. Once those ties resolve, the number of equilibria drops dramatically. For *ɛ* = 10*^−^*^4^, we find that 1050 strategies are equilibria for a two-dimensional region of games; every game carries between 0 and 1024 equilibria. The largest region that any strategy holds is half of all games and belongs to *ALLC* and *ALLD*. Measured over the directions of the plane, four shares are exact for every 0 *< ɛ <* 1*/*2 and they partition it: 1*/*4 *−* (*τ* + *σ* + *ϱ*)*/π* of games keep no equilibrium, 3*/*8 + (3*τ* + 2*σ* + 2*ϱ*)*/*2*π* keep equilibria of which not one is efficient, 1*/*4 *− τ/π* keep both kinds and 1*/*8 + *τ/*2*π* keep none but efficient ones. The quantities *τ*, *σ* and *ϱ* are angles that depend on *ɛ*, the third identically zero up to *ɛ* = 1*/*4 and awake only beyond. Binary memory-one strategies, computed the same way, carry two of the four shares exactly, differ in the other two by the two interior slivers — and, for *ɛ >* 1*/*4, by the two intrusion arcs — in opposite senses, and none of their shares jumps in the limit. The discontinuity belongs to memory-two.

The map is built to be used, above all by evolutionary dynamics. Every evolutionary or learning process on this strategy space moves over the landscape drawn here. The 475 self-play distributions determine the behaviour of every homogeneous population, and the equilibrium polygons say which of those populations resist invasion, and where. The outcome of any such process can therefore be classified, game by game, as an efficient equilibrium, an inefficient one, or no equilibrium at all, and the question of whether selection finds the efficient equilibria among the many becomes a measurement (36). The census is the input that informs finer criteria such as stochastic stability, evolutionary robustness, or the size of basins of attraction. It also locates the ties on which drift decides. The exact data behind every figure are public, and the method — one half-plane per deviation — carries over unchanged to asymmetric games, to discounting and to larger action sets.

## Acknowledgments

I have used several AI assistants — mostly Claude Opus 5.0 and Claude Fable 5.1 (Anthropic) but also GPT-5.6 (OpenAI) — for optimizing and writing code, running and checking computations, and drafting and editing text. I have verified every result reported here and take full responsibility for the content. Support from Harvard University Research Computing and AI resources is gratefully acknowledged.

## Author contributions

M.A.N. conceived the study, performed the analysis, made the figures and wrote the text.

## Code availability

All code was written in C, Fortran and Python. It is available at: https://github.com/martin-mn/MapBinM2, together with the computed output behind every figure and table. The repository’s `exact` folder is the proof record of **SI** §7 — the exact kernel, its independent validators, the certificates, the data and their SHA-256 hashes — and `make test` there re-derives a slice of every kind of result in about ten minutes on a laptop.

## Methods

### Payoffs and games

In each round two individuals independently choose *C* or *D*. The payoff matrix has entries *R* for mutual cooperation, *S* for cooperating against a defector, *T* for defecting against a cooperator and *P* for mutual defection. Any such symmetric game with *R > P* is equivalent, under a positive affine rescaling that leaves strategic structure untouched, to (*R, S, T, P*) = (1*, u,* 1 + *v,* 0), which is (1). The donation game with benefit *b* and cost *c* is (*b − c, −c, b,* 0); dividing through by *R* places it on the half-line *u* + *v* = 0, *v >* 0, with *c/b* = *v/*(1 + *v*). That ray is a single family rather than a generic slice of the plane, and it is the dash-dotted cyan ray of Figure 3.

### Strategies and the error rate

A memory-two strategy is a vector **p** = (*p*_1_*, …, p*_16_), where *p_j_* is the probability of cooperating given the outcomes of the last two rounds; we index states by (most recent outcome, the one before). A binary strategy *intends* either cooperation or defection at each state; the error rate *ɛ* is the probability that the realised action differs from the intended one, independently across rounds and players, so every realised cooperation probability lies in *{ɛ,* 1 *− ɛ}*, a strategy is a 16-bit integer of intended actions, and the space has 2^16^ = 65536 elements. Relabelling *C* and *D* exchanges the four outcomes in pairs, *CC* with *DD* and *CD* with *DC*, so it carries the state indexed *j* to the state indexed 17 *− j* and flips the action taken there: a strategy’s conjugate is its sixteen-bit string reversed and complemented. Two memory-two strategies define a Markov chain on the sixteen states; for *ɛ >* 0 every transition the shift permits is positive, so every state is reached from every other in two rounds and the chain is irreducible, and its unique stationary distribution determines both players’ payoffs.

### Two equilibrium notions

A resident **p** is a Nash equilibrium of a game at a fixed *ɛ* if no strategy earns strictly more against it — equivalently, if in a large population of residents no rare mutant is favoured by selection. At *ɛ >* 0 this needs only the sixteen single-component deviations to be tested (49): against a fixed memory-two opponent, a deviator faces an average-reward Markov decision process on the same sixteen states, every feasible transition is bounded below by *ɛ*^2^ *>* 0 and every state is reached from every other in two rounds, and unimprovability is therefore equivalent to optimality. Since a memory-two continuation depends only on the last two outcomes, and at *ɛ >* 0 every one of the sixteen states carries positive probability, optimality from every state is optimality after every history: within this strategy class the fixed-*ɛ* criterion is subgame perfection, with *ɛ* doing the work Selten’s trembling hand was introduced for (33). In the limit *ɛ →* 0 that reduction fails (50, 51) — a component governing a state that is never visited contributes nothing, the corresponding constraint becomes vacuous, and the test admits strategies it should not — so the limit computation compares every resident against all 65535 rivals. The limit notion is defined by the perturbation: **p** is an equilibrium of the limit at a game exactly when

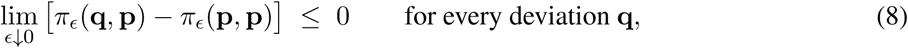

a *perturbation-selected zero-noise Nash relation* in Selten’s sense rather than the Nash set of the unperturbed repeated game, whose *ɛ* = 0 payoffs can depend on the initial history and on the selection among recurrent classes (**SI** §3). The difference between the two notions is not only in the *definition*: the deviation coefficients themselves move by *O*(*ɛ*), so the fixed-*ɛ* set is not contained in the limit set — 382 residents, 162 of them on an open set of games, are equilibria at *ɛ* = 10*^−^*^4^ and at no game at all in the limit: all equilibria of the limit game are weak, and a positive error rate is what decides the ties they leave open. Details, and the numerical treatment of the limit, are in **SI** §§3–4 and §7.

### Shares of the –plane

A share of an unbounded plane is not a number until a window is named, and every window gives a different one. The four shares quoted above are therefore natural densities: the limit, as *r → ∞*, of the share of the disk of radius *r*. Each set of games here is a finite Boolean combination of half-planes, differences included, so that limit exists and equals the fraction of directions at infinity the set occupies, which is what makes all four exact rather than approximate. A density is not the share of anything drawn, and here the two genuinely differ: the games with no equilibrium lie inside a cone with its apex at the origin but do not fill it, since a neighbourhood of the origin inside that cone still carries equilibria. On a bounded window the share therefore reads low and climbs to the density only as the window grows.

### Exactness

Every *ɛ →* 0 quantity in this paper is computed symbolically. The transition probabilities of a pair chain are *ɛ^k^*(1 *− ɛ*)^2^*^−k^*, so by the Markov chain tree theorem each limiting stationary probability is a ratio of integer counts of minimal-error spanning trees; those integers are computed exactly, with a proved bound in place of any tolerance, for all 4.29*×*10^9^pairs (**SI** §7). Self-play distributions, cooperation rates and the limit deviation coefficients are exact rationals by construction — no extrapolation, no floating point and no rational reconstruction enter the argument — and the resulting geometry — region dimensions, facets, areas — is done in exact rational arithmetic with no tolerance and no bounding box. Efficiency is decided exactly: a strategy is efficient at a game when its payoff equals *E*_max_ as an identity between rational numbers. Throughout, a strategy is labelled by its exact *ɛ →* 0 pattern of play even when equilibrium membership is tested at *ɛ >* 0, because at a positive error rate nothing sits exactly on a corner of the simplex and any thresh-old would silently do the classifying. The limit fields of Figures 2 to 4 are drawn from the exact polygons of the arrangement rather than sampled, and the fixed-*ɛ* fields of Figures 2 to 4 are rasters evaluated exactly at pixel centres. All four of Figure 5’s fields are rasters, at 6001^2^, in double precision, which the width of its narrowest feature — *σ*, which is 1.98*^◦^* at the *ɛ* = 10*^−^*^1^ of its right-hand column — makes ample; every line, every ring and every number drawn over them is exact. The drawn maxima of Figure 2**b** and Figure 3**b** are therefore 968 and 222, short of the certified whole-plane values 1024 and 245, which are attained only in the far field; Figure 2**a**’s maximum of 22069 is attained at the corner (*u, v*) = (*−*4*, −*2) of its region and is drawn exactly.

