## Supplementary Information for "An exhaustive map of binary memory-two direct reciprocity over symmetric two-player games"

Martin A. Nowak

Department of Mathematics, Department of Organismic and Evolutionary Biology, Harvard University, Cambridge, MA 02138, USA

The primary object of study is the space of binary memory-2 (M2) strategies, which answer each of the 16 two-round histories by cooperation or defection. There are $2^{16} = 65536$ binary M2 strategies. The game is infinitely repeated and the future is not discounted, so a payoff is the long-run average per round, read from the stationary distribution of the Markov chain that two strategies generate. We assume that each decision is implemented subject to a small error rate, $\epsilon$. There are two cases that need to be distinguished: (i) the limit $\epsilon \to 0$ and (ii) a fixed positive error rate $\epsilon > 0$.

First we study the distribution over the four outcomes of a single round, $CC, CD, DC, DD$, that each binary M2 strategy achieves in self play. This calculation is independent of any payoff matrix. For the limit $\epsilon \to 0$, we find 475 possible one move distributions and 229 distinct cooperation rates. The spectrum of cooperation rates has large gaps between 0 and $1/10$ and between $9/10$ and 1. For $\epsilon = 10^{-2}$, we find 14300 possible one move distributions and 14241 distinct cooperation rates; the large gaps vanish.

Next we project the finite strategy space unto the uncountable infinity of all symmetric two-person games. For each of the 65536 strategies we compute the set of games at which it is a Nash equilibrium (NE). For the limit $\epsilon \to 0$, we find that 23861 strategies are NE for a 2-dimensional set of games, 3774 strategies are NE along a line, 4875 strategies are NE for a single game and 33026 strategies are not a NE at any game. For $\epsilon = 10^{-2}$, we find that 1050 strategies are NE for a 2-dimensional set of games, 564 strategies are NE along a line, 1907 strategies are NE for a single game and 62015 strategies are not a NE at any game. We obtain exactly the same numbers for $\epsilon = 10^{-3}$ and $\epsilon = 10^{-4}$ and very similar numbers for $\epsilon = 10^{-1}$.

The above computation also answer the question: for any particular game what are the binary M2 strategies that are NE. For the limit $\epsilon \to 0$, the number of NE at a specific game varies from 299 to 22069. For $\epsilon = 10^{-2}$, the number of NE at a specific game varies from 0 to 1024.

Next we ask how many NE are efficient, in the sense of reaching the maximum payoff in self play that a game allows. For the limit $\epsilon \to 0$, we find that 9431 strategies are efficient NE for a 2-dimensional set of games, 2737 strategies are efficient NE along a line, 2589 strategies are efficient NE for a single game and 50779 strategies are not efficient NE at any game. The number of efficient NE at a specific game varies from 8 to 14757. For $\epsilon = 10^{-2}$, we find that 245 strategies are efficient NE for a 2-dimensional set of games, 154 strategies are efficient NE along a line, 390 strategies are efficient NE for a single game and 64747 strategies are not efficient NE at any game. The number of efficient NE at a specific game varies from 0 to 245.

These computations enable us to draw maps on the space of games indicating the regions that carry a certain number of NE, or efficient NE, or the ratio of efficient NE over all NE at a game (See Main Text Figures 2-4). For the limit $\epsilon \to 0$, we find that NE and efficient NE are available everywhere and there are large regions where all NE are efficient. For any $\epsilon > 0$, we find regions that have no NE and regions that have no efficient NE; we still find a region where all NE are efficient. We measure all those regions and give precise expressions (see also Main Text Figure 5).

We compare all our findings to binary memory-1 (M1) strategies. There are only 16 binary M1 strategies and they have been studied exhaustively. We find interesting similarities and differences. A curiosity is that the measurements of the previous paragraph are discontinuous in the limit $\epsilon \to 0$ for M2, but continuous for M1.

This SI is in seven parts. Section 1 reviews the related literature and states which of the ingredients and conclusions are known. Section 2 constructs the ladder of achievable self-play cooperation levels, for M1 where it is already in print and for M2 where it is not. Section 3 shows that the Nash region of a strategy is a convex polyhedron in the plane of games, that all 65535 deviations rather than sixteen must be tested in the limit, and how the limit is taken exactly. Section 4 shows that every equilibrium of the $\epsilon \to 0$ limit game is weak — at a fixed $\epsilon > 0$ they are not, and that asymmetry is the point — and draws out the methodological consequence. Section 5 gives the census of regions, the efficiency results and the tie set $\Lambda$. Section 6 divides the directions of the plane into the four kinds of game — no equilibrium, equilibria of which not one is efficient, equilibria of both kinds, and equilibria all of which are efficient — and measures each of them exactly. Section 7 gives the exact methods — the tree-theorem kernel, the complete constraint sets, the all-rate certification — and the verification record, every part of which is now proved rather than measured.

This supplement carries six figures, referred to as SI Figure 1 to SI Figure 6. Four of the results below are drawn in the main text rather than here: its Figure 2 gives the Nash set of the plane stratified by dimension — in the $\epsilon \to 0$ limit on the left, at $\epsilon = 10^{-4}$ on the right — its Figure 3 the number of maximally efficient equilibria in the two notions of equilibrium, one above the other, its Figure 4 the fraction of a game's equilibria that are efficient, and its Figure 5 the four kinds of game of §6 on four disks — memory-2 above, memory-1 below, the $\epsilon \to 0$ limit on the left and $\epsilon = 10^{-1}$ on the right — which is Table 2 of this supplement drawn, in that table's own four colours. SI Figure 6 draws the three sets themselves, one to a panel, with the lines that fix them — and, in a fourth panel, the first set above the threshold $\epsilon = \frac{1}{4}$, where the intrusion arcs of §6 are open — and §6 derives every entry. Where a statement below is about the arrangement of the plane rather than about the ladder, those are the figures it refers to.

### 1 Related literature

**What this section marks, and how.** In the following we discuss the literature on which this work builds, and mark explicitly where a statement made later in this SI is a rediscovery of something already published, where it sharpens a known result, and where it is new.

#### 1.1 Direct reciprocity, and the whole space of two-player games

**Where direct reciprocity is usually studied.** Direct reciprocity is a mechanism for the evolution of cooperation (1–5). It is usually studied in the repeated Prisoner's Dilemma (PD) or in its special case the donation game, but extensions to other social dilemmas have been considered, including the Snowdrift game (6) and arbitrary repeated $2\times2$ games (7–9). According to the Folk theorem, repeated games admit a wide range of equilibria (10, 11), which is the reason why an equilibrium analysis alone rarely settles what a population does.

**Two parameters suffice.** Since positive affine transformations of the payoffs change neither the best replies nor the efficiency ordering, the space of symmetric $2\times2$ games is two-dimensional. That reduction, and the fact that the relevant equilibrium conditions depend on the payoffs only through two combinations, is argued in detail by Wang et al (12), while Ito and Tanimoto (13) organise the five reciprocity mechanisms on the resulting plane. Phase diagrams over the whole plane, with the four classical families in the four quadrants, are by now a standard instrument (14–17). The particular normalisation used here, $(R, S, T, P) = (1, u, 1 + v, 0)$, is the one of LaPorte et al (18), who use it to define a two-dimensional space of games in which the four classical social dilemmas are the four quadrants of the plane.

**Direct reciprocity over the whole plane.** Martinez-Vaquero et al (7) study the 16 deterministic memory-1 strategies, perturbed by an implementation error, over the entire space of symmetric $2\times2$ games. They determine which strategies and which mixed populations are stable in each part of the plane. This is the memory-1 ancestor of the present computation. Their coverage, however, is by *representative points per region* rather than by exact construction, and that is precisely the limitation Section 5 addresses: the equilibria that live on measure-zero sets of games cannot be found by any sampling of games.

**Equalizers across the social dilemmas.** Hübner et al (9) generalise the theory of zero-determinant strategies from the donation game to all pairwise social dilemmas, which they define by $\min\{R, T\} > \max\{S, P\}$. They construct, for every game with $T > R$, equalizer strategies that sustain mutual cooperation as an equilibrium by direct reciprocity, by indirect reciprocity, or by a mixture of the two. For $T < R$, where mutual cooperation is already a one-shot equilibrium, they show that equalizers provably cannot enforce it and *ALLC* is used instead. Their result is the memory-1 version of the existence half of Section 5, and it must be read alongside it, but two differences matter. First, they identify the socially optimal outcome with mutual cooperation. Their class of games includes games with $T + S > 2R$, in which two players who alternate between the two asymmetric outcomes obtain $(T + S)/2 > R$. In our notation these are the games with $u + v > 1$, and there mutual cooperation is not payoff maximising. The efficiency normalisation used in Section 5, $E_{\max} = \max\{R, (S + T)/2\}$, follows the attainable optimum across that line, so on one side of $u+v = 1$ our notion of "maximally efficient" agrees with theirs and on the other side it does not. Second, their result is an existence proof: it shows that suitable strategies exist. The question of *how many* equilibria there are and what fraction of them are efficient — which is what Section 5 answers — is not asked there.

**The closest predecessor, and what separates it.** LaPorte et al (18) are the natural point of comparison for Section 3. Working with memory-1 strategies, they give explicit analytic Nash conditions for five classes of equilibria, cooperators, defectors, trans-alternators, cis-alternators and CD-repeaters. They prove that among memory-1 strategy profiles of those types there are no further equilibria. They plot, as a function of the discount factor, the precise set of games for which a (LF-stable) Nash equilibrium type can appear. Their conditions are linear in the payoffs, so their regions are intersections of half-planes. Three things distinguish their work from the present computation. Their analysis is for memory-1 and by strategy class. Ours is for memory-2 and by individual strategy, over the complete space of 65536 residents and 65535 deviators. They do not state the convex-polyhedron result of Section 3 as such, they do not compute the resulting arrangement of the plane, and they do not treat the degenerate lines that are the subject of Section 5. They do, however, already contain a worked memory-2 Nash equilibrium in a Snowdrift game, and a proof that a memory-1 characterisation does not always extend to equilibrium strategies with longer memory, which is one motivation for the present work.

#### 1.2 Deterministic strategies with implementation error, and exhaustive enumeration

**The strategy space is not new.** A memory-$m$ strategy that responds deterministically to the last $m$ rounds, subject to a small probability of mistaken implementation, is the strategy space of Lindgren (19), a binary genome of $2^{2m}$ bits; for $m = 2$ the genome has 16 bits and the space has $2^{16} = 65536$ members, which is exactly the space studied here. The encoding of a deterministic memory-$m$ strategy as a bit string goes back to Axelrod's genetic algorithm (20), whose 64 genes are the responses to the three-round histories; what Lindgren adds is the implementation mistakes and a variable memory length. Hauert and Schuster (21) examined numerically how outcomes change as memory is extended beyond one round.

**Noise, and why the probabilities are held at $\epsilon$ and $1-\epsilon$.** That an implementation error changes the equilibrium analysis qualitatively, by making every history reachable and hence every deviation payoff-relevant, is the point of Boyd (22): with mistakes a pure strategy can be evolutionarily stable, provided it is a best reply to itself after every history. The parameterisation used here is that of Nowak and Sigmund (23), who take the 16 deterministic memory-1 rules — numbered 0 to 15 by the binary expression of the index, which is the coding used throughout this SI — and take uncertainty into account by replacing 1 by $1-\epsilon$ and 0 by $\epsilon$ in the quadruples, with $\epsilon$ the frequency of errors. Everything below is the memory-2 analogue of that construction: the two-point set $\{\epsilon, 1-\epsilon\}$, not an interval, and the noise floor read as an error rate rather than as a strategy parameter. They note that at $\epsilon > 0$ the first round no longer matters, which is why the limiting distribution is well defined without an initial condition, and they already connect the device to Selten's trembling hand. Lorberbaum et al (24) later supply the justification for restricting to those two points: allowing the memory-1 probabilities to range over the whole closed interval $[\epsilon, 1-\epsilon]$, they show that the search for evolutionarily stable strategies reduces to the *boundary* strategies whose probabilities are exactly $\epsilon$ or $1-\epsilon$, and exhibit three genuine evolutionarily stable strategies there. Section 4 must therefore be read carefully: our statement that no binary memory-2 strategy is a *strict* equilibrium is a statement about the $\epsilon \to 0$ limit game, and it is consistent with both of those papers, because at fixed $\epsilon > 0$ the deviation deficits we measure are positive and of order $\epsilon$.

**The $2^{16}$ space has been enumerated before.** Hilbe et al (25) state that "there are $2^{16} = 65{,}536$ pure memory-2 strategies" and perform an exhaustive analysis of all of them at a fixed error rate $\epsilon = 0.01$, asking for each resident whether any other pure strategy can obtain at least the same payoff — that is, testing all deviations rather than local ones. In the donation game they classify the resulting equilibria into four groups: eleven that yield the mutual-cooperation payoff (four equivalent to $AON_2$, four to $AON_1$, three delayed $AON_1$), fifteen at mutual defection including *ALLD* and *Grim*, eight self-alternating strategies, and a class with two absorbing states. They also prove that $AON_n$ is a subgame-perfect equilibrium if $b/c \geq (n+1)/n$, so that memory-1 requires $b/c \geq 2$ and memory-2 only $b/c \geq 3/2$. Two consequences for the present work should be stated plainly. First, the memory-2 cost threshold is theirs, not ours: the facet $T \leq 3R$ of Section 5, contributed by the *ALLD* deviation, intersects the donation line $v = -u$ at $c/b = 2/3$, which is exactly their $n = 2$ threshold; we recover it rather than establish it. Second, they call their memory-2 equilibria strict Nash equilibria of the game at $\epsilon = 0.01$, which is the correct statement at fixed $\epsilon$ and does not contradict Section 4. Their treatment of "consistent" $n$-histories — imposing the equilibrium conditions only on those histories that are revisited with positive probability, on the grounds that in the limit of rare errors a strategy may never experience certain histories at all — is moreover the closest published statement of the mechanism behind the second half of Section 3, though it appears there as a definitional restriction rather than as a demonstrated failure of a test.

**The $\epsilon \to 0$ limit is likewise established practice.** Murase and Baek (26) enumerate the deterministic strategy spaces exhaustively at memory-1 (16 strategies), memory-2 (65536) and memory-3 (about $1.8 \times 10^{19}$, on a supercomputer) under an implementation error, with efficiency defined as a property of the limit $\epsilon \to 0$; they show that efficiency and defensibility are jointly unattainable at memory-1, that among deterministic memory-2 strategies only four — the TFT-ATFT family of Yi et al (27) — satisfy all three criteria, and that memory-3 admits some $4 \times 10^{12}$ such strategies with strictly shorter error-recovery paths, CAPRI among them. See also (28, 29) for the automaton representation of such strategies and for the exhaustive treatment of all memory pairs with $m_1 + m_2 \leq 4$, and Yi et al (27) for an explicit deterministic memory-2 strategy that is efficient, defensible and a cooperative Nash equilibrium under vanishing noise. A full sweep of the $2^{16}$ space combined with the $\epsilon \to 0$ limit is therefore not the novelty here; the novelty is what is computed with it. Deterministic memory-2 strategy pairs have been studied analytically by Ueda (30), who gives a necessary condition for them to form symmetric mutual-reinforcement-learning equilibria — a notion he distinguishes from Nash equilibrium — together with three examples. Memory-two zero-determinant strategies, which are stochastic and are not analysed there as equilibria, were constructed by Ueda (31); they enforce *linear* relations among correlation functions of the two players' payoffs in successive rounds, the memory-one case of which is a linear relation between the average payoffs. That last result is the published mechanism behind the exactly antiparallel constraint pairs of Section 5.

**The self-play spectrum, which §2 constructs, is published at memory-1.** The $\epsilon \to 0$ one-move self-play distributions of the 16 binary memory-1 strategies are the diagonal of Table 1 of Nowak, Sigmund and El-Sedy (32), which tabulates the whole $16 \times 16$ matrix of limiting distributions for this construction; Table 2 of Kim, Choi and Baek (33) gives the same sixteen self-play distributions in leading-order $\epsilon$ form. The first of those also proves a finiteness statement wider than self play — across all 256 memory-1 matchups the limiting distributions take only eight forms up to permutation — and identifies the monomorphic optima. So the memory-1 half of §2 is a rereading of published tables, not a computation, and we present it as such. What is not in those papers is the reading: no count of the distinct points, no cooperation rate, and no statement of the empty band below full cooperation. At memory-2 we have found no published spectrum at all; the one number that is published is 7639, the size of the efficient set at $m = 2$ in Murase and Baek (26), whose efficiency criterion is exactly $\gamma = 1$ in the limit. Hilbe et al (25) ran the identical $2^{16}$ sweep and isolated, among their equilibria, a class with two absorbing states; but that classification is made at fixed $\epsilon$ under a Nash restriction, and the strategies carrying the $9/10$ ceiling of §2 are not equilibria in their setting, so they do not appear in it. No spectrum is reported there. Here $m$ is the memory length, so $m = 2$ is this paper's space of $2^{16}$ strategies, and $\gamma$ is the self-play cooperation rate, defined at (11).

**Longer memory.** That longer memory facilitates cooperation, and that it does so by making new equilibria available, is a recurring finding (21, 25–27, 29, 34–37). Memory-3 has $2^{64} \approx 1.8 \times 10^{19}$ strategies (25), and criterion-based sweeps of that space have been carried out on a supercomputer (26); but such a sweep costs $O(N)$, whereas the pairwise Nash census performed here costs $O(N^2)$, which is why memory-2 is probably the last case in which a complete census of this kind is possible at all.

#### 1.3 Which deviations have to be tested

**The best reply against a memory-$n$ resident.** Against a fixed memory-$n$ opponent the deviator faces a finite Markov decision process on the joint histories, so a best reply exists within memory-$n$ and can be computed. Levínský et al (38) prove the existence half, under the discounted criterion, for factored strategies with a recursive factor, of which memory-$n$ is the leading case; Lesigang et al (39) and LaPorte et al (40) develop the consequences, the latter formalising best-reply-completeness and payoff-completeness and showing that bounded memory has both. In the computer-science literature the same reduction appears in Chen et al (41), who represent the resident by a transition graph on $n^{2k}$ history vertices and identify the optimum with a maximum-mean cycle, and in Zhu and Lin (42), who build the history-augmented MDP explicitly. For reciprocity specifically, Glynatsi et al (37) prove that verifying the Nash property of a reactive-$n$ strategy needs only the finitely many deterministic self-reactive-$n$ deviations. Related machinery for computing best replies against memory-1 opponents is in (43–46). The average-reward dynamic programming behind the single-state test — unimprovability implies optimality under a unichain assumption — is standard (47).

**Why the reduction fails in the limit.** At $\epsilon > 0$ every transition of our pair chain that the shift permits is at least $\epsilon^2$ — the state is the last two outcomes, so each of the sixteen has exactly four successors and the other 192 entries are identically zero — and every state is reached from every other in two rounds, so the chain is irreducible under every policy, and the sixteen one-component deviations suffice. As $\epsilon \to 0$ the chain becomes multichain, and it is a classical fact that gain-unimprovability then no longer certifies optimality and that a nested hierarchy is required (48). The precise object is the singularly perturbed limiting-average Markov control problem, for which Bielecki and Filar (49) and Abbad et al (50) show that the $\epsilon = 0$ problem is *not* the limit of the $\epsilon > 0$ problems and give algorithms for limit-optimal policies. The phenomenon that a reduction which is exact for every $\epsilon > 0$ can fail non-uniformly as $\epsilon \to 0$ has close analogues in evolutionary game theory (51, 52), and the general question of when such limits may be interchanged is treated by Sandholm (53). Within direct reciprocity the gap is acknowledged but left open: Glynatsi et al (37) note that with vanishing errors *almost all* of their partner strategies remain only *approximate* Nash equilibria, and state explicitly that the characterisation of partners at positive error rates is not settled. Section 3 therefore does not use the reduction at all: every limit result below is computed from all 65535 deviations of every one of the 65536 residents, the fixed-$\epsilon$ results from the sixteen. The observation that the sixteen-deviation test silently accepts residents in the limit is, as far as we are aware, not stated in the reciprocity literature.

##### 1.4 Weak equilibria: why nothing here is strict

**The classical result.** That equilibria of the repeated Prisoner's Dilemma are weak rather than strict is one of the oldest results in the field. Boyd and Lorberbaum (54) show that for any pure strategy there is a mutant that earns exactly the same payoff against the resident; Farrell and Ware (55) extend the no-ESS conclusion to finite mixtures of pure strategies, and Lorberbaum (56) completes it for completely probabilistic strategies, there by a single invading deviant rather than by a neutral tie. van Veelen (57) builds the apparatus that the weakness makes necessary, defining robustness against indirect invasions and proving it equivalent to membership of a minimal evolutionarily stable set; García and van Veelen (58) prove that in repeated games with sufficiently high continuation probability there is no evolutionarily stable strategy but very many neutrally stable ones, with a stepping-stone path leading out of each, and García and van Veelen (3) state the conclusion in its bluntest form, that no strategy can win. See also (59) for the effect of complexity costs on the stability ranking and (60–62) for the graded stability notions that were introduced because strictness is unavailable. Section 4 is the exact, exhaustive form of this statement inside one finite strategy space and over a whole plane of games; the statement itself is not new.

**Two mechanisms that produce exact ties.** The first is off-path indifference, which is Selten's original motivation for perturbing a game so that every information set is reached (63); our construction — probabilities held at $\epsilon$ and $1 - \epsilon$ so that all sixteen histories occur, with the limit taken afterwards — is precisely that programme, and Selten's warning that the limit set differs from the unperturbed equilibrium set is the conceptual parent of Sections 3 and 4. The connection between the trembling hand and an error rate in the repeated Prisoner's Dilemma is drawn already by Nowak and Sigmund (23). The second is payoff control: an equalizer strategy makes the co-player's payoff independent of the co-player's strategy, so that *every* deviation ties exactly. Equalizers were constructed by Boerlijst et al (64) and subsumed into the zero-determinant strategies of Press and Dyson (43), generalised to arbitrary action spaces by McAvoy and Hauert (65) and to memory-two by Ueda (31); Adami and Hintze (66) observe that such strategies are at most *weakly* dominant. In economics the corresponding notion is the belief-free equilibrium, defined by requiring each player's continuation strategy to be optimal after every private history independently of his belief about the co-player's history, which in the standard constructions forces exact indifference (67, 68); see (69) for the weakly belief-free relaxation. Finally, Dutta and Siconolfi (70) characterise, for one-period-memory strategies, exactly which stage games admit an equilibrium in which both players are indifferent after every history — a small-scale precursor of the region $Z(p)$ of Section 3.

**Where strictness does survive, and why that is not a contradiction.** Alongside Boyd (22) and Lorberbaum et al (24), discussed above, Mailath and Olszewski (71) obtain a folk theorem with bounded-recall strategies whose equilibria are essentially strict. All of these are statements at a positive noise level or in a discounted game. The claim of Section 4 is confined to the $\epsilon \to 0$ limit of the undiscounted game, where the binding constraints hold with equality; at $\epsilon > 0$ the same computation gives strictly negative best-deviation gains, of order $\epsilon$, at every game interior to a resident's region. The methodological corollary — that because every binding constraint of the limit holds with equality, no uncalibrated floating-point sign test can decide equilibrium membership at all — appears to be new, and it is the reason every tie in this paper is decided by the exact integer computation of Section 7.

##### 1.5 Efficiency at equilibrium

**Partners, and the cost thresholds.** A strategy for which full cooperation is an equilibrium — so that the co-player maximises its own payoff by cooperating — is a *partner* (45, 72). The memory-1 partners are characterised by Akin (44) and Hilbe et al (45), the reactive-$n$ partners by Glynatsi et al (37), and the memory-$n$ cost threshold $b/c \geq (n+1)/n$ by Hilbe et al (25). Adaptive dynamics in the memory-1 space is studied by LaPorte et al (73). Everything in Section 5 that concerns full cooperation is therefore a rediscovery of a known threshold in a new coordinate system: what is added is the *shape* of the region in game space, and the identification of its two facets with two specific deviations.

**Does bounded memory cost efficiency?** This is a settled debate in economics that the reciprocity literature rarely engages. Cole and Kocherlakota (74) show that with imperfect monitoring the memory-$K$ equilibrium payoff set can collapse to the single point of mutual defection, and that the $K \to \infty$ limit need not recover the unbounded-recall set; Hörner and Olszewski (75) prove a folk theorem with finite memory and show that the collapse is an artefact of an additional restriction; Barlo et al (76, 77) give bounded-memory folk theorems. Section 5 lands squarely in this debate, on the side of attainability, but in a much more restrictive setting than any of these papers: sixteen bits, no discounting, and a fixed noise floor.

**Existence versus selection.** That the Folk theorem's multiplicity of equilibria makes equilibrium existence uninformative, and that the selection among them is a question for the evolutionary dynamics, is a standard position, stated in almost exactly the words of Section 5 by Tkadlec et al (78), who write that the Folk theorem "guarantees the existence of a multitude of equilibria" and that which one is chosen is therefore a question of evolutionary dynamics; it is stated by Fudenberg and Maskin (79) and argued at length by García and van Veelen (3, 58). The contribution of Section 5 is not that position but a measurement of it: over the whole plane, exactly how many equilibria there are and exactly what fraction of them are sub-maximal. That quantitative census has, as far as we are aware, no published counterpart at any memory length.

##### 1.6 Alternation as the social optimum, and turn-taking

**The dichotomy is known and named.** In games with $2R < S + T$ the payoff-maximising outcome is not mutual cooperation but an alternation in which the two players collect $S$ and $T$ in turn. This is called ST-reciprocity, or alternating reciprocity, and it has been mapped over the whole plane of $2 \times 2$ games by Tanimoto and co-workers (80–82), in the first two cases explicitly with two-length memory strategies. The efficiency normalisation of Section 5, which switches the reference optimum at $u + v = 1$, is that dichotomy written in our coordinates, and we claim no novelty for it.

**The line $S = T$ has been noticed before, and in the same strategy space.** Wang et al (83) study turn-taking with $k$-step memory over an extended $S$–$T$ plane and report that with two-step memory a turn-taking pattern emerges under $S + T > 2R$ *and* $S \neq T$ — most of Snowdrift, all of Leader and Hero, and part of the Prisoner's Dilemma and of harmony — whereas on $S + T > 2R$ with $S = T$ a different, non-alternating strategy takes over. The overlap with the present work is closer than that summary makes it sound, and it is worth stating exactly. Their two-step memory is coded by the same four bits of history as ours and therefore ranges over the same $2^{16}$ deterministic strategies; their reduced payoff matrix $\left(\begin{smallmatrix} 1 & S \\ T & 0 \end{smallmatrix}\right)$ is the normalisation of (13), so their $S + T = 2R$ is our $u + v = 1$ and their $S = T$ our $u - v = 1$. Both of the lines this document is organised around are therefore already visible in their figures. What differs is the rest of the apparatus. They carry no implementation error, and what they report is the attractor of a genetic algorithm sampled at 225 points of the plane, not a statement about equilibria — so the finding comes without a mechanism, and its apparent orientation is the opposite of ours: alternation *retreats* from $S = T$ there, where we find it pinned exactly onto that line. Section 5 should be read as supplying the equilibrium-theoretic reason for both. What we find is that on $u - v = 1$ neither role of an alternation pays more than the other, so that no deviator can profit by seizing one of them.

**Turn-taking in repeated games.** That efficiency in a symmetric repeated game may require asymmetric within-round outcomes reached by alternation is the subject of Bhaskar (84) — whose Proposition 4 is refuted by Kuzmics and Rogers (85), who give counterexamples and show that it is not correctable — and of Lau and Mui (86, 87), whose turn-taking strategies must solve two problems that are exactly the ones we meet: who takes the favourable role first, and how a deviation from the turn-taking path is deterred. Kuzmics et al (88) characterise the symmetric equilibrium payoffs of repeated allocation games and single out alternation as the focal Pareto-optimal symmetric equilibrium, with experimental support. On the evolutionary side, coordinated alternating reciprocity evolves in simulations of games with asymmetric efficient outcomes (89, 90), in a generalised hawk–dove model with evolving updating rules (91), and in human route-choice experiments (92). In the memory-1 direct-reciprocity literature the alternating strategies appear as the trans-alternators of LaPorte et al (18), and Stewart and Plotkin (8) show that an evolutionarily robust memory-1 strategy must be of one of three types, self-cooperating, self-defecting or self-alternating.

**One terminological warning.** The alternation studied here is an outcome distribution $(0, \frac{1}{2}, \frac{1}{2}, 0)$ of a game in which both players move *simultaneously*. It should not be confused with the alternating-move protocol, in which the players take turns to act. That is a different model, introduced by Nowak and Sigmund (93), with its own literature (94–96).

##### 1.7 The geometry of the space of games

**The polyhedron is a known theorem.** That the set of games at which a fixed strategy profile is an equilibrium is cut out by one linear inequality per deviation, and is therefore a convex polyhedron, is not a new mathematical fact; Section 3 is a specialisation of it. In its abstract form it belongs to the study of the equilibrium correspondence over payoff space: Kohlberg and Mertens (97) prove that the graph of the Nash correspondence over the space of games is homeomorphic to the space of games itself, Harsanyi (98) that the degenerate games form a closed set of measure zero, and Levy (99) studies slices of that manifold. In inverse reinforcement learning the same object is the *feasible reward set*: Ng and Russell (100) give the set of reward functions making a policy optimal as a system of linear inequalities, and observe that the characterisation is degenerate; see also Metelli et al (101). In inverse game theory it is the *inverse Nash set*: Wu et al (102) state that the set of games with a given joint policy as its unique Nash equilibrium is a polytope, and Kuleshov and Schrijvers (103) and Bertsimas et al (104) recover games or utilities from observed equilibrium play. Within repeated games, Baklanov (105) characterises all Nash equilibria among reactive strategies *and* determines all symmetric stage games admitting each one, which is the inverse map of Section 3 one memory level down, and Lesigang et al (106) reduce the symmetric reactive equilibrium conditions, for the class of repeated *additive* games, to a finite system of linear equalities and inequalities in the strategy parameters.

**Arrangements and degenerate games.** What is new in Section 5 is not the half-plane structure but the *arrangement*: the $254$ distinct facet lines, the $944$ distinct polygons they cut out, and the $44$ lines carrying the one-dimensional regions. The nearest published analogue is Falmagne (107), who superimposes the boundaries of the standard strategic properties — Nash equilibrium, Pareto efficiency, maximin and the social optimum — on a two-dimensional phase space of all symmetric $2\times 2$ games, partitioning it into regions of constant strategic structure. That is the same idea for the *one-shot* game, with a handful of lines rather than hundreds, and it already draws our line $u + v = 1$ as the social-optimum boundary. The ordinal tradition of Robinson and Goforth (108) and Bruns (109) places the degenerate games on the boundaries of a non-metric layout of the $2\times 2$ games. We have found no published construction of the lower-dimensional strata of game space for a repeated-game strategy space, and none in which such a stratum is labelled by the cooperation rate of the strategies that live on it.

#### 1.8 The vanishing-noise limit of a perturbed Markov chain

**The algebra of the limit.** The statement in Section 3 that the limit coefficients are ratios of spanning-tree counts of the $\epsilon \to 0$ transition structure is the Markov chain tree theorem, which is due in the form used here to Leighton and Rivest (110) and, for perturbed families, to the $W$-graph formalism of Freidlin and Wentzell (111). Its use to extract the $\epsilon \to 0$ limit in games is three decades old (112, 113), and Fudenberg and Imhof (114) state in print that the Freidlin–Wentzell tree algorithm is the general method while offering a reduced substitute; see also (115, 116). Betz and Le Roux (117) give a cubic-time algorithm deciding which states retain non-vanishing weight as the noise fades, which is exactly the question of which of our sixteen pair states are visited in the limit and hence the mechanism behind Section 3. We claim no novelty for the algebraic form of the limit.

**Alternatives to the route taken here, and why they are worth knowing.** The $\epsilon$-expansion of a singularly perturbed stationary distribution can be obtained in closed recursive form, one aggregated chain per time scale (118, 119); the companion expansions of the fundamental, deviation and mean-passage-time matrices are given by (120, 121). On the numerical side, the GTH variant of Gaussian elimination (122) is stable in entrywise relative error regardless of how badly the chain is conditioned in norm (123), and Barlow (124) shows that nearly uncoupled chains, though catastrophically sensitive under norm-wise perturbation theory, are usually well conditioned under structured perturbations; Meyer (125) gives a cheap estimator for the relevant condition number, and Dayar and Stewart (126) examine the effect of substituting GTH elimination inside iterative aggregation–disaggregation, the usual route for nearly completely decomposable chains. Finally, parametric probabilistic model checking computes the reachability and expected-reward quantities of an $\epsilon$-parameterised finite chain as *exact rational functions of* $\epsilon$ by state elimination (127), which would deliver the limit rationals of Section 3 directly; a faster bounds-based alternative partitions the parameter space into regions without ever forming the rational function (128). Section 7 states what our extrapolation-based route buys and what it does not, and should be read with these alternatives in view.

##### 1.9 What is new here

**The demarcation in summary.** For the reader's convenience, the demarcation drawn above in summary. Inherited, and cited at the point of use: the binary strategy space with an implementation error floor, which is Lindgren's genome in the $\{\epsilon, 1-\epsilon\}$ parameterisation of Nowak and Sigmund (19, 23), together with its justification and its consequences for stability (22, 24, 25); the $\epsilon \to 0$ notion of equilibrium (26, 37); the two-parameter plane of games and the normalisation $(1, u, 1+v, 0)$ (12, 18); the exhaustive enumeration of the $2^{16}$ space (25, 26); the memory-1 self-play classification of §2, which is the diagonal of Table 1 of (32) and is given again by (33); the count 7639 of strategies at $\gamma = 1$ (26); the fact that the set of games supporting a given profile as an equilibrium is a polyhedron (102), with its single-agent antecedent for the reward functions making a policy optimal (100) and the determination of all symmetric stage games admitting each reactive equilibrium (105); the weakness of repeated-game equilibria (54, 58); the memory-2 cost threshold $b/c \geq 3/2$ (25); the existence of socially optimal memory-1 equilibria in all social dilemmas (9); alternation as the social optimum above $u+v=1$ (80, 82); the distinguished status of the line $S=T$ in memory-2 turn-taking dynamics, there the line on which alternation gives way to a consistent strategy (83); the spanning-tree form of the $\epsilon \to 0$ limit (110, 112); the failure of gain-improvement alone to certify average-reward optimality in the multichain case (48); and the fact that the $\epsilon = 0$ control problem is not the limit of the $\epsilon > 0$ problems (49).

New here, to the best of our knowledge:

1. *Limit* $\epsilon \to 0$. The memory-2 self-play spectrum: 475 distinct one-move distributions and 229 distinct

cooperation rates, all exact rationals; the two empty end bands $(0, 1/10)$ and $(9/10, 1)$, each of width exactly $1/10$; and the identification of the $9/10$ ceiling as a basin-weight ratio between two absorbing states rather than a cycle (§2).

2. *Limit* $\epsilon \to 0$. The exact Nash region $Z(p)$ in the plane of games, computed as a polyhedron in exact rational arithmetic for every one of the 65536 binary memory-2 strategies against all 65535 deviations — $4.29 \times 10^9$ limiting pair distributions (Section 3).

3. *Both regimes, and the contrast between them.* That the reduction to sixteen one-component deviations, valid at every $\epsilon > 0$, becomes vacuous in the limit and must be abandoned, so the limit computation compares every resident against all 65535 rivals. The mechanism is not special to memory-two: for a memory-$n$ resident, any component governing a state that self play never visits contributes a constraint reading $0 \leq 0$. What is claimed here is not the general possibility — that an $\epsilon = 0$ control problem need not be the limit of the $\epsilon > 0$ problems is Bielecki and Filar's (49), cited above — but that it disables the equilibrium test this literature uses (Section 3).

4. *Limit* $\epsilon \to 0$. That no binary memory-2 strategy is a strict equilibrium of the limit game — now by exhaustive exact certificate — and the methodological consequence that no uncalibrated floating-point sign test can decide the limit condition at all (Section 4).

5. *Limit* $\epsilon \to 0$. The census of dimensions — 23861 full-dimensional, 3774 one-dimensional, 4875 single points, 33026 empty — and of the arrangement they form: 944 distinct polygons, 254 distinct facet lines, and every one of the full-dimensional regions unbounded. The lines are not all concurrent — they run in 68 distinct directions and no more than 16 of them meet at any one point — so the unboundedness is a fact about the regions and not an automatic consequence of every line passing through a single point, which would make every region a cone (Section 5).

6. *Limit* $\epsilon \to 0$*, with a fixed-*$\epsilon$ *rider.* That maximum efficiency is attainable at *every* game in the plane, in all four families, by a binary memory-2 strategy that is simultaneously an equilibrium *of the* $\epsilon \to 0$ *limit game*, with efficiency decided exactly and with the reference optimum following the switch at $u + v = 1$ — and that the qualification is not a formality, since at every fixed $\epsilon > 0$ the guarantee fails on two classes of directions that are both empty in the limit: those carrying no equilibrium at all, contained in the Snowdrift quadrant exactly rather than asymptotically, and those carrying equilibria of which not one is efficient. Their exact densities are the first two of the four-kinds item below, item 11 (Sections 5 and 6).

7. *Limit* $\epsilon \to 0$. The quantitative census of how much company the efficient equilibrium keeps: 299 to 22069 equilibria at any game in the plane — the maximum attained on an unbounded region of positive area, and equal to the maximum over generic games (Section 5) — median 1568 and median sub-maximal fraction 0.493, both over the square $|u|, |v| \leq 8$, and the per-family breakdown (Section 5).

8. *Limit* $\epsilon \to 0$. $\Lambda$, the tie set: that the one-dimensional regions lie on exactly 44 lines, and that 31 of those lines carry strategies of a single cooperation rate — $3u - v = 1$ only $\gamma = 1/4$, $u - 3v = 1$ only $3/4$, $u - 2v = 1$ only $2/3$, $u - v = 1$ only $1/2$. Nothing in the construction imposes that rigidity (Section 5).

9. *Limit* $\epsilon \to 0$. The exact partition of the alternation corner, $3072 = 1792 + 1280$, with the mechanism: every alternator either holds an open set of games or sits exactly on $S = T$ (Section 5).

10. *Every error rate, limit and fixed alike.* That no binary memory-1 strategy can be efficient above the switch line $u + v = 1$, at any error rate, and that the ladder of §2 is the whole reason: in self play a resident at $w$ earns a payoff affine in $u + v$ with slope $w_3$, attaining the optimum above the line needs slope $\frac{1}{2}$, and the largest $w_3$ anywhere in the memory-1 space is $\frac{1}{4}$ — the equality case being exactly the alternation corner, on which memory-2 puts 3072 strategies and memory-1 none (§2 and Section 5).

11. *Fixed* $\epsilon > 0$. The four kinds of game, each with an exact density, measured over the directions at infinity — the only window-free notion of a fraction of games on an unbounded plane. Games with no equilibrium, with equilibria of which not one is efficient, with both kinds, and with none but efficient ones occupy $\frac{1}{4} - (\tau + \sigma + \varrho)/\pi$, $\frac{3}{8} + (3\tau + 2\sigma + 2\varrho)/2\pi$, $\frac{1}{4} - \tau/\pi$ and $\frac{1}{8} + \tau/2\pi$, exactly at every $0 < \epsilon < \frac{1}{2}$, disjoint and summing to 1, where $\tau = \arctan[\epsilon/(1-\epsilon)]$ is the tilt a positive error rate puts on *ALLC*'s and *ALLD*'s boundaries, $\sigma$ is the width of *one* of the two interior slivers, and $\varrho$ — identically zero up to $\epsilon = \frac{1}{4}$, positive beyond, a threshold the all-rate certification uncovered — is the width of one of the two further intrusion arcs of Section 6; each is a statement about directions and not about every game. They are exact because $Z_\epsilon(\mathit{ALLC})$ and $Z_\epsilon(\mathit{ALLD})$ are single half-planes tilted off their own axes by $\tau$, and nothing among the 65536 adds an arc of directions to *ALLC*'s cone. All four jump at $\epsilon = 0$, where they stand at 0, 0, $\frac{5}{8}$ and $\frac{3}{8}$, and the jumps cancel; every memory-1 density is continuous there, two equal to memory-2's at every $\epsilon > 0$ and two differing by exactly the two interior slivers, which the extra round of memory moves from the first kind into the second (Section 6, Table 2 and Main Figure 5).

12. *Fixed* $\epsilon > 0$. That the ladder of item 1 dissolves at a positive error rate, and by how much: at $\epsilon = 1/100$ the 65536 strategies realise 14300 distinct self-play distributions and 14241 distinct cooperation rates, all exact rationals, against 475 and 229 in the limit. Both empty end bands close: the extreme rates become $\epsilon$ and $1 - \epsilon$, and 2380 levels lie strictly between $9/10$ and $1 - \epsilon$, with as many mirrored below $1/10$. Binary memory-1 has 11 rates at the same error rate, against 5 in the limit. So the rungs and the gaps that organise the limit spectrum are properties of the limit and not of the strategy space — though the degeneracy is not, since 14300 points is a fifth of the space and the largest class still holds 864 strategies (§2).

13. *Fixed* $\epsilon > 0$. That the weakness of item 4 has a sharp converse, and that strictness at a positive error rate is only ever interior. That an equilibrium *interior* to its fixed-$\epsilon$ region is strict is not ours — §1 credits it to Boyd (22) and to Hilbe et al at $\epsilon = 0.01$ (25). What is ours is where it stops, and what the limit then does.

It stops at the lower strata: a region with empty interior keeps some deviation constraint tight throughout it, so each of the $564 + 1907$ residents of item 14 whose fixed-$\epsilon$ region is a line or a single game ties its best deviation *wherever* it is an equilibrium — 2471 of the 3521, with every full-dimensional resident weak on its own boundary besides. And the converse of item 4 is sharp: a resident that is an equilibrium of the limit game *alone* is not left merely non-strict at $\epsilon > 0$ but strictly *beaten*, by a gain of order $\epsilon$, which is the mechanism behind the collapse in the next item (Section 4).

14. *Fixed* $\epsilon > 0$. The census at a positive error rate, exact and over the whole plane rather than a box: 1050 full-dimensional, 564 one-dimensional, 1907 single points and 62015 empty at $\epsilon = 10^{-4}$, $10^{-3}$ and $10^{-2}$, and $1050/563/1907/62016$ at $10^{-1}$ — so the four nonempty sets have union 3521 and intersection 3520, one resident separating them. Three things in it are not small corrections to the limit. *Nine in ten* of the full-dimensional limit regions die outright: 21700 of the 23861 are equilibria at no game whatever at $10^{-4}$, $10^{-3}$ and $10^{-2}$, and 21701 at $10^{-1}$ — the same one resident that separates the union from the intersection above. The tie set shatters: the 564 survivors are scattered over 492 separate lines where the limit's 3774 shared just 44. And the single-game stratum, which the limit fills with 4875 strategies at 52 games, does not empty but holds 1907 strategies at 1128 games, 596 of them at the one game $\big((1-\epsilon)/(1-2\epsilon),\ \epsilon/(1-2\epsilon)\big)$, which is $(9999/9998,\ 1/9998)$ at $\epsilon = 10^{-4}$: the two counts are flat in $\epsilon$, that location is not (Section 5).

15. *Fixed* $\epsilon > 0$. That the equilibrium count at a single game runs from 0 to 1024 at every one of the four error rates, against 299 to 22069 in the limit — so the floor falls to nothing while the ceiling falls twentyfold — and that the maximum is settled by recession cones rather than by an arrangement, since a convex region reaches arbitrarily far along a direction exactly when every facet normal opposes it. Exactly 26 of the 1050 cannot join the maximum, and they sit at three self-play points, one of them *TFT*'s (Section 5).

16. *Fixed* $\epsilon > 0$. That no perfect alternator holds a two-dimensional region at any positive error rate. Of the 3072, none is full-dimensional, 26 hold a line, 108 a single game and 2938 nothing at all — identically at $10^{-4}$, $10^{-3}$, $10^{-2}$ and $10^{-1}$. Since perfect alternation is the only way to attain the optimum above the switch line, this is why the $90^\circ$ of directions above it that carry equilibria of both kinds in the limit — the upper half of the Prisoner's Dilemma and the upper half of Harmony — carry none that is efficient once $\epsilon > 0$. With the lower half of the Prisoner's Dilemma, where instead no fully cooperating strategy survives as an equilibrium in the far field, that is the $135^\circ$ which passes *entire* from the coexistence kind into the kind that carries equilibria of which none is efficient: the largest single transfer among the four (Sections 5 and 6).

17. *Fixed* $\epsilon > 0$. The efficient layer at a positive error rate: 245 strategies are efficient equilibria on a two-dimensional set of games, 154 on a line and 390 at a single game, a subtotal of 789 that is *flat* across all four error rates, computed separately at each. The structure behind the flatness is that at any one $\epsilon$ the 964

half-planes bounding the Nash regions $Z_\epsilon(p)$ of the 245 carry only two normal directions, $(-1,(1-\epsilon)/\epsilon)$ and $((1-\epsilon)/\epsilon,-1)$, so every region is a translate of one pointed cone and no intersection of them can be empty; the cone itself turns with $\epsilon$, which is why the strata repeat while nothing about their position does. Below the switch line the union of the 245 full-dimensional regions is the explicit wedge (17), whose one region facet is contributed by $AON_2$ alone — at $\epsilon = 10^{-1}$ it acquires a further facet. And the maximum of 245 at one game is attained only in the far field: at $\epsilon = 10^{-4}$ its nearest point is $|u| = 2.000080 \times 10^7$. The strata are flat in $\epsilon$; not one of these locations is (Sections 5 and 6).

#### 2 The ladder of cooperation levels

Everything below is a statement about where a strategy sits on a discrete ladder of achievable cooperation levels, so we construct the ladder first. The object is elementary — a monomorphic population playing itself — but the fact that the achievable levels form a *finite* set of exact rationals with conspicuously empty end bands is what makes the equilibrium census of §5 a statement about levels rather than about a continuum.

**The one-move distribution.** A memory-2 strategy plays a copy of itself. At $\epsilon > 0$ the 16-state chain is ergodic; marginalising its stationary distribution onto a single round gives

$$w \;=\; (w_1, w_2, w_3, w_4) \qquad \text{over} \qquad CC,\; CD,\; DC,\; DD, \tag{9}$$

The focal player's action is first, and we take $\epsilon \to 0$. In self play the two seats are exchangeable: the co-player reads the same history from the other seat, which relabels the states by the involution $CC \mapsto CC$, $CD \leftrightarrow DC$, $DD \mapsto DD$ applied to both rounds. That relabelling carries the chain onto itself, so

$$w_2 \;=\; w_3 \qquad \text{exactly, for every strategy and every } \epsilon. \tag{10}$$

The distribution therefore has two degrees of freedom and lives in a triangle with corners $w_4 = 1$ (mutual defection), $w_1 = 1$ (mutual cooperation) and $w_2 + w_3 = 1$ (perfect alternation); SI Figure 1 draws it with mutual defection at bottom left, mutual cooperation at bottom right and alternation at the apex, and SI Figure 2 draws the same two families at $\epsilon = 1/100$ — where the ladder below dissolves into 14300 points and 14241 levels, the corners empty, and the gap of §2 closes. Everything in this section is therefore a statement about the limit. The scalar we report is the self-play cooperation rate

$$\gamma \;=\; w_1 + w_2 \;=\; w_1 + \tfrac{1}{2}(w_2 + w_3), \tag{11}$$

a linear functional on that triangle, equal to 1, 0 and $\frac{1}{2}$ at the three corners. The definition of $\gamma$, the ordering of the four outcomes, and the reading of $\gamma$ in self play as $\epsilon \to 0$ are those of Baek et al (129), who call a strategy whose $\gamma(p, p)$ tends to one a *self-cooperator*. Conjugating $C \leftrightarrow D$ maps $w \mapsto (w_4, w_3, w_2, w_1)$ and hence $\gamma \mapsto 1 - \gamma$, so every count below is mirror symmetric about $\gamma = 1/2$.

**Memory-1: five levels, and they are already in print.** The 16 binary memory-1 strategies realise only six distinct points of the triangle, and projecting on $\gamma$ leaves five rates, which are exactly the quarters:

$$\gamma \in \{0, \tfrac{1}{4}, \tfrac{1}{2}, \tfrac{3}{4}, 1\}, \qquad \text{with 3, 2, 6, 2, 3 strategies.} \tag{12}$$

These numbers are not new. Table 1 of (32) — the technical companion promised by (23) — prints the full $16 \times 16$ matrix of $\epsilon \to 0$ one-move distributions for exactly this construction, and its diagonal is (12) together with the six points; Table 2 of Kim, Choi and Baek (33) gives the same sixteen distributions again, in leading-order $\epsilon$ form. What those papers do not do is read the table as a spectrum: none remarks that no monomorphic binary memory-1 population has $\gamma \in (3/4, 1)$. That reading, and the observation that the rate does not determine behaviour — the two points at $\gamma = 1/2$ are behaviourally opposite, one alternating half the time and the other never — is what SI Figure 1 adds.

**Memory-2: $475$ points and $229$ levels.** The $2^{16} = 65536$ binary memory-2 strategies realise $475$ distinct points of the triangle and $229$ distinct cooperation rates, all exact rationals. The largest denominator occurring in any *coordinate* is $146$, attained by $(29/146, 37/146, 37/146, 43/146)$ and its conjugate; the largest denominator among the $229$ *rates* is $106$. Rates $1$ and $0$ carry $7639$ strategies each, rate $1/2$ carries $15102$, and the five rates $\{0, \frac{1}{3}, \frac{1}{2}, \frac{2}{3}, 1\}$ account for $63\%$ of the space. The count $7639$ is in print: it is the size of the memory-2 "efficient" set of Murase and Baek (26), whose efficiency criterion is exactly $\gamma = 1$ in the $\epsilon \to 0$ limit. The rest of the spectrum, as far as we are aware, is not.

**The two end bands.** Walking down from full cooperation the rates begin $1, \frac{9}{10}, \frac{8}{9}, \frac{7}{8}, \ldots$, so *no monomorphic binary memory-2 population has* $\gamma \in (9/10, 1)$. With its mirror $(0, 1/10)$ these are by a wide margin the two widest gaps in the spectrum, each exactly $1/10$; the next widest is $1/42$ and the narrowest is $1/4270 = 2.34 \times 10^{-4}$. Memory-2 therefore raises the ceiling on sub-perfect monomorphic cooperation from $3/4$ to $9/10$.

**What the ceiling is made of.** The $\gamma = 9/10$ point is not a cycle. It is carried by exactly three strategies, and for each the limiting distribution is supported on exactly two of the sixteen states: mutual cooperation carrying $9/10$ and mutual defection carrying $1/10$. The level is therefore a *basin-weight ratio between two absorbing states* of the $\epsilon \to 0$ chain, fixed by the spanning-tree counting of §3 over minimal-error escape paths, and the ceiling question becomes a question about which such ratios are achievable: why can the weight on the defective absorbing state not lie in $(0, 1/10)$? We do not have the answer. Two absorbing states is in any case a small-denominator mechanism: $16384$ strategies have both $CC|CC$ and $DD|DD$ absorbing, only $526$ have a limit supported on the two absorbing states alone, and that family realises $37$ distinct points with largest coordinate denominator $16$. The denominator $146$ comes from the opposite regime, fifteen states in the limiting support and five to eight competing recurrent classes. Simple supports give simple rationals; the arithmetically awkward rates are where many recurrent classes compete.

**The point set refines the spectrum.** Because $\gamma$ is a single linear functional, the 229 rates are a one-dimensional shadow of the 475 points, and the refinement is substantial: $\gamma = 1$ and $\gamma = 0$ are single corners, but $\gamma = 2/3$ is 13 behaviourally distinct residents and $\gamma = 1/2$ is 15. *The level a population sits on does not determine its behaviour.* This is the fact that makes §5's single-level rigidity a statement with content: a line of $\Lambda$ that carries one rate may still carry many distinct behaviours, and the 1280 alternators on $S = T$ are remarkable precisely because they are one rate *and* one point.

#### 3 The Nash region of a strategy is a polyhedron, and the limit is taken exactly

**Games, in one normalisation.** A symmetric two-player game is fixed by its payoffs $R, S, T, P$ for the outcomes $CC$, $CD$, $DC$, $DD$ of one round, the focal player's action written first. *Positive* affine transformations of the payoffs change neither the best replies nor the efficiency ordering, so for any game with $R > P$ we may set $R = 1$ and $P = 0$ and write

$$(R, S, T, P) = (1,\, u,\, 1 + v,\, 0), \qquad u = S, \quad v = T - 1. \tag{13}$$

The plane of $(u, v)$ then contains every game with $R > P$, with the four classical families in its four quadrants: the Prisoner's Dilemma at $u < 0,\ v > 0$; snowdrift at $u > 0,\ v > 0$; the Stag Hunt at $u < 0,\ v < 0$; and harmony at $u > 0,\ v < 0$. This is the normalisation and the reading of LaPorte et al (18). Two lines through it matter throughout. On $u + v = 1$ the attainable social optimum switches from mutual cooperation, $R$, to alternation, $(S + T)/2$, which is the ST-reciprocity boundary of Tanimoto and co-workers (80, 82); and $u - v = 1$ is $S = T$, the perfectly symmetric anti-coordination game in which neither role of an alternation pays more than the other.

**The deviation gain is affine in the game.** For a pair of strategies $(q, p)$ playing at noise floor $\epsilon$, let $\nu(q, p) = (\nu_1, \nu_2, \nu_3, \nu_4)$ be the one-move distribution of the pair over $CC, CD, DC, DD$ with the focal player $q$'s action written first, and let $w = \nu(p, p)$. In the normalisation (13) the focal payoff is $E = \nu_1 + u\,\nu_2 + (1 + v)\,\nu_3$, so the gain a deviator $q$ makes against a resident $p$ is

$$E(q,p) - E(p,p) \;=\; \alpha_q + u\,\beta_q + v\,\delta_q, \qquad \begin{aligned} \alpha_q &= (\nu_1 - w_1) + (\nu_3 - w_3), \\ \beta_q &= \nu_2 - w_2, \\ \delta_q &= \nu_3 - w_3. \end{aligned} \tag{14}$$

The three coefficients are built from outcome frequencies alone. *None of them contains $u$ or $v$*: the transition matrix of the pair chain is a function of the two strategies and of $\epsilon$, never of the payoffs, and the payoffs enter only through the linear reward vector. Each deviation is therefore a half-plane in the plane of games, and

$$Z(p) \;=\; \big\{\, (u,v) \;:\; \alpha_q + u\,\beta_q + v\,\delta_q \le 0 \quad \text{for all } q \,\big\} \tag{15}$$

is a closed convex polyhedron: *the set of games at which $p$ is a Nash equilibrium.* This is why the whole plane costs one computation. A single sweep over the 65535 deviations of one resident settles every game simultaneously, and there are no games left to sample.

**This structure is known; the computation is what is new.** That the set of games supporting a given equilibrium is cut out by one linear inequality per deviation, and is hence a polyhedron, is a standard fact in several guises — the feasible reward set of inverse reinforcement learning (100, 101), the inverse Nash set of inverse game theory (102), and the map from a reactive equilibrium back to the stage games admitting it (105). LaPorte et al (18) give precisely such regions for five classes of memory-1 equilibrium. We claim no novelty for (15). What is new is that it is evaluated exhaustively and exactly, for every strategy of a $2^{16}$ space against every deviation, so that the resulting decomposition of the plane — its facet lines, its polygons, and above all its lower-dimensional strata — can be read off rather than guessed.

**All deviations, not sixteen.** Against a fixed memory-2 opponent a deviator faces an average-reward Markov decision process on the same 16 states, and a best reply may be taken within the same memory class (38, 40, 41). At $\epsilon > 0$ every transition probability the shift permits is at least $\epsilon^2 > 0$, and each state has all four of its successors available, so any state is reached from any other in two steps and the chain is irreducible under every policy, and the policy-improvement theorem then makes unimprovability equivalent to optimality (47): it is enough to test the 16 one-component deviations. That reduction is what makes the fixed-$\epsilon$ question cheap, but it is not available for $\epsilon \to 0$. In this limit the coefficients $\alpha, \beta, \delta$ of a deviation that touches only states the pair never visits all tend to zero together, the constraint becomes vacuous, and the test silently accepts residents that a genuine multi-state deviation defeats. The underlying fact is classical — in the limit the chain is multichain, and gain-unimprovability no longer certifies optimality (48), the $\epsilon = 0$ control problem not being the limit of the $\epsilon > 0$ problems (49, 50). In direct reciprocity it has been met but not named: Hilbe et al (25) restrict their equilibrium conditions to "consistent" histories exactly because in the limit of rare errors some histories are never experienced, and Glynatsi et al (37) note that under vanishing errors *almost all* partner strategies remain only approximate equilibria. Every result below for $\epsilon \to 0$ therefore uses all 65535 deviations of every one of the 65536 residents: $4.29 \times 10^9$ limiting pair distributions.

**The limit exists, and it is computed exactly.** For each $\epsilon > 0$ the pair chain is irreducible, so by Cramer's rule every stationary component is a ratio of determinants, hence a rational function of $\epsilon$ with rational coefficients; stationary probabilities stay in $[0, 1]$, so that rational function is bounded at $\epsilon = 0$ and analytic there, and $\nu(q, p)$ admits an expansion $\nu_0 + c_1\epsilon + c_2\epsilon^2 + \ldots$. The limit $\nu_0$ is itself a ratio of integers given by the

Markov chain tree theorem (110, 111): every transition probability of the pair chain is $\epsilon^k(1-\epsilon)^{2-k}$ with $k$ the number of trembles, so each tree-theorem numerator is a polynomial in $\epsilon$ of degree at most 30 whose lowest-order coefficient counts the spanning in-trees of minimal total tremble count, and

$$\nu_{0,i} \;=\; \frac{c_i\,[\,m_i = m\,]}{\sum_{j:\,m_j=m} c_j}, \qquad m = \min_j m_j,$$

with $m_i$ that minimal count for root $i$ and $c_i$ how many trees attain it. The use of the tree formula to take a small-noise limit in a game is standard (112–114); what is done here is to evaluate it *exactly*, in integer arithmetic with a proved coefficient bound, for all $4.29 \times 10^9$ pairs, so that every limiting coefficient of (14) is an exact rational by construction. How the integers are computed, the bound that makes the computation a proof, the four per-pair consistency checks, and the retrospective audit of the earlier extrapolation route are in §7. Reducibility of the $\epsilon = 0$ chain never needs a case analysis: the polynomial carries it, which is precisely the Freidlin–Wentzell picture in algebraic form.

**Which deviations decide each test, now as statements.** Two results make the two computations sufficient as well as necessary, against *all* behavioural deviations — arbitrary history-dependent, randomised, unbounded-memory rules, with the same implementation error applied to intended actions.

**Proposition 1.** *Fix a resident p, a game $(u, v)$ and $0 < \epsilon < \frac{1}{2}$, and let $g(\sigma)$ be the long-run average payoff of a deviator $\sigma$ against p. Then* $\sup_{\sigma \text{ behavioural}} g(\sigma) = \max_{q \in Q} g(q)$ *over the $2^{16}$ binary strategies Q, and the following are equivalent: (i) no behavioural deviation earns more than $g(p)$; (ii) no $q \in Q$ earns more than $g(p)$; (iii) none of the sixteen one-flip deviations earns more than $g(p)$.*

*Proof.* Against a fixed memory-two resident the deviator faces a finite average-reward MDP on the sixteen states: rewards and transitions depend on the history only through the last two outcomes, and the stationary deterministic policies — intended actions per state, realised with error $\epsilon$ — are exactly $Q$. An average-optimal stationary deterministic policy exists and is optimal among all history-dependent randomised policies (47); at $\epsilon > 0$ every policy's chain is irreducible (each state has its four successors with probability at least $\epsilon^2$, and any state is reached from any other in two steps), so the MDP is unichain and the optimal gain is constant in the initial state. This gives the display and (i)⇔(ii); (ii)⇒(iii) is trivial. For (iii)⇒(ii), let $g_p, h_p$ solve the Poisson equation for $p$ and write the Bellman slack $\phi_p(s,a) = r(s,a) + \sum_{s'} P(s' \mid s,a) h_p(s') - g_p - h_p(s)$, so $\phi_p(s, p(s)) = 0$. The gain-difference identity gives, for any stationary deterministic $q$, $g(q) - g(p) = \sum_s \nu_q(s)\,\phi_p(s, q(s))$. If $p$ is not optimal then, the MDP being unichain, the average-reward optimality equation fails at some state, i.e. $\phi_p(s_0, a_0) > 0$ for some $s_0$ and $a_0 \neq p(s_0)$; the one-flip deviation $q$ that plays $a_0$ at $s_0$ and copies $p$ elsewhere has $g(q) - g(p) = \nu_q(s_0)\,\phi_p(s_0, a_0) > 0$, because $\nu_q(s_0) > 0$ by irreducibility. So if (ii) fails, (iii) fails. □

**Lemma 1.** *$p$ satisfies* (8) *against every behavioural deviation — with* $\limsup$ *in place of the limit for deviations whose payoff does not converge — if and only if* $\lim_{\epsilon\downarrow 0}[\pi_\epsilon(q,p) - \pi_\epsilon(p,p)] \le 0$ *for every* $q \in Q$.

*Proof.* Necessity: each $q \in Q$ is a behavioural deviation whose limit exists (§3, the rational-function fact). Sufficiency: for behavioural $\sigma$, Proposition 1 gives $\pi_\epsilon(\sigma,p) \le \max_{q\in Q} \pi_\epsilon(q,p)$ at every $\epsilon$; the family $Q$ is finite and each $\pi_\epsilon(q,p)$ converges, so the maximum converges to the maximum of the limits, and $\limsup_{\epsilon\downarrow 0} \pi_\epsilon(\sigma,p) \le \max_{q\in Q} \lim_{\epsilon\downarrow 0} \pi_\epsilon(q,p) \le \lim_{\epsilon\downarrow 0} \pi_\epsilon(p,p)$. □

**Why the sixteen do not suffice in the limit.** Lemma 1 is the licence for sweeping all 65535 deviations, and no shorter list is licensed: all sixteen one-flip limits can be $\le 0$ while a multi-flip deviation gains, because the policy-improvement path from $p$ to the optimum passes through intermediate policies, and each single increment can vanish in the limit while their sum does not. The mechanism is the one described above — a component governing a state of vanishing weight contributes a constraint of vanishing coefficients — and it is why the $\epsilon = 0$ control problem is not the limit of the $\epsilon > 0$ problems (49, 50).

#### 4 Every equilibrium is weak, and what that forces

**The constraints are tight over the whole region.** In the limit game the best deviation does not lose to the resident, it *ties* it. For the resident 36873 at the game $(u, v) = (0.5, 1.5)$, where the Nash region is full-dimensional and therefore not a knife-edge, the maximum of (14) over all 65535 deviations is exactly zero: the exact computation of §7 returns deviations whose limiting gain coefficients $(\alpha, \beta, \delta)$ vanish identically, so the tie holds at every game, not merely at this one. This is not special to that resident. Every one of the 23861 residents whose region is full-dimensional admits such an identically tying deviation — never fewer than 351 of them, each recorded with a representative in the census output — and on the lower-dimensional regions no certificate is needed: a one-dimensional region keeps its antiparallel pair tight throughout, and a single-game region sits where several constraint lines meet, so the best deviation ties there by construction. Hence no binary memory-2 strategy is ever a strict Nash equilibrium of the limit game, by exhaustive exact certificate rather than by numerical observation.

**This is the classical picture, in exact form.** That repeated-game equilibria are weak, the best deviation tying rather than losing, is the content of the no-ESS line (54–56), of the neutral-stability apparatus built to replace strictness (57, 61, 62), and of the general theorem that every equilibrium of a sufficiently long repeated game admits a payoff-tying mutant (3, 58). Our contribution here is not the statement but its scope and its status: it holds for every one of the 65536 strategies, at every game in the plane, against every deviation, and it is established as an identity — the tying deviations' limiting gain coefficients vanish exactly in the integer tree-theorem computation of §7, with no tolerance anywhere.

**It does not contradict the results in which noise restores strictness.** At a fixed $\epsilon > 0$ strictness is available in this very parameterisation: mistakes permit a pure strategy to be evolutionarily stable (22), and strict memory-2 Nash equilibria are reported at $\epsilon = 0.01$ (25). Our claim is confined to the $\epsilon \to 0$ limit game, and the two directions have to be kept apart. Where a resident is already an equilibrium at a fixed $\epsilon > 0$ at a game interior to its region *and that game also lies in its limit region* $Z(p)$, its best-deviation gain is strictly negative and vanishes with $\epsilon$ — it must, since the limit gain is an exact tie — so for those residents the limit removes the strictness and nothing more. Interiority is not a formality: a region with empty interior keeps some constraint tight throughout, so the $564$ residents whose fixed-$\epsilon$ region is a line and the $1907$ whose region is a single game are weak wherever they are equilibria, at every $\epsilon$. The converse fails, and consequentially: a resident that is an equilibrium of the limit game alone is in general beaten at fixed $\epsilon$, the same computation returning a gain that is strictly *positive* and of order $\epsilon$. That is why the fixed-$\epsilon$ and the limit equilibrium sets are almost disjoint, which §5 takes up.

**The methodological consequence.** Because the binding constraints of (15) are satisfied with *equality* throughout $Z(p)$, no uncalibrated floating-point sign test can decide membership: rounding turns an exact tie into a number of either sign, so such a test neither reliably accepts nor reliably rejects an equilibrium, and a tolerance cannot be sharpened into a proof by improving the arithmetic. The question is intrinsically exact, which is why every tie in this paper is decided by the integer tree-theorem computation of §7, and it is the reason a tolerance-based census and an exact one can disagree while both being internally consistent.

**The coefficients are exact, and that is what makes the strata decidable.** The coefficients of (14) are ratios of spanning-tree counts, and a one-dimensional region always carries an exactly antiparallel pair of half-planes among its constraints — a coincidence that no floating-point representation can hold, since carrying computed digits verbatim would turn it into a sliver of width $10^{-34}$ and misclassify the region as full-dimensional. Here the coincidence is never at the mercy of digits: every coefficient is the exact integer-arithmetic limit of §7, so antiparallelity is an identity between integers and the strata are read off rather than reconstructed. A second route — extrapolation followed by rational snapping — is run alongside as an independent numerical cross-check, and §7 reports its agreement with the exact coefficients.

#### 5 The census, efficiency, and the tie set

##### 5.1 Dimensions, and the largest regions there are

**Dimensions.** Of the $65536$ binary memory-2 strategies,

| $Z(p)$ is | strategies | share |
|---|---|---|
| full-dimensional | 23861 | 36.41% |
| one-dimensional, an interval on a tie line | 3774 | 5.76% |
| a single point | 4875 | 7.44% |
| empty | 33026 | 50.39% |

All four entries count *strategies*, and the lower strata are shared far more heavily than the table suggests: the 3774 one-dimensional regions are only 165 distinct intervals, lying on the 44 tie lines described below, and the 4875 point regions are only 52 distinct games, every one of them within $\sqrt{5}$ of the origin. Half of all strategies are an equilibrium at no game whatsoever. Single-point regions are real and not an artefact: a strategy can be a Nash equilibrium at *exactly one* game, for instance at $(u, v) = (1, 0)$, which is the fully degenerate game $R = S = T$. This table has no published counterpart. The nearest prior census is the classification of Hilbe et al (25) at one fixed donation game, which reports eleven cooperative, fifteen defecting and eight self-alternating equilibria, plus a fourth class with two absorbing states, at $\epsilon = 0.01$; the numbers here are larger and differently defined because they count every equilibrium at every game, cooperative or not, in the limit rather than at fixed $\epsilon$.

**The largest equilibrium region there is, and it belongs to full cooperation.** No strategy is an equilibrium at every game: not one of the 23861 full-dimensional regions is unconstrained. Nor does any of them exceed a half-plane. The 23861 full-dimensional regions collapse to 944 distinct polygons, bounded by 254 distinct facet lines; those lines are not a pencil, running in 68 distinct directions with no more than 16 of them through any one point, so what follows is a fact about the regions and not the trivial consequence of a fan of concurrent lines. Every full-dimensional region is unbounded, so its area depends on the window, but the width it occupies among the directions at infinity does not, and the largest that width ever reaches is exactly $180°$ — attained by the 566 residents whose entire region is a single half-plane, and by no one else, since two non-redundant constraints already cut the cone below a half turn. That 566-fold tie is the window-free statement. It breaks under any window centred at the origin, and it breaks the same way under all of them, because half-planes $\{n \cdot x \leq c\}$ with $|n| = 1$ are ordered by $c$ alone. Sixteen distinct half-planes occur among the 566, and *exactly one* has $c > 0$: the half-plane $u + v \leq 1$, at $c = 1/\sqrt{2}$, held by the two residents 20111 and 24207. Their region is half the plane together with the strip between $u + v = 0$ and $u + v = 1$, which is 8319.5 of the 16384 of $|u|, |v| \leq 64$ against 8192 for any half-plane through the origin. Both cooperate fully — limiting self-play distribution $(1, 0, 0, 0)$ — and they differ from one another in a single one of their sixteen components. So the strategy that is an equilibrium at more games than any other is a fully cooperating one whose region is exactly the set of games at which mutual cooperation is the social optimum: it ceases to be an equilibrium precisely where it ceases to be efficient. For comparison *ALLC* holds $v \leq 0$, that is $T \leq R$, shared with 294 others, and *ALLD* holds $u \leq 0$, that is $S \leq P$, shared with 72; the complementary half-plane $u + v \geq 1$ is held by eight.

**Table 1: The two strategies with the largest equilibrium region, in full.** Each cell is the action the strategy takes in one of the sixteen memory states; the state is the pair (most recent outcome, the outcome before it), with the focal player's own action written first in each outcome — the coding used throughout this SI, under which the state index runs 0 to 15 as $j = 4\,(\text{recent}) + (\text{before})$ with the outcomes ordered $CC, CD, DC, DD$; **Methods** numbers the same sixteen states 1 to 16, so the conjugation that is $17 - j$ there is $15 - j$ here. Both strategies cooperate at every state whose most recent outcome is $CC$, and both are fully cooperating in self play, with limiting distribution $(1, 0, 0, 0)$ and $\gamma = 1$. They differ in exactly one of the sixteen components, the state $(DD, CC)$ in the bottom-left cell of each block: 20111 defects there and 24207 cooperates. It makes no difference to either the self-play point or the equilibrium region, which is the half-plane $u + v \leq 1$ for both. Half the plane is the largest region any of the 65536 strategies holds, and 566 of them hold it; these two come first only under the tie-break of the centred window.

| genotype 20111 | | | | | genotype 24207 | | | | |
|---|---|---|---|---|---|---|---|---|---|
| | the round before | | | | | the round before | | | |
| most recent | $CC$ | $CD$ | $DC$ | $DD$ | most recent | $CC$ | $CD$ | $DC$ | $DD$ |
| $CC$ | C | C | C | C | $CC$ | C | C | C | C |
| $CD$ | D | D | D | C | $CD$ | D | D | D | C |
| $DC$ | D | C | C | C | $DC$ | D | C | C | C |
| $DD$ | D | D | C | D | $DD$ | C | D | C | D |

The two blocks are the same pattern but for one cell. Both cooperate after a round of mutual cooperation, whatever preceded it. Both refuse after being exploited, unless the round before that was mutual defection. Both return to cooperation after exploiting the co-player, unless the round before that was mutual cooperation. They part company only at $(DD, CC)$ — most recent outcome mutual defection, the round before it mutual cooperation — a state reachable only when both players err in the same round, which is why the difference costs neither of them anything in the limit and neither region nor self-play point can tell them apart.

**At a positive error rate the title changes hands, and to the two strategies one can write down.** The same question at fixed $\epsilon$ has a different answer, and it needs no search. By (24) of §6, proved there by hand, the regions of *ALLD* and *ALLC* are *exactly* the half-planes $(1-\epsilon)u \leq \epsilon v$ and $(1-\epsilon)v \leq \epsilon u$ for every $\epsilon \in (0, \frac{1}{2})$: all sixteen one-component deviations give the one constraint, the varying factor cancelling. A half-plane occupies 180° of the directions at infinity, and no convex proper subset of the plane can occupy more, so those two attain the maximum outright. They also tie exactly, since both bounding lines pass through the origin — offset 0 — and a half-plane whose boundary passes through the origin covers exactly half of any centrally symmetric window, whatever its normal. (Offsets do *not* order half-planes by $c$ alone in a centred square: at the same $c$ a diagonal half-plane covers more than an axis-aligned one. Only the $c = 0$ case is needed here.)

Nothing else comes near, and nothing else ties. Enumerating the recession cone of every full-dimensional region at each of $\epsilon = 10^{-4}$, $10^{-3}$, $10^{-2}$ and $10^{-1}$ gives a runner-up of exactly $90° - 2\tau$ — 89.988540°,

$89.885294°$, $88.842549°$ and $77.319617°$ — which is the opening of the intersection of the two tilted half-planes, the antipode of the empty cone $K_\epsilon$ of §6. The cliff is structural: every facet of the fixed-$\epsilon$ arrangement belongs to one of the two pencils, so a region with two distinct facet normals is one half-plane from each pencil and can do no better. A third half-plane region is excluded without appeal to that enumeration's scope, since a half-plane meets every centred box in positive area and would be recorded as full-dimensional by any of the region files.

The contrast with the limit is sharper than a change of winner. In the limit 180° is a 566-fold tie which *ALLC* and *ALLD* are themselves part of, and 20111 and 24207 come first only under the tie-break: theirs is the one half-plane among the sixteen with a positive offset, so it wins in every origin-centred window, while of the other fifteen two pass through the origin and thirteen exclude it, so every one of them covers at most half of any centred window. Two things happen at once when the error rate becomes positive. The regions of 20111 and 24207 become *empty* — they are equilibria at no game whatever, at all four rates — and the 566-fold tie collapses to exactly two, so there is nothing left to break. The strategy that is an equilibrium at more games than any other in the limit is an equilibrium at none once a hand can tremble, and what is left holding the widest region is the pair whose regions can be written down by hand.

**Full cooperation, and a known threshold recovered.** Among the 23861 full-dimensional regions the leading cooperation rate is $\gamma = 1$ with 7639 strategies. One of these, 36873, has region exactly $\{u \leq 1\} \cap \{v \leq 2\}$, that is $\{S \leq R\} \cap \{T \leq 3R\}$, and the two facets are contributed by two specific deviators: one that becomes a permanent sucker and earns $S$, and *ALLD*, which earns $T/3$. On the donation line, where $(R, S, T, P) = (1 - c, -c, 1, 0)$ and hence $v = -u = c/(1 - c)$, the facet $T \leq 3R$ reads $c/b \leq 2/3$. That is exactly the memory-2 case $b/c \geq (n + 1)/n$ of the *AON*$_n$ threshold of Hilbe et al (25), obtained here without being put in, and it is the sharpest single check that the construction is right. The corresponding memory-1 statement is the $n = 1$ case of the same threshold, $b/c \geq 2$ for *WSLS* (25), equivalently the $\delta \to 1$ donation-game specialisation of the partner condition for *WSLS* in Hilbe et al (45).

#### 5.2 Efficiency: how much of it there is, and where

**Efficiency.** We use the normalisation of the companion paper (130): $E_{\max} = \max\{R, (S + T)/2\}$ and $E_{\min} = \min\{P, (S + T)/2\}$, and a monomorphic resident with limiting self-play distribution $w$ earns $\bar{E} = w_1 + u\,w_2 + (1 + v)\,w_3$, so its efficiency is $(\bar{E} - E_{\min})/(E_{\max} - E_{\min})$. This is evaluated in exact rational arithmetic on the exact $\epsilon \to 0$ self-play point of the resident, so "maximally efficient" means efficiency equal to 1 exactly and no threshold enters. Maximum efficiency means two different things on the two sides of $u + v = 1$: mutual cooperation below it, alternation above. That switch is not our observation; it is the R-reciprocity versus ST-reciprocity dichotomy of Tanimoto and co-workers (80–82), and it is the one point on which our efficiency accounting differs from that of Hübner et al (9), who take the social optimum to be mutual cooperation throughout.

**Why "efficient" has to be a limit label while "Nash" need not.** The test cannot be run at $\epsilon > 0$ instead. Errors put weight on every outcome, so every strategy falls short of $E_{\max}$ by $O(\epsilon)$, an exactly evaluated $\epsilon$-efficiency bit is identically zero, and a tolerance would let $\delta$ perform the classification silently. Throughout this document, therefore, "efficient" is the $\epsilon \to 0$ label and "Nash" is the label of whichever layer is under discussion, which puts the $\epsilon$-dependence where it physically is. The two are mixed deliberately wherever a fixed-$\epsilon$ equilibrium is called efficient, and that mixture is what (17) below counts.

**The first result.** Rasterising the exact $\epsilon \to 0$ polyhedra onto a $2001 \times 2001$ grid over $[-8, 8]^2$ — $4004001$ games — gives two facts, and both are exact counts rather than estimates. At *no* grid point is the set of equilibria empty, and at *no* grid point is the best efficiency achieved by an equilibrium less than 1: the minimum over the whole plane of "the best efficiency any Nash equilibrium achieves" is 1.000000. Maximum efficiency is therefore attainable at every game in the plane, in all four families, by a binary memory-2 strategy that is simultaneously an equilibrium *of the limit game*. That qualification is essential and not a formality: at a fixed $\epsilon > 0$ the equilibrium set is a different and much smaller object, and a large part of the plane carries no memory-2 equilibrium at all (the grey of Main Figure 2b). SI Figure 3 draws the same picture at $\epsilon = 10^{-2}$ and $10^{-1}$. That part has a clean shape and an exact measure: rasterising the full-dimensional $Z_\epsilon(p)$ over $[-8, 8]^2$ on the same $2001 \times 2001$ grid the limit result uses, the games carrying no equilibrium are $0.2391$ of the square at $\epsilon = 10^{-2}$, $0.2419$ at $10^{-3}$ and $0.2420$ at $10^{-4}$ — and they are *contained in the Snowdrift quadrant*, of which they are $0.9566$, $0.9678$ and $0.9681$, while exactly none of the Prisoner's Dilemma, the Stag Hunt or Harmony is left without an equilibrium. The companion statement — that of the games which do keep an equilibrium, a large share keep none that is efficient — is the same computation with the efficient count in place of the total: on the disk the figures draw it is $32.6\%$ at $\epsilon = 10^{-4}$ and at $10^{-3}$ and $33.2\%$ at $10^{-2}$ — SI Figure 3 renders the same disk on a finer grid and gives $33.3\%$ — and on the square $|u|, |v| \le 8$ it is $38.4\%$, $38.5\%$ and $39.1\%$. Both readings are window-dependent and both are conditional on the game keeping an equilibrium at all; the window-free version of the same statement is (33) of §6, $\frac{3}{8} + (3\tau + 2\sigma + 2\varrho)/2\pi$ of all directions ($\varrho = 0$ at every rate reported here) and $0.5000509$ of the directions that keep an equilibrium — just over one half, where the disk reads $32.6\%$ and the square $38.4\%$. The main text names the families as well: Harmony above $u + v = 1$ keeps not one efficient equilibrium off a set of measure zero, and the Prisoner's Dilemma keeps none in the far field but for such a set. The statement can be sharpened from "at least one" to a count: the number of strategies that are simultaneously a Nash equilibrium and maximally efficient is never below 8, the minimum being attained on the Prisoner's Dilemma wedge lying immediately *below* $u + v = 1$, between $v = 2$ and $u + 2v = 2$; and it is 7639 over the whole region $S \le R, T \le R$, with a single exception at the degenerate game $(1, 0)$. That count, over the whole plane and in both notions of equilibrium, is Main Figure 3: the drop from the limit's 8–7639 to the far narrower range a positive error rate leaves is the same collapse this section reports, seen strategy by strategy rather than game by game.

**What was already known about that, and what is added.** The qualitative statement is not new at memory-1: Hübner et al (9) prove that socially optimal cooperation can be sustained as a Nash equilibrium in every pairwise social dilemma, and Hilbe et al (25) exhibit memory-2 equilibria at the mutual-cooperation payoff in the donation game. Three things are added here. First, the attainable optimum is tracked across $u+v=1$, so that above that line the equilibrium we exhibit is an alternating one earning $(S+T)/2$ and not a cooperative one earning $R$; in that region the cited results do not speak to the social optimum. Second, the statement is proved for a *finite, exhaustively enumerated* strategy space of sixteen bits rather than by constructing a suitable strategy in a continuous space, so no auxiliary parameter is available to be tuned. Third, the region of validity is the whole plane and is established by exact polyhedra rather than by a lattice of games; earlier lattice computations, including our own, could only ever report what they happened to land on.

**The second result, which is the informative one.** Existence, then, is settled and carries no information; what varies over the plane is how much company the efficient equilibrium keeps. Main Figure 2a plots, at each game, the number of the 65536 strategies whose exact *full-dimensional* polyhedron (15) contains that point; the 3774 one-dimensional and 4875 point regions are Lebesgue-null and are not counted there, which matters only on the tie lines themselves. The count runs from 299, at $(u,v) \approx (-1.25, 2.03)$ deep in the Prisoner's Dilemma, to 22069 in the Stag Hunt, and it is piecewise constant on large pieces: the 254 facet lines cut the plane into 22872 faces, the count is constant on each and takes only 3151 distinct values across all of them, and every boundary in the panel is one of those lines, so the visible structure is the arrangement itself. The four families are ordered, brightest to darkest, Stag Hunt, then harmony and snowdrift, then the Prisoner's Dilemma; the per-family medians are 533 for the Prisoner's Dilemma, 1568 for snowdrift, 1571 for harmony and 22043 for the Stag Hunt. And no part of the panel is empty.

**The distribution over games is bimodal, with an exact mechanism.** Weighted by area, the counts fall into two modes separated by an empty gap: no game anywhere carries between 4849 and 7639 equilibria. *Every share quoted in this paragraph and the two that follow is of the square* $|u|, |v| \leq 8$*, on which each of the four quadrants is exactly a quarter*; the shares are not window-independent, and on the disk the figures draw — which compactifies the whole plane — the same quantities read 33.9%, 86.8%, 48.7% and 68.1% where the square gives 37.5%, 96.9%, 53.4% and 90.7%. The gap is the largest in the achieved counts by a wide margin — a ratio of 1.58 against 1.06 for the next largest — and it splits the square 71.85% below to 28.15% above. The upper mode is not a diffuse cloud but an exactly delimited region: it is precisely the set of games with $u \leq 1$ and $v \leq 0$, that is $S \leq R$ and $T \leq R$, comprising the closed Stag Hunt quadrant together with the strip $0 < u \leq 1$, $v \leq 0$ of harmony. Those are exactly the games in which mutual cooperation is already a Nash equilibrium of the *one-shot* game and is simultaneously the social optimum, so every one of the 7639 strategies with self-play cooperation rate $\gamma = 1$ becomes an equilibrium at once — and indeed the number of *maximally efficient* equilibria is exactly 7639 at every game of the upper mode but one. The bimodality is that family switching on, not a statistical accident. The one exception is the degenerate game

$(u, v) = (1, 0)$ itself, where $R = (S + T)/2$ so that mutual cooperation and alternation are both optimal — and where efficiency asks no more than that mutual defection carry no weight, $w_4 = 0$, so the maximally efficient count is 14757: the whole $\gamma = 1$ family, all 3072 perfect alternators of (18), and a further 4046 strategies belonging to neither. At that one game the limit-Nash set and the $w_4 = 0$ set coincide exactly, so every equilibrium there is maximally efficient; 9431 of the 14757 hold a full-dimensional region and the rest a line or the point itself. The median count is also a plateau rather than a central tendency: 1568 is not only the median over the plane but the single most common value, attained at 653449 of the $2001^2$ grid points of that square, 16.3% of it, every one of them in snowdrift, where it covers 65.4% of the quadrant. So "the median game carries 1568 equilibria" is a statement about the generic snowdrift game rather than an average of dissimilar things, and the 1568 are the alternators of (18), all of them maximally efficient.

**Efficiency resolves the same equilibria, and its median is a different statistic.** Main Figure 3 counts, at each game, the equilibria that are maximally efficient. Dividing that count by the total of Main Figure 2a gives the share of a game's equilibria that are maximally efficient, which is what Main Figure 4 draws and SI Figure 3's third row at the two further error rates. The rest of this paragraph describes the part left over, the share that is *sub*-maximal. That is the complement of the first result: since an efficient equilibrium exists everywhere, it is never a statement about existence, only about how heavily that equilibrium is outnumbered. Snowdrift is almost uniformly zero, the Stag Hunt uniformly intermediate, and the largest values are a wedge of the Prisoner's Dilemma and a wedge of harmony. Per family the medians are 0.807 for the Prisoner's Dilemma, 0.653 for the Stag Hunt and 0.000 for both snowdrift and harmony; snowdrift never reaches 0.48 anywhere, and the Stag Hunt is confined to 0.1079–0.654. The sharpest feature is the discontinuity along $u + v = 1$, and it is a change in the question rather than in the strategies: below the line the reference optimum is mutual cooperation, above it alternation. The two families straddling the line respond oppositely. In harmony, crossing upward turns cooperative equilibria from optimal into sub-optimal without changing them, the median sub-maximal fraction going from 0.000 below the line to 0.832 above it. In the Prisoner's Dilemma the effect runs the other way, because above the line alternation becomes the target and some of the equilibria are alternators: the median falls from 0.907 below the line to 0.563 above it, and the global maximum of the whole map, 0.993 — exactly 1083/1091 — is attained *below* $u + v = 1$.

**Two spikes and a spread.** Weighted by area the sub-maximal fraction is not a hump around its median but two spikes and a spread. The larger spike is at zero: at 37.5% of all games *every* equilibrium is maximally efficient, which is 96.9% of the snowdrift quadrant and 53.4% of harmony and no part at all of the Prisoner's Dilemma or the Stag Hunt. The second spike, 14.1% of the plane, is dominated by the single value $14430/22069 = 0.6539$, held on 9.4% of the plane: that is the deep Stag Hunt plateau of Main Figure 2a seen from the efficiency side, where 22069 strategies are equilibria and exactly 7639 of them are maximal, so the ratio is one constant. Those two spikes together are 51.7% of the plane and the remaining 48.3% is spread thinly between them. The median of this quantity, 0.493, must therefore be read differently from the median count above: it falls in the low ground between the two spikes, on $1.23\%$ of the plane, so it is an honest fifty-fifty split point and a poor description of any actual game. The plane is not made of games at which about half the equilibria are inefficient, but of a large set at which none is and a large set at which almost two thirds are. We are not aware of a comparable quantitative census at any memory length; the qualitative expectation that longer memory multiplies the ways of stabilising cooperation is in Stewart and Plotkin (35) and Hilbe et al (25).

**Why the main text quotes no share that depends on a window.** Every share in the three paragraphs above is a share of a window, and the window is a choice the answer depends on. The shares the main text does quote are of the other kind — natural densities, equal to a fraction of the directions at infinity and so free of any window; they are §6, and nothing in this paragraph applies to them. The disk of Main eq. (2) is $(u, v) \mapsto (u, v)/\sqrt{k^2 + u^2 + v^2}$ with $k = 4$; since $\varphi_k(p) = \varphi_1(p/k)$, changing $k$ is precomposing with a dilation of the plane, and the arrangement is not dilation-invariant — $u = -4$, $v = -2$, $u + v = 1$ and most of the $254$ facet lines miss the origin. So every such share moves with $k$ exactly as it moves with a square's half-width:

| | games at which *every* equilibrium is maximally efficient | | | median efficient share |
|---|---|---|---|---|
| measure | of all games | of Snowdrift | of Harmony | PD below $u + v = 1$ |
| disk, $k = 1/2$ | $6.3\%$ | $11.7\%$ | $13.7\%$ | $89.0\%$ |
| disk, $k = 1$ | $15.8\%$ | $33.1\%$ | $30.3\%$ | $85.1\%$ |
| disk, $k = 2$ | $26.9\%$ | $63.9\%$ | $43.6\%$ | $66.3\%$ |
| disk, $k = 4$ (the figures) | $33.9\%$ | $86.8\%$ | $48.7\%$ | $31.9\%$ |
| disk, $k = 8$ | $36.5\%$ | $96.2\%$ | $49.9\%$ | $9.3\%$ |
| disk, $k = 64$ | $37.5\%$ | $99.9\%$ | $50.1\%$ | $6.4\%$ |
| square, $\lvert u\rvert, \lvert v\rvert \le 8$ | $37.5\%$ | $96.9\%$ | $53.4\%$ | $9.3\%$ |
| square, $\lvert u\rvert, \lvert v\rvert \le 64$ | $37.6\%$ | $100.0\%$ | $50.5\%$ | $6.4\%$ |
| directions at infinity | $37.5\%$ | $100\%$ | $50\%$ | $6.4\%$ |

Both families converge, and to the same place. Far from the origin an arrangement of finitely many lines is a fan of angular sectors, so the $k \to \infty$ and $M \to \infty$ limits are the same object — a fraction of the *directions* at infinity, which no parameter enters. For "every equilibrium is maximally efficient" that fraction is exactly $135°/360° = 3/8$: the whole Snowdrift sector together with the half of Harmony below $u + v = 1$. Two things follow. The square $|u|, |v| \le 8$ quoted above already sits essentially at that limit, so those percentages report the far field rather than the near-origin arrangement they appear to describe; and the main text, which states these results as the *sets* of games on which they hold, needs no window and loses nothing by doing without one.

**What "maximally efficient" forces on the pattern of play.** One of those sets can be written down exactly:

the games at which *every* equilibrium is maximally efficient. The reason is a two-line consequence of the efficiency definition above. The two players are interchangeable, so $w_2 = w_3$ and $\bar{E} = w_1 + w_2(1+u+v)$ with $w_1 + 2w_2 + w_4 = 1$. Below the switch line $E_{\max} = 1$, and $\bar{E} = E_{\max}$ reduces to $w_2\,(u+v-1) = w_4$, whose left side is non-positive and whose right side is non-negative; both therefore vanish and $w = (1,0,0,0)$. Above the line $E_{\max} = s/2$ with $s = 1+u+v > 2$, the same substitution gives $w_2 = \frac{1}{2} + w_4/(s-2)$, while $w_2 \le \frac{1}{2}$ always, so again $w_4 = 0$ and $w = (0, \frac{1}{2}, \frac{1}{2}, 0)$. *Below $u+v=1$ a strategy is maximally efficient exactly when it cooperates fully, and above it exactly when it alternates perfectly*; nothing else attains the optimum on either side. This is an identity of sets, and it was checked as one (§7).

**The set on which every equilibrium is efficient.** The share is 1 exactly at the games whose equilibrium set is contained in one of those two families. Testing that over the 23861 full-dimensional regions in exact rational arithmetic gives, up to a Lebesgue-null set of tie lines, a union of three open convex regions and nothing else in the plane,

$$\{u>1,\ v>0\} \,\cup\, \{u>0,\ v>2\} \,\cup\, \{u>\tfrac{2}{3},\ \ u-v>1,\ \ u+v<1\}. \tag{16}$$

The first two are Snowdrift with the rectangle $0 < u \le 1$, $0 < v \le 2$ removed, one corner at $(1,2)$, and there every equilibrium is an alternator. The third is a wedge of Harmony, unbounded below, with corners $(2/3, -1/3)$ and $(1,0)$; it needs no $v<0$ clause, since $u-v>1$ and $u+v<1$ force it, and there every equilibrium cooperates fully. All three require $u>0$, so no game of the Prisoner's Dilemma, of the Stag Hunt, or of Harmony above the switch line has all of its equilibria efficient. Each bounding line is a facet that readmits the other family and each is excluded, at a witness on each: on $u=1$ the count is 1568 efficient of 2276 at $(1, \frac{1}{2})$, on $v=2$ it is 832 of 860, on $u=2/3$ it is 7639 of 8239, and on $u-v=1$ — which is $S=T$, the line the pinned alternators of the tie set occupy — it is 7639 of 7985 at $(\frac{3}{4}, -\frac{1}{4})$. The single exception is $u+v=1$ along the wedge's own edge, where all 3049 equilibria are efficient at $(\frac{3}{2}, -\frac{1}{2})$; that count is not constant along the edge, but every equilibrium on it is efficient.

**Two ways to get that region wrong, both of which we took first.** (16) is a statement about generic games; deciding it at a given game needs all three strata, and reading it off the full-dimensional regions alone inverts it: at $(3, \frac{1}{2})$, which lies on the tie line $u-4v=1$, every one of the 1216 full-dimensional equilibria is an alternator and so efficient above the switch line, while the true count is 1224, and the eight the open cells miss are one-dimensional regions at $\gamma = 4/5$, efficient on neither side of the line — so the game falls outside (16) for exactly the reason a dimension filter cannot see. The authority everywhere is the whole-plane polyhedron of §7, built from the complete constraint set with no bounding box. And (16) is not monotone in $u$ at fixed $v$ — its Snowdrift part begins at $u=1$ for $0<v\le 2$ but at $u=0$ for $v>2$ — so a bisection in $u$, the obvious way to find a left edge, returns edges that are not there. A grid is safe only if it avoids the facet lines: with a single offset added to both coordinates, $u - v$ keeps the grid spacing and every fourth sample lands exactly on $u - v = 1$, where floating point calls the line interior and the exact count does not.

**Which games keep an efficient equilibrium at a fixed error rate.** The same lemma answers the fixed-$\epsilon$ question, and the answer is a wedge. Below the switch line the question is whether any of the 7639 fully cooperating strategies survives: only 245 of them hold a two-dimensional region at $\epsilon = 10^{-4}$, and the same number at $10^{-3}$, $10^{-2}$ and $10^{-1}$, and the union of those regions has exactly *one* facet below the line — contributed by the single resident 36873, which is $AON_2$, and whose limit region is the $\{S \le R\} \cap \{T \le 3R\}$ of §5 above. The boundary of $W_\epsilon$ is therefore the $AON_2$ facet that carries the memory-2 cost threshold, tilted by $\epsilon$. Above the switch line the question is whether any of the 3072 perfect alternators survives, and at $\epsilon = 10^{-4}$ exactly 134 of them are Nash at some game, 26 on a line and 108 at a single game, and none on a region. So the games that keep an efficient equilibrium are, up to a Lebesgue-null set and at the three smaller reported rates $\epsilon = 10^{-4}, 10^{-3}, 10^{-2}$, exactly the wedge

$$W_\epsilon \;=\; \{\, u + v < 1 \,\} \;\cap\; \Big\{\, v - \tfrac{\epsilon}{1-\epsilon}\, u \;\le\; \kappa(\epsilon) \Big\}, \qquad \kappa(\epsilon) = 2 - 8\epsilon + O(\epsilon^2), \tag{17}$$

with $\kappa = 1.99920010$, $1.99200999$ and $1.92099402$ at those three rates. It is the limit facet $T \le 3R$ tilted by $\epsilon$ and slid inward by $8\epsilon$. At $\epsilon = 10^{-1}$, where $\kappa = 1.29422222$, the wedge (17) is an outer description only: the union there carries exactly one further facet, found where the check against the raw sixteen half-planes — 20256 games spanning ten decades below the switch line, with no disagreement at the three smaller rates — returns its two disagreements.

**The three numbers of Main Figure 3b, which are easy to confuse.** Two of them coincide, and the third is a raster artefact. 245 is both the largest number of efficient equilibria at any *single game*, certified over the whole plane, and the number of fully cooperating *strategies* that hold a two-dimensional region anywhere in the plane: at $\epsilon = 10^{-4}$ these are the same 245 strategies, simultaneously Nash at every game of the maximising set. That set is not a finite list of vertices but an unbounded region of positive area, whose nearest point is the corner $|u| \approx 2.00008 \times 10^7$, the meet of genotype 28663's deviation 14 with genotype 6015's deviation 7. No nearest or farthest vertex of that region may be quoted from the kit that found it (§7). 244 is what a region file clipped to $|u|, |v| \le 2^{20}$ resolves: it misses genotype 6015, whose region lies at $|u| \sim 10^{23}$ (§7). Evaluated against the raw sixteen half-planes, which are the exact condition everywhere, 6015 is a genuine equilibrium there. 222 is the largest value the raster attains at a pixel centre, and it falls short of 245 because the maximising games are $10^7$ or more from the origin, where the disk carries almost no pixels — the same shortfall as Main Figure 2b's 968 against its certified 1024.

**Where exactly one efficient equilibrium survives, and which one.** *ALLC*. Its fixed-$\epsilon$ region is $v \le \epsilon u/(1-\epsilon)$, which contains the whole of Harmony; every other fully cooperating strategy carries a facet at $u \le 1$ or tighter — $AON_2$'s region is $u \le 1$, $v \le 2$ and *WSLS*'s is $u \le 1$, $v \le 1$ — so beyond $u = 1$ they have all dropped out. Hence at every game with $u > 1$ and $u + v < 1$ the efficient equilibrium set is exactly $\{ALLC\}$. Below $u = 1$ the count falls to one as well, but only much further out, the $\epsilon$-tilt of the other regions' facets deciding where — at $u = 0.99$ not until $v \approx -10^2$, at $u = 0.1$ not until $v \approx -10^4$. And it does not stop at the $v$ axis: it continues into the Stag Hunt as a wedge hugging that axis, reaching $u = -9$ at $v = -10^5$, $u = -99$ at $v = -10^6$ and $u = -999$ at $v = -10^7$ — a reach of order $\epsilon|v|$, dying out below $|v| \approx 10^4$ and subtending some $6 \times 10^{-3}$ degrees at the rim, which is why Main Figure 3b shows the region as Harmony alone.

**At a generic Prisoner's Dilemma the count is at least four.** Scanning ten decades in each coordinate, the non-zero counts observed there are 4, 5, 6, 7, 11, 12, 15, 16, 19 and 22: at a generic game, where the Prisoner's Dilemma keeps an efficient equilibrium at all it keeps at least four. A scan cannot see the lines and single games, and there it can keep exactly one: at $(u, v) \approx (-9998, 0.9994)$, an endpoint of 3927's one-dimensional efficient region, that strategy is the only one. *ALLC* is never among them, and that is the reason — its region is $v \leq \epsilon u/(1-\epsilon)$, which for $u < 0$ requires $v < 0$, so the quadrant is excluded outright. What survives instead is the $AON_2$ family: the smallest non-zero set is the four residents 32777, 32783, 36873 and 36879, which differ only in the row for a most recent $CC$ and in the component $(DD, CC)$, and of which 36873 is $AON_2$ itself. Snowdrift is more extreme again at a generic game, never keeping fewer than ten there, though not near the corner $(1, 0)$: at $(u, v) = (9999/10000, 10^{-6})$ the only efficient equilibrium is *ALLC*.

**What that one half-plane accounts for, and what it does not.** For $u < 0$ the term $-\epsilon u/(1-\epsilon)$ is positive and grows with $|u|$, and the consequences are immediate. The Stag Hunt keeps an efficient equilibrium until $v + \epsilon|u|/(1-\epsilon)$ exceeds $\kappa$, that is until $|u| \sim 2/\epsilon$: a wedge along $v = 0$ opens at $|(u, v)| \approx 2 \times 10^4$ at $\epsilon = 10^{-4}$, and inside it equilibria survive of which, off a set of measure zero, none is efficient. The count is not constant along it — twenty at $(-10^5, -5)$, twenty-four at $(u, v) \approx (-2 \times 10^7, -1/5)$ where 5503 alone is efficient, and one, *ALLD*, at $(-10^{12}, -5 \times 10^7)$. The Prisoner's Dilemma keeps at least one near the origin and none in the far field, for the same reason with $v$ helping. Harmony below the switch line keeps at least one at *every* game, since $v < 0$ and $-\epsilon u/(1-\epsilon) < 0$ both make the left-hand side negative. Harmony above the line keeps none off a set of measure zero, failing the first half-plane outright; on it, three alternators hold rays running out into that half of Harmony. And the donation game at cost 8/9 is the point $(u, v) = (-8, 8)$: it satisfies $u + v = 0 < 1$ but gives $v - \epsilon u/(1-\epsilon) = 8.0008$ against $\kappa = 1.9992$, so it carries no efficient equilibrium — predicted by (17) rather than found.

**Two effects, at two scales.** That half-plane is an *existence* statement, and it does not account for the collapse in counts. A positive error rate acts on the equilibrium set in two ways, and only the second is a far-field phenomenon. First, the ties resolve. Every equilibrium of the limit game is weak (§4), and the constraints holding with equality there are the ones whose coefficients $(\alpha, \beta, \delta)$ vanish together as $\epsilon \to 0$. Such a constraint reads $0 \leq 0$ in the limit; at $\epsilon > 0$, divided through by its own scale, it is an honest half-plane whose direction is $O(1)$. It therefore does not displace a boundary by $O(\epsilon)$ — it cuts the region down, or removes it altogether, at every scale including the origin. Of the 23861 strategies holding a two-dimensional region in the limit, 21700 are Nash at no game whatever at $\epsilon = 10^{-4}$; genotype 24207, whose limit region is the half-plane $u + v \leq 1$, has empty fixed-$\epsilon$ region at every $\epsilon$ measured. At $(u, v) = (-8, -8)$, 21424 of the 22069 limit equilibria are lost, each of them at least 4 units of game space inside its own limit boundary, and that count is unchanged across $\epsilon = 10^{-2}$ to $10^{-4}$. Second, the boundaries that survive move. A constraint with a nonzero limit triple keeps its half-plane and acquires an $O(\epsilon)$ rotation and offset: *ALLC*'s $v \leq 0$ becomes $v \leq \epsilon u/(1-\epsilon)$, and *AON*$_2$'s $v \leq 2$ becomes $v - \epsilon u/(1-\epsilon) \leq \kappa(\epsilon)$. A rotation of order $\epsilon$ competes with an $O(1)$ intercept only once $|(u, v)|$ approaches $1/\epsilon$, which is why the Stag Hunt wedge opens at $|u| = \kappa(\epsilon)(1-\epsilon)/\epsilon$ — measured at 19990.00, 1990.02, 190.18 and 11.65 for $\epsilon = 10^{-4}$, $10^{-3}$, $10^{-2}$ and $10^{-1}$. The first effect sets which strategies are in play; the second sets where their boundaries lie.

**What the census cannot settle.** The equilibrium notion has therefore stopped discriminating. It is not that efficient outcomes are unavailable, nor that inefficient ones are excluded: both are available at essentially every game, in numbers, and which of them a population actually reaches is a question about the evolutionary dynamics (130) and not about the equilibrium set. That conclusion is not new — it is the standard reading of the Folk theorem (10, 11), stated for direct reciprocity in almost these words by Tkadlec et al (78) and argued in detail by Fudenberg and Maskin and García and van Veelen (3, 58, 79) — but it is here measured rather than asserted, and the measurement is what makes it the strongest argument the equilibrium analysis can offer for an evolutionary one (130).

**Friendly rivals: the one class the census does single out.** A strategy is a *rival* if no co-player ever earns more than it does in their interaction, $\pi(\sigma, \tau) \geq \pi(\tau, \sigma)$ for every $\tau$, read in the limit $\epsilon \to 0$ (45, 72); a *friendly rival* is a strategy that is efficient and a rival (26). With $w$ the limiting pair distribution seen from $\sigma$'s side, $\pi(\sigma, \tau) - \pi(\tau, \sigma) = (T - S)\,(w_{DC} - w_{CD}) = (1 + v - u)\,(w_{DC} - w_{CD})$, so rivalry depends on the game only through the sign of $u - v - 1$, exactly as efficiency depends only on the sign of $u + v - 1$. Three consequences. (i) An efficient rival is a Nash equilibrium at every game, since $\pi(\tau, \sigma) \leq \frac{1}{2}[\pi(\tau, \sigma) + \pi(\sigma, \tau)] \leq E_{\max} = \pi(\sigma, \sigma)$; friendly rivals are therefore partners, and a co-player who ties them must have let them earn $E_{\max}$ too, so they satisfy the tie clause of Akin (44) as well. (ii) The set of friendly rivals is constant on each of the four open quarter-planes cut out by $u + v = 1$ and $u - v = 1$, which meet at $(1, 0)$. Testing all 65536 residents against all 65536 co-players, 2640 strategies are rivals for $T > S$ and, by the $C \leftrightarrow D$ conjugation (Main Methods), the same number for $T < S$; intersected with the efficient sets this gives 8 friendly rivals on $\{u + v < 1,\ u - v < 1\}$, 1519 on $\{u + v < 1,\ u - v > 1\}$ and 80 on each of the two quarter-planes above the switch line, the latter two conjugate to each other. On the switch line itself the class is larger, 116 on the $T > S$ side and 2067 on the $T < S$ side, because some of the 4046 strategies that are efficient only on that line join it; on $T = S$ every strategy is a rival, so there the friendly rivals are simply the efficient strategies. (iii) The eight of the Prisoner's Dilemma quarter-plane are, by genotype, 19079, 19087, 20103, 20111, 23175, 23183, 24199 and 24207. 23183 is *TFT-ATFT* (27), the first friendly rival exhibited under implementation errors, and 20111 and 24207 — the two strategies whose equilibrium region is exactly the half-plane $u + v \leq 1$ (§5.1) — are the two of the eight that are rivals for either sign of $T - S$. Each of the eight is a partner at every game of the quarter-plane by (i), and Main Figure 3a counts exactly 8 partners on the wedge $v > 2$, $2 - 2v < u < 1 - v$ where it attains its minimum, so the eight *are* that minimum: in the hardest Prisoner's Dilemmas the only partners left are the friendly rivals. The 1519 of the $T < S$ quarter-plane are a cheaper class than the name suggests: they are the conjugates of the 1519 fully defecting rivals, and when $T < S$ "never outperformed on the asymmetric exchange" is satisfied by being exploited, so the class contains *ALLC*. Murase and Baek's original notion is the error-free *defensibility*: no co-player can drive the cumulative payoff difference to $-\infty$ from any state at $\epsilon = 0$, which for a deterministic strategy is the absence of a negative cycle on its sixteen states (26). Under it 2144 of the 2640 rivals qualify and the four counts become 8, 1036, 80, 80: the eight and the alternators are the same under either reading. The rivalry test is a sign test on the same limiting pair distributions as the census. An independent 95-line program, `pairs.c` in the repository's `friendlyrivals` folder, recomputes those limits by GTH state reduction in leading-order arithmetic and reproduces every count of this paragraph in about a minute on twelve cores; the smallest nonzero $|w_{DC} - w_{CD}|$ over all $2640 \times 65536$ rival pairs is $7.0 \times 10^{-3}$, so no sign decision is close.

##### 5.3 $\Lambda$, the tie set, and the alternation corner

**$\Lambda$, the tie set: measure-zero equilibria, found by construction.** Let $\Lambda$ denote the union of the one-dimensional regions. These sets are Lebesgue-null and therefore invisible in any raster of the plane, and a lattice of games misses them with probability one — which is exactly why they must be constructed rather than sampled, and why the representative-point coverage that is standard in whole-plane studies (7, 15, 16) cannot reach them. A region of (15) is one-dimensional precisely when two of the half-planes are *exactly antiparallel*: a constraint together with its negative forces equality and pins the resident to a line. Because the coefficients are exact rationals of §7, that condition is an identity between integers, and the line comes out as a primitive integer relation in $(u, v)$. That exactly antiparallel pairs should exist at all is intelligible from the zero-determinant theory: a memory-2 strategy can enforce a linear relation between the players' payoffs (31, 43, 65), and a linear relation enforced in both directions is a pair of opposite half-planes.

**Forty-four lines.** The 3774 one-dimensional regions lie on 44 distinct lines. The striking feature is a rigidity that nothing in the construction imposes: 31 *of the* 44 *lines carry strategies of one single level of cooperation*. The line $3u - v = 1$, that is $T = 3S$, carries 72 strategies and every one has $\gamma = 1/4$; $u - 3v = 1$ carries 74, all at $\gamma = 3/4$; $u - 2v = 1$ carries 61, all at $2/3$; $2u - v = 1$ carries 26, all at $1/3$; and $u - v = 1$, that is $S = T$, carries 1280, all at $\gamma = 1/2$. The two most populated lines together hold 72.5% of the 3774 strategies, and the larger of the two, $u + v = 1$, is much the biggest exception to the pattern: it carries eight distinct levels, which is unsurprising since it is the line on which the social optimum itself changes character. The single-level lines are a majority of the lines but a minority of the strategies: the 31 of them carry 1819, 48.2% of $\Lambda$, and 1486 of those — 81.7% — are at $\gamma = 1/2$, spread over nine of the 31. Neither extreme level ever pins a line — no line has $\gamma = 1$ or $\gamma = 0$ as its unique level, consistent with the census, since those are the largest and the third-largest full-dimensional families and a strategy that holds an open set of games is not on a tie line at all. A rule predicting which level goes with which line, or the converse, is the natural next question and is not yet answered. We have found no published construction of this kind — a finite list of degenerate lines of games, each labelled by the cooperation level of the strategies it supports — at any memory length; the general fact that the degenerate games form a null set (98) is of course classical, and the one-shot analogue is the line arrangement of Falmagne (107).

**The alternation corner is exactly partitioned.** The 3072 strategies whose limiting self-play distribution is $(0, \frac{1}{2}, \frac{1}{2}, 0)$ — perfect alternation, $\gamma = 1/2$, and maximally efficient wherever alternation is the social optimum — divide with nothing left over:

$$3072 = \underbrace{1792}_{\text{full-dimensional } Z(p)} + \underbrace{1280}_{\text{pinned to } u-v=1} . \tag{18}$$

Every alternator either holds an open set of games or sits exactly on $S = T$, and the reason is transparent: on $S = T$ neither role of the alternation pays more than the other, so no deviator can profit by seizing one of them, while off that line the two roles differ and only those alternators that punish the seizure survive. The two problems that make the difference — who takes the favourable role, and how the seizure is deterred — are the two that the turn-taking literature identifies as the substance of a turn-taking equilibrium (84, 86–88). The line $S = T$ has been flagged once before as distinguished for memory-2 alternators, and over this same space of $2^{16}$ strategies (§1.6): Wang et al (83) find that two-step-memory turn-taking emerges when $S + T > 2R$ and $S \neq T$, and that on $S = T$ a non-alternating strategy prevails instead. That is a simulation result without a mechanism, and it points the other way — alternation leaves the line there, and is pinned to it here. (18) is, we believe, the first equilibrium-theoretic account of why the line is special. The levels of the 23861 full-dimensional regions are led by $\gamma = 1$ with 7639 strategies, then $\gamma = 1/2$ with 6736, $\gamma = 0$ with 4990, 2/3 with 2444 and 1/3 with 2052.

**The most equilibria any game carries are carried on a region, not on a line.** The range 299 to 22069 of Main Figure 2a counts full-dimensional regions, so it is exact at every game off the arrangement and silent on the lines themselves, where a pinned strategy is an equilibrium and nowhere else. Those lines must be tested separately, and doing so is delicate in a way §7 sets out. Its facet pair bounds the whole line, and a membership test built from facets alone therefore accepts the entire line rather than the interval; the endpoints have to be carried explicitly. The lines do carry more than their neighbours — on $u - v = 1$ at $u = 1$ the count is 14757, of which 2966 are one-dimensional regions and 2360 are single-point regions at the degenerate game $(1, 0)$, against a flat 3072 at every $u > 1$ on the same line — but, as it turns out, never the most.

The maximum over the *whole* plane can be settled exactly, and the argument is short. Every $Z(p)$ of (15) is a closed polyhedron whose facets lie on the 280 lines that occur in the construction — the 254 facet lines together with the 26 of the 44 tie lines that bound no full-dimensional region — so every $Z(p)$ is a union of closed cells of that arrangement, refined by the finitely many interval endpoints and single-point regions. If a point lies in the relative interior of a cell contained in $Z(p)$, so does the closure of that cell; the count can therefore only rise as one passes to a lower-dimensional stratum, and the maximum must be attained at a vertex. Every cell's closure holds one, unbounded cells included, so the far field needs no separate treatment. Evaluating all 22127 vertices and 49814 edges in exact integer arithmetic, with no tolerance anywhere (§7), gives

$$\max_{(u,v)\in\mathbb{R}^2} \#\{p : (u, v) \in Z(p)\} \;=\; 22069, \tag{19}$$

the Stag Hunt figure with nothing added to it: not one of the 22069 is a tie-line or a single-point region, so $\Lambda$ contributes nothing to the extremum. The set of games attaining it is the closed region

$$\{(u, v) \,:\, u \leq -4,\ v \leq -2\}, \qquad \text{that is} \qquad S \leq -4 \text{ and } T \leq -1 \text{ at } R = 1,\ P = 0, \tag{20}$$

and nothing else in the plane; it is the deep plateau of Main Figure 2a. It is unbounded, it has positive area — $9.375\%$ of the square $[-8, 8]^2$, $22.7\%$ of the $[-64, 64]^2$ the construction was carried out in, and $5.16\%$ of the disk Main Figure 2 draws, which compactifies the whole plane rather than showing a window of it — and it has a single corner, at $(u, v) = (-4, -2)$. The *same* 22069 strategies are equilibria at every one of its games, so the region is exactly the intersection of their 22069 polyhedra; it is not one cell of the arrangement but a union of many, with 31 arrangement vertices in it, 3 of them interior, and the count is constant across all of them and out to infinity within it. What fixes each of its two boundaries is a count of facets: exactly 3 of the 23861 full-dimensional regions carry the facet $u \leq -4$ and exactly 1249 carry $v \leq -2$, so the count falls to 22066 on crossing the first and to 20820 on crossing the second. So the games with the most Nash equilibria anywhere in the plane are the Stag Hunts in which defecting against a cooperator pays less than mutual defection and being defected against while cooperating costs at least four $R - P$ below $P$ — and they are an ordinary open set of games rather than an exceptional one.

The same monotonicity settles the other extreme. Since the count can only rise as one descends a stratum, the minimum is attained on an open set and not on its boundary: it is the 299 of Main Figure 2a, held on an open triangle of area $1/24$, on each of whose three bounding lines the count is strictly larger — 387 on $u + v = 1$, 421 on $v = 2$ and 301 on $4u + v = -3$. So both extremes of the equilibrium count are attained on open sets of games, and a sufficiently fine grid finds both. What $\Lambda$ costs a representative-point study is therefore not the extremes but the strata: the 3774 strategies whose entire equilibrium set is a line, and the 4875 whose entire equilibrium set is a single game — 8649 in all, more than one strategy in eight — are invisible to any scheme that samples games, however fine, and they are what Main Figures 2**c** and 2**e** exist to show.

**At a positive error rate the maximum retreats to the far field.** The argument above is a limit argument, and its two halves survive the perturbation unequally. The monotonicity is unaffected — every $Z_\epsilon(p)$ is a closed polyhedron, so the count still rises only as one descends a stratum — but the arrangement is not the same object. Its facet lines must be recomputed for each $\epsilon$ over the whole plane: at $\epsilon = 10^{-4}$ there are 1522 of them, against the 254 of the limit, and the count of $\Lambda$-carried strategies falls from 3774 on 44 clean lines to 564 scattered over 492 separate ones — so the arrangement is cut by 2008 lines in all where the limit needs 280. Taking the recession cone of each region, which decides how far out it reaches without enumerating the arrangement at all, gives a maximum of 1024, the same at every one of $\epsilon = 10^{-1}$, $10^{-2}$, $10^{-3}$ and $10^{-4}$, against the 22069 of (19). It is exactly *twenty-six* short of the number of full-dimensional regions at that $\epsilon$ — 1050 at every one of them — so at those games all but twenty-six of the strategies that hold a full-dimensional region are equilibria *at once*, and the twenty-six that cannot join are the same twenty-six at every $\epsilon$: seventeen at the self-play point $(\frac{1}{2}, \frac{1}{4}, \frac{1}{4}, 0)$, five at $(0, \frac{1}{4}, \frac{1}{4}, \frac{1}{2})$ and four at $(\frac{1}{4}, \frac{1}{4}, \frac{1}{4}, \frac{1}{4})$, which is *TFT*'s, the only full-dimensional regions in this space whose self-play point is neither mutual cooperation nor the alternation corner and still carries weight on $S$ and $T$. And the maximum sits where the limit's does not. The games attaining it are an intersection of the 1024 regions that achieve it, hence convex, and that set is *full-dimensional*: a strictly interior point can be exhibited, and its directions at infinity fill 88.8°, 89.9° and 90.0° of the plane as $\epsilon$ falls — one quadrant, and that quadrant is the Stag Hunt whose flat asymptotic count Main Figure 2a already records. But it begins far out, where the limit's maximising region reaches in to $S = -4$, $T = -1$. Not one of the achieving regions is one-dimensional, at any $\epsilon$ or in the limit: the tie set never carries the maximum, at either notion of equilibrium. So the extremum is an open set of games at both notions of equilibrium, and what the error rate changes is where one has to look: the limit's maximising region is found by any grid fine enough to resolve an area of 9.375% of $[-8, 8]^2$, while at each of the three error rates measured — $\epsilon = 10^{-2}$, $10^{-3}$ and $10^{-4}$ — the maximising region begins beyond any window a figure of this kind draws, and the $4000^2$ raster of Main Figure 2b understates it by 56 — 968 against 1024 — for want of pixels that far out. A fixed-$\epsilon$ study must sample far; a limit study need only sample finely.

#### 5.4 Memory-one, for contrast

**Memory-1, for contrast, and it is a different picture entirely.** The same census over the 16 binary memory-1 strategies is exact and symbolic in $\epsilon$, so none of the numerical caveats of §7 apply to it. The deviation set is all 16 strategies, so comparing against every one of them *is* the full best-reply test and the difficulty of §3 does not arise. SI Figure 4 draws all of what follows. Of the 16, *eight* hold a full-dimensional region of games, *four* are equilibria only along a line, *three* only at a single game — 1100 and 0011 at the same game $(0, 0)$, and *Anti TFT* at $(\frac{1}{2}, -\frac{1}{2})$ — and one — *TFT* — is an equilibrium at *no game whatever*: its constraint set contains both $u + v \leq 0$ and $u + v \geq 2$ and is therefore infeasible, which no single constraint of it reveals. *Anti TFT* is the one of the pair that survives, and only at that single game. The arrangement has 20 distinct facet lines against the 254 of memory-2, and a game carries between 0 and 8 equilibria against the 299 to 22069 memory-2 carries at a generic game.

*TFT* is worth one more sentence, because it is this document's own distinction in miniature. Recomputing the same regions at $\epsilon = 10^{-2}$, $10^{-3}$ and $10^{-4}$ leaves every one of the sixteen dimensions unchanged *except TFT*'s, which is non-empty at all three — and is a *single game*, exactly one, at each: $(u, v) = (-49, 49)$, $(-499, 499)$ and $(-4999, 4999)$. All three lie on $u+v = 0$, so *TFT* is an equilibrium at exactly one donation game at each error rate, at cost $c = 1 - 2\epsilon$. That is the mechanism of the discontinuity rather than a symptom of it: the game is real at every positive $\epsilon$ and recedes like $1/2\epsilon$ as $\epsilon$ falls, so the resident that is an equilibrium somewhere at every error rate we measured is an equilibrium nowhere in the limit. The direction of that failure is the opposite of the one §3 warns about — there the limit *over*-accepts, here it rejects outright — and both have the same root, that the $\epsilon = 0$ problem is not the limit of the $\epsilon > 0$ problems (49).

The zero is the substantive difference. *ALLD*'s region is exactly $\{u \leq 0\}$ and *ALLC*'s exactly $\{v \leq 0\}$, so between them they cover three quadrants, and in $u, v > 0$ the only strategy holding an open set is *WSLS*, whose region $\{u \leq 1\} \cap \{v \leq 1\} \cap \{u + v \leq 2\} \cap \{u + 2v \leq 3\}$ meets that quadrant in exactly the unit square: the last two constraints are implied by the first two once $u, v > 0$, so the island is $[0, 1]^2$ and its area is exactly 1. Three one-dimensional regions reach into Snowdrift as well — genotypes 2 and 11 along $v = 3u - 1$ and $3v = u - 1$, both running to infinity, and genotype 13 bounded within Snowdrift — and they carry no area, so off those three lines and outside *WSLS*'s square every Snowdrift game has *no* binary memory-1 equilibrium at all — 25.000% of the box $|u|, |v| \leq 2^{20}$ the memory-1 kit uses (which is 25% minus exactly that one square unit, so the figure is a rounding and not an identity) and 23.16% of the disk as drawn — where memory-2 has at least 299 everywhere. The two unbounded rays run in the directions $v = 3u$ and $v = u/3$, which are the two the memory-2 slivers of (26) tend to as $\epsilon \to 0$. The structure is not an artefact of the limit (SI Figure 4 draws the limit and $\epsilon = 10^{-4}$): at $\epsilon = 10^{-2}$, $10^{-3}$ and $10^{-4}$ the same eight strategies hold full-dimensional regions and every region keeps its dimension *except TFT*'s, as above, the empty set staying at 22.94%, 23.14% and 23.16% of the disk. What moves is the boundaries, by $O(\epsilon)$, and the facet count, which nearly triples to 52 as ties that coincided in the limit separate.

**Above $u + v = 1$ memory-1 cannot be efficient, and the ladder is the whole reason.** SI Figure 5 draws it: above that line the page is grey at every game, dark where equilibria survive without an efficient one and light where none survives at all, the part of Snowdrift beyond *WSLS*'s island being the light kind, as at $(5, 5)$. Write $s = u + v$. In self play $w_2 = w_3$ by (10), so a resident at $w$ earns $E = w_1 + w_3 + w_3\, s$ — affine in $s$ with slope $w_3$ — while for $s > 1$ the attainable optimum is $E_{\max} = (1+s)/2$, affine with slope $\frac{1}{2}$. Attaining it therefore needs $(w_3 - \frac{1}{2})\, s = \frac{1}{2} - w_1 - w_3$. The largest $w_3$ anywhere in the memory-1 space is $\frac{1}{4}$, so the slope is short of $\frac{1}{2}$ at every one of the six points and the two lines meet at most once: at $s = -1$ for $(0,0,0,1)$ and $(0, \frac{1}{4}, \frac{1}{4}, \frac{1}{2})$, at $s = 0$ for $(\frac{1}{4}, \frac{1}{4}, \frac{1}{4}, \frac{1}{4})$ and $(\frac{1}{2}, 0, 0, \frac{1}{2})$, and at $s = 1$ exactly — the switch line itself — for $(1,0,0,0)$ and $(\frac{1}{2}, \frac{1}{4}, \frac{1}{4}, 0)$. *No memory-1 point meets $E_{\max}$ at any $s > 1$*, so memory-1 has no efficient strategy above the switch line, let alone an efficient equilibrium, at any error rate. The equality case is what memory-2 supplies and memory-1 cannot: slope $w_3 = \frac{1}{2}$ with $w_1 = 0$ is the alternation corner $(0, \frac{1}{2}, \frac{1}{2}, 0)$, which no binary memory-1 strategy reaches — its largest alternation mass $w_2 + w_3$ is $\frac{1}{2}$ against the corner's 1 — and on which memory-2 puts 3072 strategies exactly (§5 above).

What does *not* follow, and it is worth saying because it is the tempting summary: memory-1 equilibria are not confined to the mutual-cooperation payoff. All four line regions run to infinity, and on two of them the payoff is unbounded — 0100 is an equilibrium exactly on the ray $\{v = 3u - 1,\ u \geq 0\}$, where it earns $E = u$, and 1101 exactly on $\{v = (u-1)/3,\ u \geq 1\}$, where it earns $(2+u)/3$. What collapses is not the payoff but the *ratio*: on the first ray $E/E_{\max}$ is pinned at exactly $\frac{1}{2}$ however far out one goes, and on the second it decays from 1 at $u = 1$ through $4/5$, $4/7$ and $34/67$ at $u = 2, 10, 100$ towards $\frac{1}{2}$. On the other two rays the payoff does not grow at all, and the formula above says why: $E$ depends on the game only through $s = u + v$, and those two rays pin it — 0010 is an equilibrium exactly on $\{u + v = -1,\ u \leq -1\}$ and 1011 exactly on $\{u + v = 1,\ u \geq 0\}$, the switch line itself — so $E$ is constant along each of them however far out one goes. So the right contrast is in efficiency and not in payoff: memory-1 equilibria can earn arbitrarily much and still capture only half of what the game allows, while above the switch line memory-2 equilibria capture all of it.

#### 6 The four kinds of game, and how much of the plane each one takes

Everything in this section is a statement about $0 < \epsilon < \frac{1}{2}$, which is the whole range in which the strategy space exists: at $\epsilon = \frac{1}{2}$ every component of every strategy equals $\frac{1}{2}$ and all 65536 collapse to one object. The endpoint is treated at the close.

**A share of the plane needs a definition before it can be a number.** The plane of games is unbounded, so "the fraction of games such that ..." depends on the window one measures in, and every window gives a different answer: the share of games at which every equilibrium is maximally efficient runs from 6.3% to 37.5% as the disk parameter $k$ of Main eq. (2) goes from $\frac{1}{2}$ to infinity (§5). We therefore use the natural density

$$\mu(X) \;=\; \lim_{r\to\infty} \frac{\left|X \cap rB\right|}{|rB|}, \qquad B \text{ the unit disk.} \tag{21}$$

Every set considered here is a finite Boolean combination of half-planes, so its asymptotic cone exists, the limit (21) exists with it, and it equals the normalised angular measure of that cone: a fraction of the *directions* *at infinity*, free of any window. The tool is the recession cone. A convex region $Z$ runs arbitrarily far out in the direction $w$ exactly when every one of its facet normals $(\beta, \delta)$ satisfies $(\beta, \delta) \cdot w \leq 0$, so the directions that keep an equilibrium are a finite union of closed cones, computable in integer arithmetic from the exact half-planes of §3. Write throughout

$$\tau \;=\; \arctan \frac{\epsilon}{1-\epsilon}, \tag{22}$$

the angle by which a positive error rate tilts every facet of the arrangement off the axes.

**Two regions that can be written down by hand, and they are the extreme ones.** Let the resident be *ALLD*. Its intended action is $D$ in every state, so its realised action is $D$ with probability $1-\epsilon$ *independently of the history*. A deviator $\mathbf{q}$ with stationary cooperation rate $c_{\mathbf{q}}$ therefore meets the four outcome frequencies $\big(c_{\mathbf{q}}\epsilon,\ c_{\mathbf{q}}(1-\epsilon),\ (1-c_{\mathbf{q}})\epsilon,\ (1-c_{\mathbf{q}})(1-\epsilon)\big)$ for $CC, CD, DC, DD$, and

$$E(\mathbf{q}, \mathit{ALLD}) \;-\; E(\mathit{ALLD}, \mathit{ALLD}) \;=\; \big(c_{\mathbf{q}} - \epsilon\big)\,\big[(1-\epsilon)u - \epsilon v\big]. \tag{23}$$

At $\epsilon > 0$ the pair chain is irreducible on all sixteen states, so $c_{\mathbf{q}} > \epsilon$ for every $\mathbf{q} \neq \mathit{ALLD}$ and the bracket alone decides. The same computation for *ALLC* carries the factor $c_{\mathbf{q}} - (1-\epsilon) \leq 0$. Both factors are non-zero exactly when $\epsilon < \frac{1}{2}$, since $\epsilon \leq c_{\mathbf{q}} \leq 1-\epsilon$; this is where the hypothesis $\epsilon < \frac{1}{2}$ enters, and it is the only place it does. Hence, for $0 < \epsilon < \frac{1}{2}$, exactly and at every game in the plane,

$$Z_\epsilon(\mathit{ALLD}) = \{(1-\epsilon)u \leq \epsilon v\}, \qquad Z_\epsilon(\mathit{ALLC}) = \{(1-\epsilon)v \leq \epsilon u\}, \tag{24}$$

two half-planes through the origin, each the reflection of the other in $u = v$, each tilted off its axis by exactly $\tau$. All sixteen single-component deviations give the one half-plane; the varying factor in (23) cancels.

**Every game outside one cone keeps an equilibrium.** The complement of the union of the two half-planes (24) is the open cone

$$K_\epsilon \;=\; \big\{(1-\epsilon)v > \epsilon u\big\} \;\cap\; \big\{(1-\epsilon)u > \epsilon v\big\}, \tag{25}$$

which lies inside the Snowdrift quadrant $u, v > 0$ and is that quadrant with a wedge of angle $\tau$ removed along each axis: opening $\pi/2 - 2\tau$, apex at the *origin*. So the games with no equilibrium at all are contained in $K_\epsilon$, exactly rather than asymptotically. Because a cone through the origin is carried by the map of Main eq. (2) to a circular sector of the same angle at every $k$, the containing cone has the same share in the plane and in the drawn disk at every $k$. In a centred square it does not: a cone of opening $\pi/2 - 2\tau$ takes $(1 - \tan\tau)/4$ of any centred square, against the density $\frac{1}{4} - \tau/\pi$, and the two differ at first order in $\tau$. The set it contains does not: a neighbourhood of the origin inside $K_\epsilon$ still carries equilibria, so $K_\epsilon$ is a bound on any bounded window and an equality only in the density (21).

**The cone is not the whole answer: four slivers, not two.** Two mirror-image slivers inside $K_\epsilon$ do keep an equilibrium arbitrarily far out, at every error rate in $(0, \frac{1}{2})$, held by six residents: 62143, 48059, 61695 and their images 688, 240, 8738 under the conjugation of **Methods** — relabelling $C$ and $D$ and exchanging the two roles — which acts on the plane as $u \leftrightarrow v$. In each triple the first holds a two-dimensional region and the other two hold lines. For $0 < \epsilon \leq \frac{1}{4}$ these six are the whole story inside $K_\epsilon$, and that they are is certified over the entire interval rather than sampled; above $\frac{1}{4}$ four further residents intrude, and the paragraph after next is theirs. Both edges of the sliver are exact cubics in $\epsilon$,

$$\big(6 - 19\epsilon + 24\epsilon^2 - 12\epsilon^3,\ \ 2 - 5\epsilon + 8\epsilon^2 - 4\epsilon^3\big) \quad \text{and} \quad \big(6 - 15\epsilon + 12\epsilon^2 - 4\epsilon^3,\ \ 2 - \epsilon - 4\epsilon^2 + 4\epsilon^3\big), \tag{26}$$

so the sliver width is $\sigma = \arctan(\Xi/\Theta)$ with $\Xi = 8\epsilon\,(1-\epsilon)(1-2\epsilon)^2(2-3\epsilon+2\epsilon^2)$ and $\Theta = 2\,(20 - 108\epsilon + 257\epsilon^2 - 336\epsilon^3 + 248\epsilon^4 - 96\epsilon^5 + 16\epsilon^6)$. As $\epsilon \to 0$ the two slivers tend to the directions $v = u/3$ and $v = 3u$, each of width $\frac{2}{5}\tau$. Hence, with $\varrho$ the third angle defined in the next paragraph — identically zero on $(0, \frac{1}{4}]$ and positive beyond —

$$\mu\{\text{no Nash equilibrium}\} \ = \ \frac{1}{4} - \frac{\tau + \sigma + \varrho}{\pi} \ = \ \frac{1}{4} - \frac{7\epsilon}{5\pi} - \frac{14\epsilon^2}{25\pi} + O(\epsilon^3) : \tag{27}$$

it tends to a quarter and equals a quarter at no positive error rate. The values are 0.2499554, 0.2495542, 0.2455259 and 0.2037719 at $\epsilon = 10^{-4}, 10^{-3}, 10^{-2}, 10^{-1}$, all four below the threshold $\frac{1}{4}$ at which $\varrho$ wakes. The two edge slivers are the *ALLC* and *ALLD* shave $2\tau$, which is provable by hand; the two interior slivers are $2\sigma$, which is not, and at $\epsilon = 10^{-1}$ they are just under a quarter of the whole correction, $\sigma$ being $31\%$ of $\tau$ there.

**Above $\epsilon = \frac{1}{4}$ the cone loses more than the slivers.** Exactly four further residents hold equilibria arbitrarily far out inside $K_\epsilon$ at large error rates: 1680 and 1232, with their conjugates 63135 and 62687, none of them ever efficient. The whole constraint system of 1680 carries just three normal directions, identically in $\epsilon$, and the edge of its region that matters is perpendicular to the first of them: the region is unbounded exactly along the cone from

$$w_0(\epsilon) \ = \ \big(\,8\epsilon - 8\epsilon^2 - 1,\ \ 1 + 4\epsilon - 8\epsilon^2\,\big) \tag{28}$$

to $K_\epsilon$'s own edge $(\epsilon,\, 1-\epsilon)$, and the cross product of the two directions is $(4\epsilon - 1)(1 - 2\epsilon)^2$ up to a positive factor: the intrusion switches on at exactly $\epsilon = \frac{1}{4}$, stays on for every larger rate below $\frac{1}{2}$, and its width is

$$\varrho \ = \ \arctan \frac{(4\epsilon - 1)(1 - 2\epsilon)^2}{1 + 2\epsilon - 4\epsilon^2} \qquad \Big(\tfrac{1}{4} \leq \epsilon \leq \epsilon_b\Big). \tag{29}$$

The conjugate 63135 does the mirror image along $K_\epsilon$'s other edge. At $\epsilon_b$, the unique root of $8\epsilon^4 - 22\epsilon^3 + 25\epsilon^2 - 12\epsilon + 2$ in $(\frac{1}{4}, \frac{1}{2})$, which is $0.394170\ldots$, the direction $w_0$ reaches the far edge of the *mirror* sliver and the two intrusions merge: from there to $\frac{1}{2}$ the union runs from the sliver's inner edge all the way to $K_\epsilon$'s edge, the loss saturates at the whole gap between them, and

$$\varrho \;=\; \arctan \frac{2-13\epsilon+32\epsilon^2-36\epsilon^3+16\epsilon^4}{6-23\epsilon+38\epsilon^2-28\epsilon^3+8\epsilon^4} \qquad \left(\epsilon_b \le \epsilon < \tfrac{1}{2}\right), \tag{30}$$

the two branches agreeing at $\epsilon_b$. The second pair, 1232 and 62687, joins at exactly $\epsilon = \frac{1}{3}$, but its arcs lie inside the first pair's at every rate at which they exist, so $\varrho$ is the whole correction. Every statement in this paragraph is certified over its stated interval by the exact machinery of §7 — the onset factorisations and containments are polynomial identities, the persistence and nonemptiness are root-isolated sign conditions — and none of it is visible at the four error rates the rest of this paper quotes, which is precisely why an exact certification over the interval was worth having: the computed census at any sampled rate below $\frac{1}{4}$ cannot see $\varrho$ at all.

SI Figure 6**a** draws both statements, and SI Figure 6**d** draws what becomes of them at $\epsilon = 2/5$, past the threshold of the paragraph above. It is drawn at $\epsilon = 10^{-1}$ rather than at the paper's $10^{-4}$ for the reason the section is about: every angle here is a multiple of $\tau$, and at $10^{-4}$ all four of them are three orders of magnitude below one device pixel. The geometry does not change shape with $\epsilon$ — the same six residents keep the same two slivers and the boundaries are the same two pencils — so the page shows the structure at the largest of the four rates and gives the $10^{-4}$ widths in its legend. (Below $\epsilon = \frac{1}{4}$, that is: the four rates the paper quotes share one shape, and the further structure that appears beyond $\frac{1}{4}$ is the business of (28)–(30) alone.)

**The efficient measure is exact, and the whole of it is *ALLC*.** *ALLC* cooperates fully, so it is efficient at every game with $u + v \le 1$ (§5), and by (24) it is an equilibrium on a half-plane. Its efficient-equilibrium region is the intersection of the two, whose asymptotic cone runs from $180° + \tau$ to $315°$: an opening of exactly $135° - \tau$. Running the same union of recession cones over all 65536 residents returns the same measure — checked at twelve error rates spanning the interval, from $10^{-4}$ to 0.499 — so nothing adds an arc of directions to *ALLC*'s cone, only finitely many rays, which carry no measure, and for $0 < \epsilon < \frac{1}{2}$

$$\mu\{\text{no efficient Nash equilibrium}\} \;=\; \frac{5}{8} + \frac{1}{2\pi}\arctan\frac{\epsilon}{1-\epsilon}, \tag{31}$$

namely 0.6250159, 0.6251593, 0.6266076 and 0.6426116 at the four error rates. Two corollaries come with it. The wedge of the main text is *ALLC*'s own, up to a bounded offset contributed by *AON*$_2$, which is why its far field is decided by *ALLC* alone. And the Stag Hunt loses its efficient equilibrium exactly on the sector from $180°$ to $180° + \tau$, so the wedge along $v = 0$ noted in §5 is $\arctan[\epsilon/(1-\epsilon)]$ wide rather than merely of order $\epsilon$.

**A third share: where every equilibrium is efficient.** The same recession cones answer the converse question. A game has *all* of its equilibria efficient exactly when it lies in none of the sets on which some resident is Nash but not efficient, and each of those is $Z_\epsilon(p)$ cut by the switch line on whichever side $p$'s efficiency region is not — or the whole of $Z_\epsilon(p)$ when $p$ is efficient at no game. Taking the union of their cones and complementing,

$$\mu\{\text{every Nash equilibrium is efficient}\} \;=\; \frac{1}{8} + \frac{\tau}{2\pi}, \tag{32}$$

counting only the games that *have* an equilibrium in the numerator; the denominator is the whole circle, here as everywhere in this section. The surviving sector is $[\,270^\circ - \tau,\ 315^\circ\,]$ — from the antipode of *ALLD*'s boundary to the switch line — so it is the lower half of Harmony, widened by one tilt into the Stag Hunt. In the $\epsilon \to 0$ limit the same computation gives exactly $135^\circ$, or $\frac{3}{8}$: the whole Snowdrift quadrant together with that half of Harmony. **So this share falls by a factor of three the instant the error rate is positive**, and what it loses is the whole of the Snowdrift *sector*: far out in every one of its directions but finitely many, a game at $\epsilon > 0$ either has no equilibrium at all or — in the two wedges of width $\tau$ along the axes and in the two interior slivers — has equilibria of which none is efficient. Near the origin the picture is not that: at $(9/10, 1/20)$ and its mirror the entire equilibrium set is ten fully cooperating strategies, all of them efficient. The density is a statement about directions, not about every game. If the vacuous games are counted instead, the share is $\frac{3}{8} - (\tau + 2\sigma)/2\pi$, which is $\frac{3}{8}$ in the limit as well, since there no game lacks an equilibrium. Values, excluding the vacuous games: 0.1250159, 0.1251593, 0.1266076 and 0.1426116 at the four error rates.

**A fourth share, and with it the four kinds of game.** (27) and (31) are not disjoint but nested: a game with no equilibrium has no efficient one, so the first is *contained* in the second, and the difference of the two is therefore itself a set's measure — the games that do keep equilibria and yet not one of them attains $E_{\max}$, which is the dark grey of Main Figures 3 and 4, here read free of any window:

$$\mu\{\text{equilibria exist, and not one of them is efficient}\} \;=\; \frac{3}{8} + \frac{3\tau + 2\sigma + 2\varrho}{2\pi}, \tag{33}$$

namely 0.3750605, 0.3756051, 0.3810816 and 0.4388398 at the four error rates; measured against the directions that keep an equilibrium at all it is 0.5000509, just over one half. Its sector decomposes exactly, and every piece has already been met: the whole Prisoner's Dilemma, $90^\circ$; the half of Harmony above the switch line, $45^\circ$, where the optimum needs perfect alternation and no alternator holds a region at $\epsilon > 0$, so nothing efficient survives there but along finitely many rays; the wedge $[\,180^\circ, 180^\circ + \tau\,]$ of Stag Hunt that *ALLC*'s tilt costs, which is the wedge along $v = 0$ of §5; and inside Snowdrift the two edge wedges of width $\tau$ together with the two interior slivers of width $\sigma$ — joined, above $\epsilon = \frac{1}{4}$, by the two intrusion arcs of width $\varrho$ from (29). So the six residents of (26) — and beyond $\frac{1}{4}$ the four of (28) — keep an equilibrium arbitrarily far out and never an efficient one, and what the extra round of memory buys at infinity is inefficient stability. In the $\epsilon \to 0$ limit (33) is exactly 0: there every game carries an equilibrium and every game carries an efficient one, so the class is empty. This share therefore rises from nothing to three eighths at any positive error rate, the largest rise of the four kinds. As with every density here the statement is about directions and not about each game: at $\epsilon = 10^{-4}$ the Prisoner's Dilemma still carries efficient equilibria near the origin, 11 of its 19 at $(u, v) = (-\frac{1}{10}, \frac{1}{7})$, and loses them only in the far field.

With (33) the four kinds of game are disjoint and exhaust the plane,

$$\left[\tfrac{1}{4} - \tfrac{\tau+\sigma+\varrho}{\pi}\right] + \left[\tfrac{3}{8} + \tfrac{3\tau+2\sigma+2\varrho}{2\pi}\right] + \left[\tfrac{1}{4} - \tfrac{\tau}{\pi}\right] + \left[\tfrac{1}{8} + \tfrac{\tau}{2\pi}\right] = 1,$$

the third term being the games that carry efficient and inefficient equilibria at once: the sector $[\,180^\circ + \tau,\ 270^\circ - \tau\,]$, the Stag Hunt shaved by one tilt at each end, and the antipode of the empty cone $K_\epsilon$. It is not the whole of the Stag Hunt, and the two wedges it gives up are not a technicality: at $(u, v) = (-10^5, -5)$ there are twenty equilibria and not one of them is efficient. Main Figure 5 draws the partition: four disks, one for each row of Table 2, shaded in that table's own four colours, with each panel's four exact densities set as a ring outside its rim and its four expressions printed beneath. It is drawn at $\epsilon = 10^{-1}$ for the reason SI Figure 6 is.

**Table 2: The four kinds of game, for memory-2 and memory-1.** Each entry is a natural density: the fraction of the *directions* at infinity the set occupies, with the whole circle as denominator in every one, so the four are disjoint and every row sums to 1. $\tau = \arctan[\epsilon/(1-\epsilon)]$ is the angle by which a positive error rate tilts *ALLC*'s and *ALLD*'s boundaries off the axes, $\sigma$ is the width of *one* of the two interior slivers of (26), and $\varrho$ — identically zero for $\epsilon \le \frac{1}{4}$ and given by (29)–(30) beyond — is the width of one of the two further intrusion arcs of (28). The square beside each heading is the colour that kind is drawn in on Main Figures 4 and 5; the two greys are also Main Figure 3's two off-spectrum categories and SI Figure 6**a** and **b**'s fields, and the yellow is SI Figure 6**c**'s, so the table is a key to those pages too. The last two columns are the *same* for the two families at every positive error rate, being cut by *ALLC*, *ALLD* and the line $u + v = 1$, all three of which the families share; the first two differ, by the two interior slivers memory-1 does not hold — and, above $\epsilon = \frac{1}{4}$, by the two intrusion arcs — in opposite senses, so the two families agree exactly on the sum of that pair at every rate. Memory-2's third column and memory-1's first coincide as numbers because their cones are antipodes of equal opening. In the limit the two families agree on nothing: memory-1's four entries are the $\tau \to 0$ values of its own formulas, so memory-1 is continuous at $\epsilon = 0$ and memory-2 is not.

| | ■ no equilibrium | ■ none efficient | ■ both kinds | ■ all efficient |
|---|---|---|---|---|
| memory-2, $\epsilon \to 0$ | 0 | 0 | $\frac{5}{8}$ | $\frac{3}{8}$ |
| memory-1, $\epsilon \to 0$ | $\frac{1}{4}$ | $\frac{3}{8}$ | $\frac{1}{4}$ | $\frac{1}{8}$ |
| memory-2, $0 < \epsilon < \frac{1}{2}$ | $\frac{1}{4} - \dfrac{\tau+\sigma+\varrho}{\pi}$ | $\frac{3}{8} + \dfrac{3\tau+2\sigma+2\varrho}{2\pi}$ | $\frac{1}{4} - \dfrac{\tau}{\pi}$ | $\frac{1}{8} + \dfrac{\tau}{2\pi}$ |
| memory-1, $0 < \epsilon < \frac{1}{2}$ | $\frac{1}{4} - \dfrac{\tau}{\pi}$ | $\frac{3}{8} + \dfrac{3\tau}{2\pi}$ | $\frac{1}{4} - \dfrac{\tau}{\pi}$ | $\frac{1}{8} + \dfrac{\tau}{2\pi}$ |

One relation survives the partition, and one piece of arithmetic must still not be done. (31) minus (32) is exactly $\frac{1}{2}$ at every $\epsilon$, because the two sectors that keep something — an efficient equilibrium, and nothing but efficient equilibria — have openings $135^\circ - \tau$ and $45^\circ + \tau$, which sum to exactly half a turn, the second lying inside the first. But the two *sets* are disjoint, so that $\frac{1}{2}$ is the measure of nothing: what lies between *an* efficient equilibrium existing and *all* of them being efficient is 1 minus their sum, $\frac{1}{4} - \tau/\pi$, which is the third term above. The subtraction that gives (33) is legitimate for precisely the reason this one is not — there the smaller set is contained in the larger.

**All four jump at $\epsilon = 0$, and the four jumps cancel.** Running (21) on the exact whole-plane limit regions of §3 gives 0 for (27), 0 for (31) and hence 0 for (33): far out in every direction of the plane some strategy is an equilibrium, and some equilibrium is maximally efficient. So in the limit the four kinds stand at 0, 0, $\frac{5}{8}$ and $\frac{3}{8}$, and at every positive error rate at just under $\frac{1}{4}$, just over $\frac{3}{8}$, just under $\frac{1}{4}$ and just over $\frac{1}{8}$. Not one of the four is continuous at $\epsilon = 0$, and because both lists sum to 1 the four jumps cancel: the quarter and the three eighths gained by the two kinds that had nothing are exactly what coexistence gives up, $\frac{5}{8} \to \frac{1}{4}$, together with what universal efficiency gives up, $\frac{3}{8} \to \frac{1}{8}$. No direction leaves the plane; they are redistributed, and the traffic is almost all one way — the whole Snowdrift quadrant leaves universal efficiency for the first two kinds, and the Prisoner's Dilemma and the upper half of Harmony leave coexistence for the second, while the only movement in the other direction is a wedge of width $\tau$ at the far end of the Stag Hunt, which joins the games whose every equilibrium is efficient. The mechanism behind this jump is the one §5 records: above the switch line the optimum is perfect alternation, and at $\epsilon > 0$ not one alternator holds a two-dimensional region, so the 90° above that line — the upper half of the Prisoner's Dilemma and the upper half of Harmony — passes from coexistence to (33). The remaining 45°, the lower half of the Prisoner's Dilemma, transfers for a different reason: below the line efficiency needs full cooperation, and in the far field of the Prisoner's Dilemma no fully cooperating strategy is an equilibrium either. Together they are the 135° that passes entire. The two $\tau$ wedges of the Stag Hunt show the same asymmetry at the game level: at $(u, v) = (-10^{12}, -5 \times 10^7)$ the only equilibrium is *ALLD* and it is not efficient, while at $(-5 \times 10^7, -10^{12})$ it is *ALLC* and it is. This is the exact form of the statement that the limit's guarantee of efficiency is withdrawn rather than weakened by a positive error rate.

**The other endpoint, $\epsilon = \frac{1}{2}$, and why it is not a limit.** At $\epsilon = \frac{1}{2}$ every component of every strategy equals $\frac{1}{2}$, so all 65536 strategies are the one object that cooperates with probability $\frac{1}{2}$ whatever the history. Any pair of them meets the outcome frequencies $(\frac{1}{4}, \frac{1}{4}, \frac{1}{4}, \frac{1}{4})$, so $E(\mathbf{q}, \mathbf{p}) = (2 + u + v)/4$ for *every* pair: all sixteen deviation triples vanish identically, every region is the whole plane, every strategy is a Nash equilibrium of every game, and every one of the four shares displayed above is 0 — the whole circle becomes the third of the four kinds, since *ALLD* is an equilibrium at every game and efficient at none while *ALLC* and the alternators are equilibria and efficient on their own sides of the switch line, so efficient and inefficient equilibria coexist everywhere. This is exactly where (23) loses its force — $c_{\mathbf{q}} \equiv \frac{1}{2} \equiv 1 - \epsilon$, the factor in front of the bracket is identically zero, and $Z_\epsilon(\textit{ALLC})$ is the whole plane rather than a half-plane. So (31) is a statement about the open interval and *fails at the endpoint*: it predicts $\frac{3}{4}$ where the answer is 0, and the surviving opening jumps from 90.11° at $\epsilon = 0.499$ to the whole circle at $\epsilon = \frac{1}{2}$. (33) inherits that failure exactly, predicting $\frac{3}{4}$ against 0, and (32) fails too, predicting $\frac{1}{4}$ against 0. Only (27) survives the endpoint, and by accident: the tilt reaches

$45°$, so the cone $K_\epsilon$ closes completely, and the sliver width carries a factor $(1-2\epsilon)^2$, leaving 0. (27), (31) and (33) are therefore 0 at both ends of the interval and positive strictly between, while (32) runs from $\frac{3}{8}$ at $\epsilon = 0$ to 0 at $\epsilon = \frac{1}{2}$. All four are discontinuous at $\epsilon = 0$, and three of the four — every one but (27) — are discontinuous at $\epsilon = \frac{1}{2}$ as well.

**Memory-one, and why two of the four do not move.** The same four densities over the 16 binary memory-1 strategies are a different computation, not a restriction of this one. A memory-1 resident is an equilibrium when no memory-1 strategy earns more, so the deviation set is the family itself, all 16 of them, and $Z^{(1)}_\epsilon(p) \supseteq Z_\epsilon(p)$ for the same $p$: there are fewer rivals to beat. The two families are nested as strategy sets and *not* as answers, so any agreement between their measures is a fact and not a corollary. Computed exactly by the recession cones of this section — all 15 deviations per resident, built from the exact symbolic pair chain and classified over the whole plane, the answer is Table 2.

**Two of the four are identical, to every digit, at every error rate.** The reason is (24): both efficiency measures are decided by *ALLC* alone; *ALLC* belongs to both families; and its region is the same half-plane in either, because the factor $c_{\mathbf{q}} - (1-\epsilon)$ in (23) is non-zero for every rival in *either* family, so which rivals are available never enters. The same holds for *ALLD*. Nothing in either family adds an arc of directions to *ALLC*'s cone, so the two sectors $135° - \tau$ and $45° + \tau$ have the same measure in either family, and with them the two columns of Table 2 that involve efficiency on both sides: their three boundaries are *ALLC*'s line, *ALLD*'s line and the switch line, and the first two belong to both families while the third belongs to the game.

**The other two differ, and by exactly the two slivers — above $\epsilon = \frac{1}{4}$, the two slivers and the two intrusion arcs — once each way.** At $\epsilon > 0$ every full-dimensional memory-1 region has as its recession cone one of only three sets — $Z_\epsilon(\textit{ALLD})$, $Z_\epsilon(\textit{ALLC})$, or their intersection — because all sixteen carry only the two normal directions $(1-\epsilon, -\epsilon)$ and $(-\epsilon, 1-\epsilon)$ of (24). The far field of memory-1 is therefore decided by those two half-planes and by nothing else, and

$$\mu_1\{\text{no Nash equilibrium}\} \;=\; \frac{1}{4} - \frac{\tau}{\pi} \;: \tag{34}$$

the cone $K_\epsilon$ entire, with no sliver taken out of it. The memory-2 value (27) is smaller by exactly $(\sigma+\varrho)/180°$ of the plane, and that difference *is* what the extra round of memory buys in the far field. It is held by the six residents of (26) — above $\epsilon = \frac{1}{4}$, together with the four of (28) — and the two that hold the full-dimensional regions at every rate, 62143 and 688, are genuinely memory-2. Of the four that hold only lines, two — 61695 and 240 — are memory-1 strategies embedded in the memory-2 space, so memory-1 does keep an equilibrium arbitrarily far out inside the cone; but a line carries no angular measure, which is why (34) is the cone entire. So the whole gain of memory-2 over memory-1, measured at infinity, is two arcs of width $\sigma + \varrho$ — two slivers of width $\sigma$ at every rate the paper quotes: at $\epsilon = 10^{-4}$ that is 0.0000127 of the plane, and at $\epsilon = 10^{-1}$ still only 0.0110. The same $(\sigma + \varrho)/\pi$ that leaves memory-2's first share enters its second, since these arcs keep an equilibrium and never an efficient one, so (33) is larger for memory-2 by exactly the amount (27) is smaller, and one number verifies both. What the extra round of memory buys in the far field is therefore stability and never efficiency.

**Memory-one does not jump at $\epsilon = 0$, and that is the sharpest contrast of all.** Setting $\tau = 0$ in memory-1's four formulas in Table 2 gives $\frac{1}{4}$, $\frac{3}{8}$, $\frac{1}{4}$ and $\frac{1}{8}$, which are exactly the values the limit computation returns, and the same holds for (31), whose memory-1 value tends to the $\frac{5}{8}$ the limit returns. All four memory-1 densities are therefore continuous on $[0, \frac{1}{2})$, the endpoint $\epsilon = 0$ included, while all four memory-2 densities are discontinuous there — $0 \to \frac{1}{4}$, $0 \to \frac{3}{8}$, $\frac{5}{8} \to \frac{1}{4}$, $\frac{3}{8} \to \frac{1}{8}$. That is what the two rows of Main Figure 5 are for: memory-1's four densities depend on $\epsilon$ through $\tau$ alone, so its two panels differ only in that tilt, while the two memory-2 panels have almost nothing in common. The abundance the limit reports for memory-2 is thus an artefact of the limit in a precise sense: memory-1 never had it, and memory-2 loses it the instant a hand can tremble. One small asymmetry runs the other way and is worth recording, since the main text states the limit version: *TFT* is a memory-1 equilibrium at no game at all in the limit, but at every $\epsilon > 0$ its region is a single game.

**What is not claimed.** The set $K_\epsilon \setminus N_\epsilon$ of games inside the cone that do keep an equilibrium is *not* bounded: 52 residents are equilibria arbitrarily deep inside $K_\epsilon$, along rays that run into its interior, and 255 to 257 are equilibria somewhere inside it. The density (21) does not require boundedness — only that the asymptotic cone of the union meets the interior of $K_\epsilon$ outside the slivers in finitely many rays, a set of angular measure zero — and that is what is computed. All of it is exact, and §7 records the checks.

#### 7 Exact methods and verification

**One exact object per pair, and everything follows from it.** Every $\epsilon \to 0$ result in this paper rests on the Markov chain tree theorem evaluated exactly. For a pair of strategies the transition probability out of any of the sixteen states is a product of two factors from $\{\epsilon, 1-\epsilon\}$, so each entry of $I - P$ is a polynomial in $\epsilon$ of degree two with integer coefficients, and each tree-theorem numerator $N_i(\epsilon)$ — the $(i, i)$ minor of $I - P$, equal to the sum over spanning in-trees rooted at $i$ of the product of the fifteen edge probabilities — is a polynomial of degree at most 30 whose lowest-order coefficient counts the in-trees of minimal total tremble count. The limit distribution of §3 is the ratio of those integer counts over the roots of minimal order, the deviation triples of (14) follow as exact rationals, and no extrapolation, no floating point and no rational reconstruction enter anywhere.

**Exact reconstruction from one word-sized prime.** A coefficient of $N_i$ is bounded in magnitude by $4^{15}\binom{30}{15} \leq 1.67 \times 10^{17}$: each of the fifteen non-root states chooses one of its four outgoing edges, and each tree contributes $\epsilon^M(1-\epsilon)^{30-M}$. The kernel evaluates the sixteen minors at the 31 points $t = 2, \ldots, 32$ modulo the prime $2^{63} - 25$ and interpolates through a precomputed inverse Vandermonde; the true coefficients are integers below half the prime with a 27-fold margin, so the symmetric-range lift is the exact integer by construction rather than by accuracy. Per evaluation point one elimination of the singular $16 \times 16$ system yields all sixteen minors up to a common scale that a single $15 \times 15$ determinant fixes; rank-deficient points fall back to sixteen determinants. Four checks run on every pair in production and abort the task on failure: the coefficient bound, the tree-count range $1 \leq c_i \leq 4^{15}$, re-evaluation of every interpolated polynomial at a 32nd point against a direct elimination, and the numerator sum. None failed over the $4.295 \times 10^9$ limits of the census.

**Scale, and where it ran.** The kernel takes 1.6 ms per pair on an Apple M2 Pro (`gcc -O2`, all four checks on) and about 4.2 ms on the cluster nodes used. The full sweep — all 65536 residents, each against its 65535 deviations, with no early abort so that every constraint set is complete — ran as a 1024-task array on the Harvard Cannon cluster (job 41384987), 64 residents per task, 5034 core-hours in about five hours of wall clock; every task COMPLETED with empty stderr and a verified final record.

**Complete constraint sets, and geometry with no box.** For each resident every one of the 65535 deviation triples is reduced over the positive common denominator to a primitive integer triple and recorded, deduplicated with multiplicity and a representative deviation — a median of 1435 distinct half-planes per resident and never more than 12583 — together with the exact count of deviations whose triple is $(0, 0, 0)$: the ties of §4, 172 841 580 of them in all. Because the recorded sets are complete, the whole-plane geometry needs no bounding box, no selection pass and no outer-bound bookkeeping: each region is classified over the whole plane by one-dimension-down redundancy removal — a constraint is a facet exactly when the region meets its line in a set of positive length, an interval computed in exact rational arithmetic — and the census, the facet lines, the tie lines, the single-game stratum and the recession cones are read from that classification.

**Two routes, one census.** The census was computed twice, by two pipelines that share nothing past the definition of the chain, and they agree in every entry: 23861 residents full-dimensional, 3774 on a line, 4875 at a single game, 33026 empty; 44 tie lines; 254 distinct facet lines; and the agreement is per resident — facet sets, tie intervals with their endpoints, and single games — with no discrepancy of any kind. The exact tree-theorem route is the authority throughout this paper; the second route is the numerical cross-check of the next paragraph. Every downstream count that reads the regions file — the efficiency strata, the counts per game, the maxima, the densities — rests on the exact route's outputs.

**The independent numerical cross-check.** The second pipeline evaluates each pair chain in quadruple precision at $\epsilon = h,\ h/10,\ h/100$ and extrapolates,

$$\nu_0 \;=\; \frac{1000\,\nu(h/100) \;-\; 110\,\nu(h/10) \;+\; \nu(h)}{891}, \tag{35}$$

then snaps each coefficient to the nearest rational of denominator at most $10^7$. Snapping is reconstruction, not derivation — exact arithmetic on snapped inputs is only as good as the snap — which is why this route plays the role of check and never of authority. As a check it is a strong one: on every facet, tie line and single game of the arrangement the snapped coefficient equals the exact one, so a floating-point pipeline and an integer pipeline, sharing nothing but the chain, return the same census.

**The fixed-$\epsilon$ layer is symbolic in $\epsilon$.** The same interpolated polynomials are exact at *every* error rate, not only in the limit: $\nu_i(\epsilon) = N_i(\epsilon)/\sum_j N_j(\epsilon)$ identically, with the denominator positive on $(0, \frac{1}{2})$ as a sum of tree weights. The sixteen one-flip half-planes of a resident are integer-polynomial triples of degree at most 60, and a fixed-$\epsilon$ statement about all error rates at once is a sign condition on such polynomials, decided over the whole of $(0, \frac{1}{2})$ by exact root isolation. That is how the all-rate results of §6 are certified — the pencil identities behind (24), the sliver structure of (26) with the width identity for $\sigma$, the intrusion structure of (28)–(30) with its onsets at exactly $\frac{1}{4}$ and $\frac{1}{3}$, and the completeness statement that no resident beyond the ten named ones ever holds a positive-measure arc inside $K_\epsilon$: every resident carries a per-resident certificate, a polynomial identity or a two-generator Farkas certificate with root-isolated determinant signs, valid on the whole interval. The published fixed-$\epsilon$ half-planes (`tools/exactcoef.py`, exact rational arithmetic at any rational rate) were verified against the symbolic triples at $\epsilon = 10^{-4}$ for all 65536 residents, sixteen triples each, with no disagreement.

**Verification, in twelve parts.** Each check is against something the computation being checked does not itself produce. *(i)* The three closed forms of §3 — $\nu(\mathit{ALLC}, 160)$, $\nu(\mathit{ALLD}, 160)$, $\nu(160, 160)$ — are reproduced exactly. *(ii)* On 6084 pairs, structured and random, an algorithmically independent reference — fraction-free Bareiss elimination over $\mathbb{Z}[\epsilon]$ in unbounded integers, with no modular arithmetic and no interpolation — returns identical order-and-count data for all sixteen roots and the identical limit distribution, pair for pair. *(iii)* All 65536 exact self-play limits equal the published self-play layer exactly, and with them the 475 points and 229 rates of §2; conjugation symmetry and $w_2 = w_3$ hold exactly across the space. *(iv)* Results known independently come out right without being put in: *ALLD* is an equilibrium exactly on $u \leq 0$ and *ALLC* exactly on $v \leq 0$; the region of resident 36873 is exactly $\{u \leq 1\} \cap \{v \leq 2\}$, the memory-2 threshold $b/c \geq 3/2$ of Hilbe et al. read on the donation line. *(v)* The kernel builds and passes its smoke tests identically under two unrelated compilers on two architectures (Apple clang on ARM, Intel `icx` on x86). *(vi)* The four always-on per-pair checks above ran on all $4.295 \times 10^9$ census limits with no failure. *(vii)* The box-free geometry pass reproduces the published census resident by resident with no discrepancy, which is simultaneously the a-posteriori audit of the retired snap. *(viii)* The weakness certificates: every full-dimensional resident carries at least 351 deviations with identically zero gain triple, and the 918 residents with none are 903 empty regions and 15 single games, where tight vertex constraints decide. *(ix)* The symbolic fixed-$\epsilon$ triples agree with the published exact fixed-$\epsilon$ layer on all 1048576 one-flip half-planes at $\epsilon = 10^{-4}$. *(x)* The two exact sign-decision procedures — Sturm chains and Möbius–Descartes bisection

— agree on every certificate polynomial they were both run on. *(xi)* The corrected density (27) matches a direct exact computation of the no-equilibrium arc, assembled from the ten intruding residents' exact cones, at $\epsilon = 0.26, 0.35, 0.42, 0.49$ — both branches of $\varrho$ — to twelve digits, while the uncorrected form errs by up to twelve percent of itself there. *(xii)* The memory-1 census is symbolic in $\epsilon$ throughout and needs none of this; it is unchanged.

**How the exact geometric tests are carried out, and why nothing overflows.** In the limit arrangement the facet coefficients are integers of modulus at most 18, so a vertex is rational with denominator at most 237 and the membership test $a + ub + vd \leq 0$ becomes an integer test; the vertices and edges of (19) are evaluated that way, with no tolerance anywhere. The fixed-$\epsilon$ maximum of 1024 (§5) enumerates its pairwise intersections in homogeneous coordinates, which keeps every product bounded. A one-dimensional region is a *segment*: the two constraints that cut its endpoints are each tight at a single vertex, so neither is a facet of it, and the endpoints are carried explicitly. The memory-1 census is symbolic in $\epsilon$ throughout: each $4 \times 4$ stationary system is solved as an exact rational function of $\epsilon$ and the limit is taken symbolically, with no floating point anywhere.

**Where each number comes from.** Every number in this document is read from a file. Table 3 is the list; the exact census and its constraint sets are the ca1 outputs, with one caveat — a maximum-tracking file from which no nearest or farthest point may be quoted — carries over unchanged for `maxneps/`.

**The proof record, pinned.** Because the exact computation is the foundation of the paper, its record is archived as an auditable object. The code archive carries, under `exact/`: the kernel `limit_exact.c` with its validation record; the independent big-integer reference `ref_limit.py`; the certification layer (`allcert.py`, `sliver_cert.py`, `signdec.py`, `sturm.py`); the reduced results `ca1_regions.csv` and `ca1_selfplay.csv` with the tie and all-rate certificate tables; `MANIFEST.sha256`, the SHA-256 hashes of every file in the folder; and `CENSUS_SHA256`, the hashes of all 1024 raw census outputs (1.5 GB, archived separately and pinned by them), with the first raw output shipped in full as test data. A reader need not rerun the $4.295 \times 10^9$ pair limits: `make test` in that folder rebuilds the kernel from source, recomputes a slice of the census and checks it byte-for-byte against the pinned cluster record and that record's hash, re-derives the closed forms of §3 and the all-rate certificates of §6 for the ten named residents, and regenerates every headline count of §5 from the shipped regions file — about ten minutes on a laptop. The archive is tagged for this version of the paper, and the tag is what the paper cites.

**Memory-3 has its route written down.** Nothing in the exact pipeline is specific to sixteen states beyond its two bounds — the tree-polynomial degree and the coefficient bound — and both scale mechanically with the state count. A future census at memory-3, where $4^3 = 64$ states put symbolic elimination far out of reach, would use exactly this route: tree-theorem numerators, evaluation modulo word-sized primes, and complete constraint sets.

**Table 3: Where each number comes from.** The principal results of this supplement against the file that produces them. Paths are relative to the project root; `ca1/` is the exact tree-theorem census and the authority for every number; the extrapolation kit `ca4/` is the independent cross-check and agrees with it entry for entry.

| result | file |
|---|---|
| *the $\epsilon \to 0$ limit, exact* | |
| the complete deviation triples, ties and census | `cannon/ca1/out/` |
| the arrangement: dimensions, facets, tie lines, points | `Analysis/ca1_regions.csv` |
| self-play points, rates and the spectrum | `Analysis/ca1_selfplay.csv` |
| equilibria at a named game, all three strata | `tools/nashat.py` |
| which strategies can be efficient at all | `tools/effdim.csv` |
| *fixed $\epsilon$, exact and symbolic* | |
| the sixteen half-planes at any rational $\epsilon$ | `tools/exactcoef.py` |
| the symbolic-in-$\epsilon$ triples and all-rate certificates | `Analysis/allcert.py` |
| the sliver and intrusion certification | `Analysis/sliver_cert.py` |
| the census by dimension, and at the rational rate $2/5$ | `tools/epsdim.csv`, `SIFigure6/build_regions_frac.p` |
| the maximum number of equilibria at one game | `tools/epsmax.csv` |
| the efficient maximum, and its caveat above | `maxneps/` |
| *the four kinds of game of §6* | |
| the shares, the partition, and the witnesses | `tools/measures.csv`, `tools/fourshares.csv`, `tools/` |
| the same shares for memory-1, symbolic | `Main/Figures/M1/m1_nash.py` |

#### Supplementary References
#### References

[22] Boyd, R. Mistakes allow evolutionary stability in the repeated prisoner's dilemma game. *Journal of*

*Theoretical Biology* **136**, 47–56 (1989).

[42] Zhu, F. & Lin, F. Constant-memory strategies in stochastic games: best responses and equilibria. *arXiv preprint arXiv:2505.07008* (2025).

[43] Press, W. H. & Dyson, F. J. Iterated prisoner's dilemma contains strategies that dominate any evolutionary opponent. *Proceedings of the National Academy of Sciences* **109**, 10409–10413 (2012).

[44] Akin, E. The iterated prisoner's dilemma: good strategies and their dynamics. In *Ergodic Theory, Advances in Dynamical Systems*, 77–107 (de Gruyter Berlin, 2016).

[45] Hilbe, C., Traulsen, A. & Sigmund, K. Partners or rivals? strategies for the iterated prisoner's dilemma. *Games and Economic Behavior* **92**, 41–52 (2015).

[46] Glynatsi, N. E. & Knight, V. A. Using a theory of mind to find best responses to memory-one strategies. *Scientific Reports* **10**, 1–9 (2020).

[47] Puterman, M. L. *Markov decision processes: discrete stochastic dynamic programming* (Wiley, New York, 1994).

[56] Lorberbaum, J. M. D. No strategy is evolutionary stable in the repeated Prisoner's Dilemma. *Journal*

*of Theoretical Biology* **168**, 117–130 (1994).

[90] Colman, A. M. & Browning, L. Evolution of cooperative turn-taking. *Evolutionary Ecology Research*

**11**, 949–963 (2009).

[106] Lesigang, F., Hilbe, C. & Glynatsi, N. E. Characterisation of reactive Nash equilibria in repeated additive games. *arXiv preprint arXiv:2606.27653* (2026).

[107] Falmagne, G. The phase space of symmetric 2x2 games. *SocArXiv preprint 10.31235/osf.io/tgh9y* (2026).

[108] Robinson, D. & Goforth, D. *The topology of the 2x2 games: a new periodic table* (Routledge, London, 2005).

[109] Bruns, B. Names for games: Locating 2 x 2 games. *Games* **6**, 495–520 (2015).

[110] Leighton, F. & Rivest, R. Estimating a probability using finite memory. *IEEE Transactions on Information Theory* **32**, 733–742 (1986).

[111] Freidlin, M. I. & Wentzell, A. D. *Random perturbations of dynamical systems* (Springer, Berlin, 2012), 3rd edn.

[119] Silvestrov, D. & Silvestrov, S. *Nonlinearly perturbed semi-Markov processes* (Springer, Cham, 2017).

[125] Meyer, C. D. Sensitivity of the stationary distribution of a markov chain. *SIAM Journal on Matrix*

*Analysis and Applications* **15**, 715–728 (1994).

[126] Dayar, T. & Stewart, W. J. On the effects of using the grassmann–taksar–heyman method in iterative aggregation–disaggregation. *SIAM Journal on Scientific Computing* **17**, 287–303 (1996).

[127] Dehnert, C. *et al.* Prophesy: A probabilistic parameter synthesis tool. In *Computer Aided Verification (CAV 2015), Lecture Notes in Computer Science 9206*, 214–231 (Springer International Publishing, 2015).

[128] Quatmann, T., Dehnert, C., Jansen, N., Junges, S. & Katoen, J. P. Parameter synthesis for markov models: Faster than ever. In *Automated Technology for Verification and Analysis (ATVA 2016), Lecture Notes in Computer Science 9938*, 50–67 (Springer International Publishing, 2016).

[129] Baek, S. K., Jeong, H. C., Hilbe, C. & Nowak, M. A. Comparing reactive and memory-one strategies of direct reciprocity. *Scientific Reports* **6**, 25676 (2016).

[130] Nowak, M. A. Simple evolution drives direct reciprocity to maximum payoff in social dilemmas (2026). arXiv preprint arXiv:2608.06147.

#### Figure legends

**SI Figure 1: Binary memory-1 and memory-2 strategies in self play.** All $2^4 = 16$ BM1 and all $2^{16} = 65536$ BM2 strategies playing copies of themselves, in the limit $\epsilon \to 0$. **Top row:** the one-move distribution $(w_1, w_2, w_3, w_4)$ of (9) in the triangle — perfect alternation at the apex, mutual defection at the bottom left, mutual cooperation at the bottom right, so that $\gamma$ increases left to right exactly as it does on the axes beneath, and each vertex sits directly above its own value there. Marker colour is $\gamma$ and marker area is proportional to $\log_{10}$ of the number of strategies at the point, with a different constant in each panel because 16 and 65536 cannot share a scale; each carries its own size key. **Bottom row:** the $\gamma$ spectrum of each memory level on a logarithmic count, one line per realisable level of height equal to the number of strategies realising it. Grey bands mark the empty end intervals. BM1 realises six points and five levels, exactly the quarters, with nothing in $(0, 1/4)$ or $(3/4, 1)$. Left to right the six are $\gamma = 0$, $n = 3$ (*ALLD*, *Grim*, *Anti WSLS*); $\gamma = 1/4$, $n = 2$ (0100, 0010); $\gamma = 1/2$, $n = 2$ at the never-alternating point $(\frac{1}{2}, 0, 0, \frac{1}{2})$ (0001, 0111); $\gamma = 1/2$, $n = 4$ at the half-alternating point $(\frac{1}{4}, \frac{1}{4}, \frac{1}{4}, \frac{1}{4})$ (1100, *TFT*, *Anti TFT*, 0011); $\gamma = 3/4$, $n = 2$ (1101, 1011); and $\gamma = 1$, $n = 3$ (1110, *WSLS*, *ALLC*). A bit string $b_1b_2b_3b_4$ denotes $p_i = 1 - \epsilon$ where $b_i = 1$ and $p_i = \epsilon$ where $b_i = 0$. In self play $w_2 = w_3$ exactly (10), and for memory-1 — and only for memory-1 — exchanging $p_{CD}$ with $p_{DC}$ leaves the distribution unchanged, which is why *TFT* shares a point with 1100; the corresponding memory-2 swap changes the limit point for 24580 of the 65536. The six points and the five levels are the diagonal of Table 1 of Ref. (32), read as cooperation rates; Table 2 of Ref. (33) gives the same sixteen distributions in leading-order $\epsilon$ form. BM2 realises 475 distinct points and 229 levels, five of which $(0, \frac{1}{3}, \frac{1}{2}, \frac{2}{3}, 1)$ carry 63% of the strategies. Its empty end intervals are $(0, 1/10)$ and $(9/10, 1)$, each of width exactly $1/10$ and by a wide margin the largest gaps in the spectrum; the next largest is $1/42$, occurring twice as a conjugate pair, and the smallest is $1/4270 = 2.34 \times 10^{-4}$, also a conjugate pair. Memory-2 therefore raises the ceiling on sub-perfect monomorphic cooperation from $3/4$ to $9/10$. The count 7639 at $\gamma = 1$ is the memory-2 efficient-set size of Ref. (26); the remainder of the spectrum is, as far as we are aware, reported here for the first time.

**SI Figure 2: The same two families at $\epsilon = 1/100$.** SI Figure 1 redrawn at a positive error rate, with the same layout, the same mirrored barycentric map and the same colour scale, so the two pages may be compared panel for panel. Everything the limit page reports is a property of the limit, and this is what replaces it. **The ladder dissolves.** Memory-2's 475 points become 14300 and its 229 levels become 14241; memory-1's 6 and 5 become 11 and 11. The rungs are gone and what is left is a cloud with internal texture — dense horizontal bands, two spurs running along the base towards each corner, and a knot still riding the apex at perfect alternation. **The corners empty.** *ALLD* sits at $\gamma = \epsilon$ and *ALLC* at $\gamma = 1 - \epsilon$, so no strategy reaches a vertex and nothing at all lies outside $[\epsilon, 1 - \epsilon]$: that, and only that, is what the grey bands at the two ends of each spectrum mark. **The gap closes.** The limit's largest gaps, $(0, 1/10)$ and $(9/10, 1)$, are populated here — 2380 levels lie in $(9/10, 1-\epsilon)$ and 125 within $10^{-3}$ of the maximum — so memory-2's ceiling of $9/10$ on sub-perfect cooperation is a limit statement and nothing more. The widest surviving gap is 0.0117, an order of magnitude narrower and no longer at the top: it runs from 0.19279 to 0.20450, and by conjugation from 0.79550 to 0.80721. **What survives.** $w_2 = w_3$ still holds exactly (10), so the distributions stay in the triangle; conjugation is still an exact symmetry of the spectrum, verified here as an equality of multisets rather than of counts; and the degeneracy does not simply dissolve — 14300 points is still only 22% of 65536, the largest class holds 864 strategies and only 470 are alone. Memory-1's eleven levels are 0.0100, 0.0149, 0.0294, 0.2550, 0.4951, 0.5000, 0.5049, 0.7450, 0.9706, 0.9851, 0.9900; six of them lie within 0.03 of an end and two within 0.005 of $1/2$, so that panel cannot resolve them all at this width and the numbers are given here instead. Marker area is proportional to $\log_{10}$ of 10 times the number of strategies at a point, not to $\log_{10}$ of the number: at a positive error rate 470 memory-2 points and 8 memory-1 points carry a single strategy, and $\log_{10} 1 = 0$ would draw them at zero size. For the same reason each spectrum line is drawn from the floor of its axis rather than from 1. Every stationary distribution behind this page is solved in exact rational arithmetic and two points are the same point when their 4-tuples are equal as fractions — a count of distinct points is a question about ties, and a float pass puts the memory-2 count anywhere between 14301 and 17575 depending on where it is rounded.

**SI Figure 3: The count, the efficient count and the share, at the two larger error rates.** The three quantities of Main Figures 2, 3 and 4 — all Nash equilibria at a game, the efficient ones among them, and the fraction of a game's equilibria that are efficient — over the two error rates the main text does not draw. Rows are the three quantities and columns are $\epsilon = 10^{-2}$ and $10^{-1}$; the disk and the map are Main Figure 2's; the switch line $u + v = 1$ is drawn here in crimson, as on Main Figures 3 and 4. Each row has one bar, shared by both columns, so a row is read *across*: what a larger error rate does to a single quantity. The two count rows are logarithmic and their bars stop at the whole-plane maximum *certified* in §5, 1024 and 245, not at the largest value the drawing itself reaches — a field whose maximum sits deep in the far field is understated by any evaluation on a grid, and a bar told its range by the picture would disagree with the text. Light grey marks games with no equilibrium of any kind and dark grey games that have equilibria but not one that is efficient, the same pair Main Figures 3 and 4 use; the disk carries no rim.

Every panel is the field evaluated exactly at each of $3200^2$ pixel centres and drawn as filled contour polygons, so the page carries no raster and no value on it is interpolated, smoothed or averaged; each boundary is a polygonal trace of the true straight facet lines, to one part in 3200 of a disk. *The arrangement is nevertheless finer than the page, and these panels do not pretend otherwise.* At the printed size the map puts 19.6 points on one unit of $u$ at the origin, and along the segment $|u| \leq 1$ of the $u$ axis alone there are 441 region boundaries at $\epsilon = 10^{-2}$ and 451 at $\epsilon = 10^{-1}$: their median separation is $3.0 \times 10^{-4}$ at the smaller rate, an eighth of a drawn pixel, and 78% of the gaps there — 46% at $\epsilon = 10^{-1}$ — are narrower than one pixel. The median band that survives onto the page is accordingly 1.08 points wide in the top right panel and 0.29 in the middle left, and a quarter of the bands in the $\epsilon = 10^{-2}$ column are narrower than 0.15 points, which is 0.05 mm. The terracing near the origin and along the two quadrant axes is therefore real structure compressed below the width of a printed line. It records how densely the levels change there and does not depict individual regions: read these panels for where a level lies and how the strata are arranged, not for the width or the exact position of any single band.

What the two columns show is that existence is nearly settled once $\epsilon$ is positive at all while efficiency is not. The full-dimensional stratum holds at 1050 at both, and both rates reach the certified maxima 1024 and 245, so the counts do not fall between them. What does move is the dark grey: the games that carry equilibria but not one efficient one rise from 32.6% of them at $\epsilon = 10^{-4}$ to 33.3% and then 40.7% — shares of the disk as drawn, and of the games in it that keep an equilibrium, not the density (33), which is the same category read at infinity and over all directions — while the quadrant carrying nothing at all shrinks from 21.7% of the disk to 21.5% and then 18.6%. A larger error rate costs fewer games their equilibria outright and more of them their efficient ones.

**SI Figure 4: Where a memory-one strategy is an equilibrium.** Main Figure 2 for the $2^4 = 16$ binary memory-1 strategies: rows are the dimension of the set of games at which a strategy is an equilibrium, columns the vanishing-error limit and $\epsilon = 10^{-4}$. Two things differ from the memory-2 page and both follow from 16 against 65536. First, the objects are named rather than coloured: four rays and two or three games, each the whole equilibrium set of one strategy — of two where 1100 and 0011 share $(0, 0)$ — so a colour scale would carry almost no information. Second, the count field runs 0 to 8 rather than 299 to 22069, so its bar gives every integer a band and the two field panels share it; and grey, marking the games with no equilibria at all, is needed in the *limit* as well as at $\epsilon > 0$, which memory-2 never requires. The two field panels carry no rim; the four sparse ones keep theirs, being drawn on white paper with nothing else to bound them. **a**, **d**, the same eight strategies hold a full-dimensional region at both error rates, and seven of them hold very nearly the same region — each moves by under a tenth of a percentage point of the square $|u|, |v| \leq 8$. The eighth, 1110, is the whole of the difference: in the limit its region is the half-plane $v \leq 0$, and at $\epsilon = 10^{-4}$ it is that half-plane pushed out beyond $u \approx -1/2\epsilon$, so that inside the drawn window it holds nothing at all. That is why the count is unchanged over half the square and one lower over the other half, and why the maximum falls from 8 to 7: the region does not shrink, it *recedes*, on the same $1/2\epsilon$ scale as *TFT*'s single game in **f**. **b**, **e**, the four rays, unchanged but for a shift of order $\epsilon$. **c**, **f**, the single games: 1100 and 0011 share the game $(0, 0)$ and *Anti TFT* holds $(\frac{1}{2}, -\frac{1}{2})$, and at $\epsilon = 10^{-4}$ *TFT* joins them at $(-4999, 4999)$ — on the donation line, at cost $c = 1 - 2\epsilon$, which recedes to infinity as $\epsilon$ falls and is why *TFT* is an equilibrium at no game whatever in the limit.

**SI Figure 5: What a positive error rate costs the memory-one efficient equilibria.** Main Figures 3 and 4 for the 16 binary memory-1 strategies, as one page: row 1 the number of efficient equilibria at each game, row 2 the share of that game's equilibria which are efficient, and in both rows the vanishing-error limit on the left and $\epsilon = 10^{-4}$ on the right. Light grey marks games with no equilibrium of any kind and dark grey games that have equilibria but not one that is efficient, the pair Main Figures 3 and 4 use; no disk carries a rim. Both bars are discrete, and the lower one is discrete for a reason worth stating: over 16 strategies the share is a ratio of two very small integers and takes only ten values in all, seven in the limit and six at $\epsilon = 10^{-4}$, so the bar names each of them exactly rather than implying a continuum. It is coloured by rank, which Main Figure 4's is not — there the same quantity takes 3656 values and a gradient is the honest picture. Above the crimson line $u + v = 1$ the page is grey at every game, at both error rates: dark where equilibria survive without an efficient one, light over the part of Snowdrift beyond *WSLS*'s island where none survives at all. That is the sharpest memory-one statement in this supplement drawn rather than asserted, because above that line the optimum is perfect alternation, no binary memory-1 strategy alternates more than half the time, and so memory-one has no efficient equilibrium there at any error rate. What a positive error rate costs memory-one is depth and not territory. By area of the disk as drawn, the three categories — no equilibrium, equilibria but none efficient, at least one efficient — stand at 23.16%, 28.44% and 48.39% in the limit and at 23.16%, 28.45% and 48.39% at $\epsilon = 10^{-4}$, the same to five thousandths of a percentage point, the $O(\epsilon)$ motion of the boundaries and nothing else. What changes is the count: the most efficient equilibria any game carries falls from 3 to 2, and the whole field steps down with it. Read at infinity instead of on the disk, the same three categories are 0.249968, 0.375048 and 0.374984 — the first two the memory-1 row of Table 2, the third its last two columns together — the gap being the window and nothing else. Memory-2 loses a quarter of the plane's directions to games with no equilibrium at all at the same error rate; memory-1 loses none of it, having already had that hole in the limit.

**SI Figure 6: The three shares of the plane, and the lines that fix them.** The sets of §6 themselves, one to a panel, where Main Figure 5 draws the four kinds they make — and, in the fourth panel, what the first becomes above the threshold $\epsilon = \frac{1}{4}$. All four panels draw the disk of Main eq. (2) at $k = 4$, the first two in Main Figures 3 and 4's two greys; the box beside each title is that panel's colour, and every share here is angular, read at the rim and not from the visible area. **a**, the games at which not one of the 65536 strategies is a Nash equilibrium, (27). They lie inside the cone cut by *ALLC*'s boundary, blue, and *ALLD*'s, purple, which at a positive error rate are exactly $(1 - \epsilon)v \leq \epsilon u$ and $(1 - \epsilon)u \leq \epsilon v$. Both pass through the centre, so the cone that contains this set is the same at every $k$; the set itself is not, since the white centre is the near field, where equilibria survive. Two slivers, green, keep an equilibrium arbitrarily far out — three strategies in one and their mirror images in the other, the same six at every error rate below $\frac{1}{4}$ — beyond it four more join, (28) — (26). **b**, the games at which no equilibrium attains $E_{\max}$, (31). Its two greys are its two shares: the lighter is **a**, the darker the games that carry equilibria without an efficient one, (33). The one kind of the four that no panel here draws on its own is the third, where efficient and inefficient equilibria coexist. The surviving white wedge is *ALLC*'s own, bounded below by its boundary, through the centre, and above by the crimson switch line $u + v = 1$, which is not. **c**, the games that carry equilibria and all of them are efficient, (32), in the yellow of Main Figure 4's fraction one; it runs from the antipode of *ALLD*'s boundary to the switch line, and in the limit that sector is instead the whole Snowdrift quadrant together with the same half of Harmony, $\frac{3}{8}$ of the directions. **d**, panel **a**'s set at $\epsilon = 2/5$, past both the onset $\frac{1}{4}$ of (28) and the bridging point $\epsilon_b \approx 0.394$ of (30): the amber dotted rays are the intrusion edges $w_0$ and its mirror, the residents 1680 and 63135 hold equilibria arbitrarily far out between those rays and the boundary lines, each intrusion has met its sliver, and the grey that remains — (27) with $\varrho$ awake — is the wedge between the two inner sliver edges. Panels **a** to **c** are drawn at $\epsilon = 10^{-1}$, where $\tau = 6.34°$ and $\sigma = 1.98°$; at $10^{-4}$ they are $0.0057°$ and $0.0023°$; at panel **d**'s rate $\tau = 33.69°$, $\sigma = 1.33°$ and $\varrho = 1.13°$. Every line is exact; the fields are rasters.

**SI Figure 1**

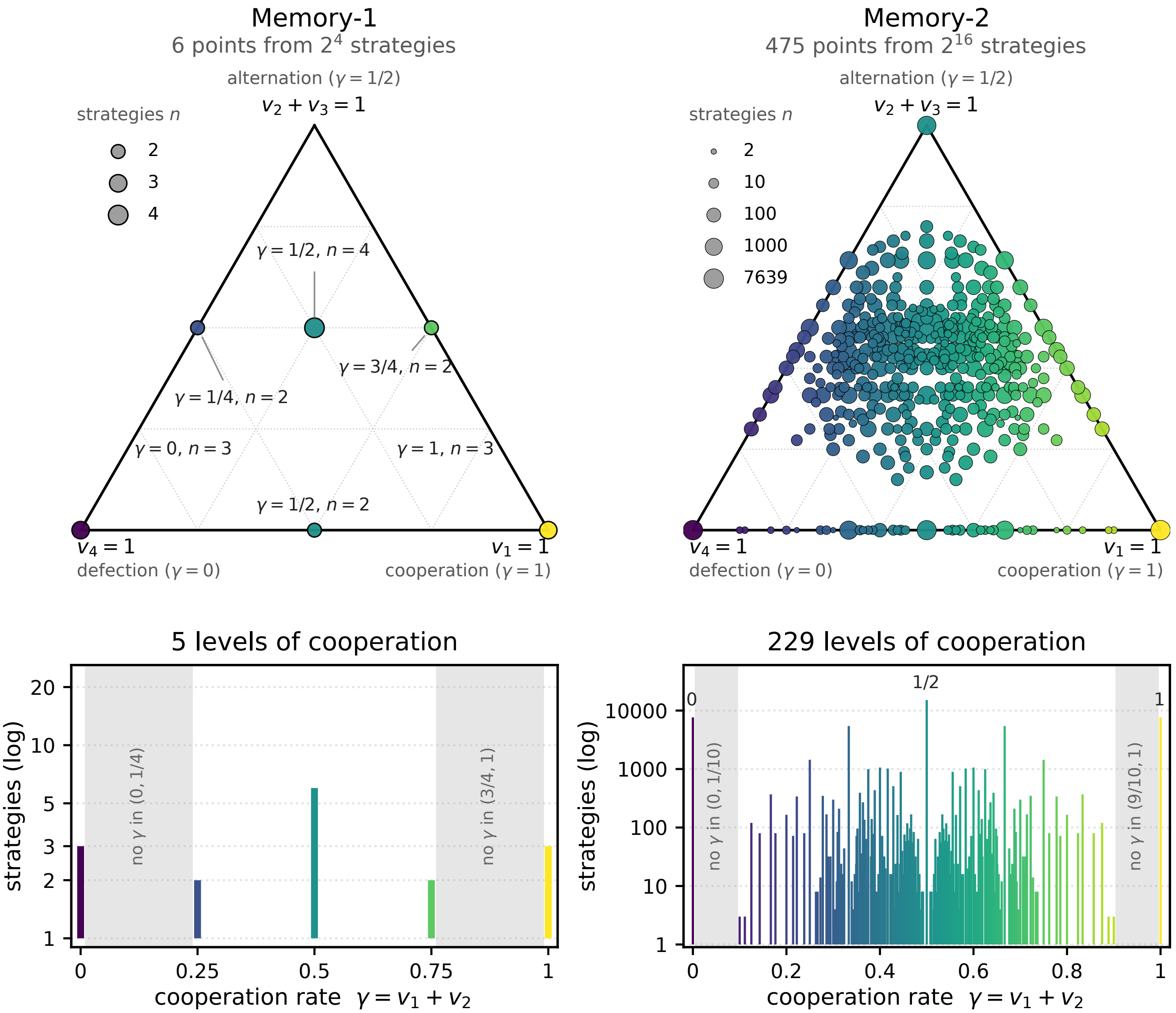

Memory-1
6 points from $2^4$ strategies
alternation ($\gamma = 1/2$)
$v_2 + v_3 = 1$
strategies $n$
2
3
4
$\gamma = 1/2$, $n = 4$
$\gamma = 3/4$, $n = 2$
$\gamma = 1/4$, $n = 2$
$\gamma = 0$, $n = 3$
$\gamma = 1$, $n = 3$
$\gamma = 1/2$, $n = 2$
$v_4 = 1$
defection ($\gamma = 0$)
$v_1 = 1$
cooperation ($\gamma = 1$)
Memory-2
475 points from $2^{16}$ strategies
alternation ($\gamma = 1/2$)
$v_2 + v_3 = 1$
strategies $n$
2
10
100
1000
7639
$v_4 = 1$
defection ($\gamma = 0$)
$v_1 = 1$
cooperation ($\gamma = 1$)
5 levels of cooperation
strategies (log)
20
10
5
3
2
1
no $\gamma$ in (0, 1/4)
no $\gamma$ in (3/4, 1)
0
0.25
0.5
0.75
1
cooperation rate $\gamma = v_1 + v_2$
229 levels of cooperation
strategies (log)
10000
1000
100
10
1
0
1/2
1
no $\gamma$ in (0, 1/10)
no $\gamma$ in (9/10, 1)
0
0.2
0.4
0.6
0.8
1
cooperation rate $\gamma = v_1 + v_2$

**SI Figure 2**

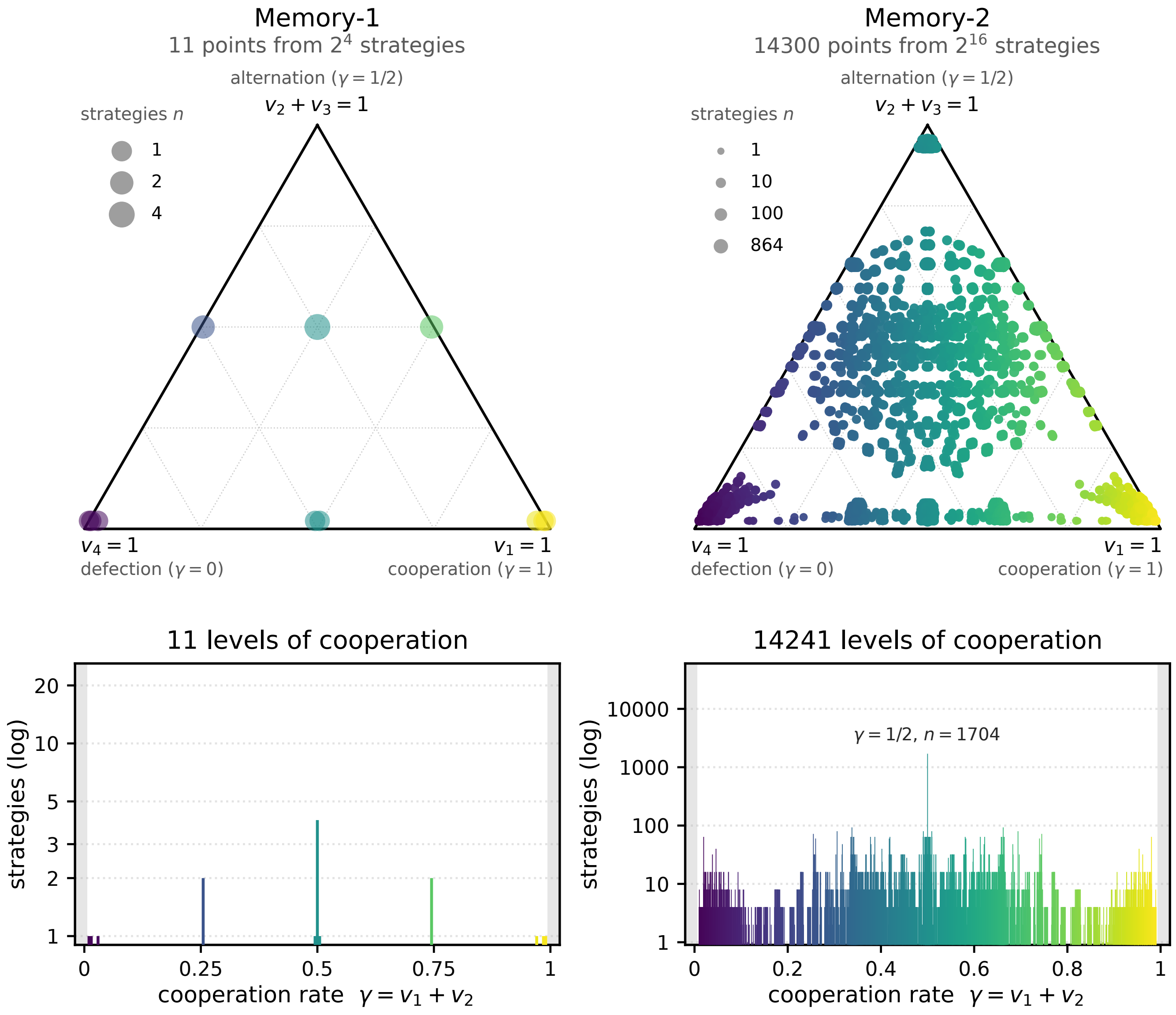
Memory-1
11 points from $2^4$ strategies
alternation ($\gamma = 1/2$)
$v_2 + v_3 = 1$
strategies $n$
1
2
4
$v_4 = 1$
defection ($\gamma = 0$)
$v_1 = 1$
cooperation ($\gamma = 1$)
Memory-2
14300 points from $2^{16}$ strategies
alternation ($\gamma = 1/2$)
$v_2 + v_3 = 1$
strategies $n$
1
10
100
864
$v_4 = 1$
defection ($\gamma = 0$)
$v_1 = 1$
cooperation ($\gamma = 1$)
11 levels of cooperation
strategies (log)
20
10
5
3
2
1
0
0.25
0.5
0.75
1
cooperation rate $\gamma = v_1 + v_2$
14241 levels of cooperation
strategies (log)
10000
1000
100
10
1
$\gamma = 1/2$, $n = 1704$
0
0.2
0.4
0.6
0.8
1
cooperation rate $\gamma = v_1 + v_2$

**SI Figure 3**

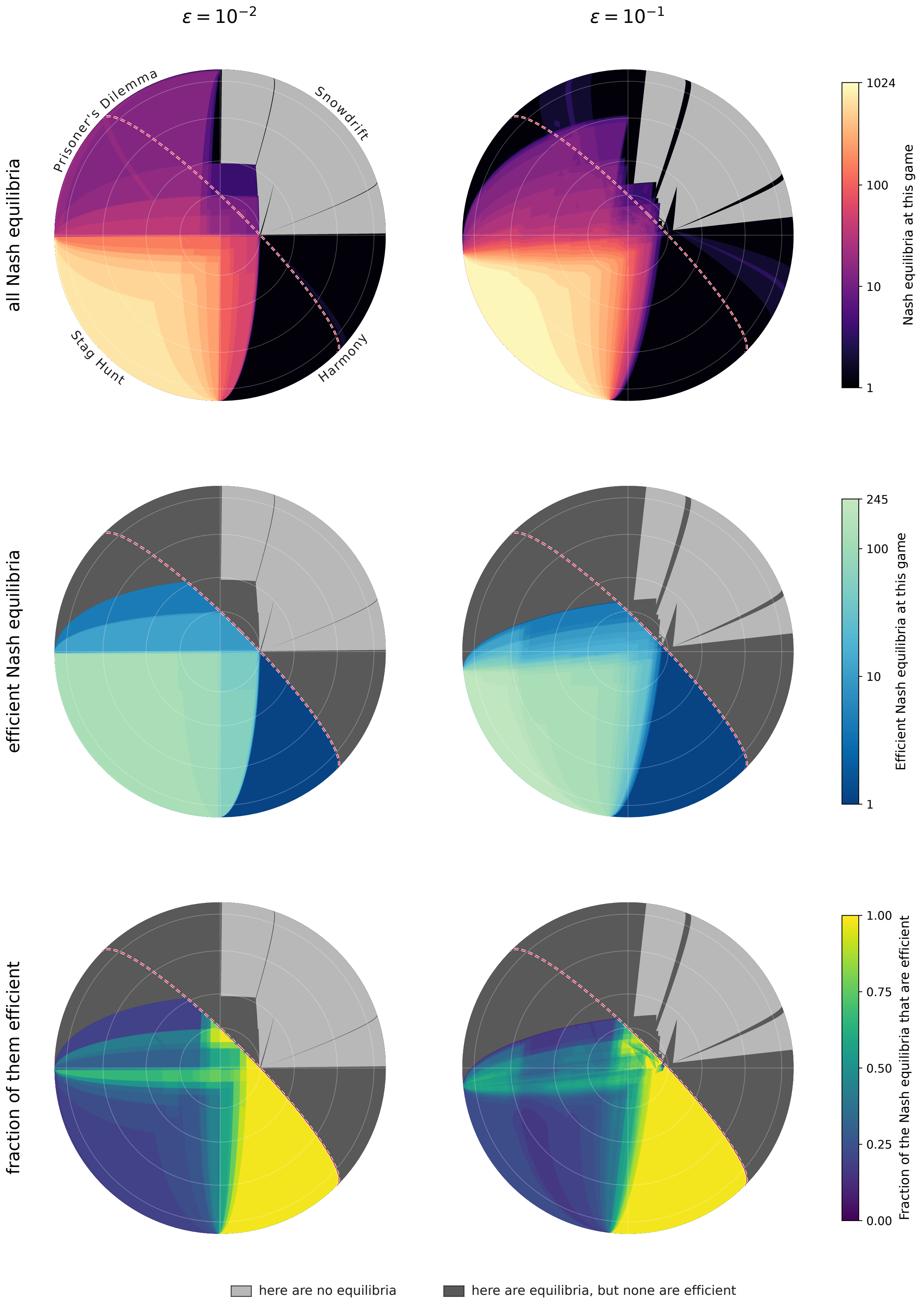

$\varepsilon = 10^{-2}$
$\varepsilon = 10^{-1}$
all Nash equilibria
efficient Nash equilibria
fraction of them efficient
Prisoner's Dilemma
Snowdrift
Stag Hunt
Harmony
Nash equilibria at this game
1024
100
10
1
Efficient Nash equilibria at this game
245
100
10
1
Fraction of the Nash equilibria that are efficient
1.00
0.75
0.50
0.25
0.00
here are no equilibria
here are equilibria, but none are efficient

**SI Figure 4**

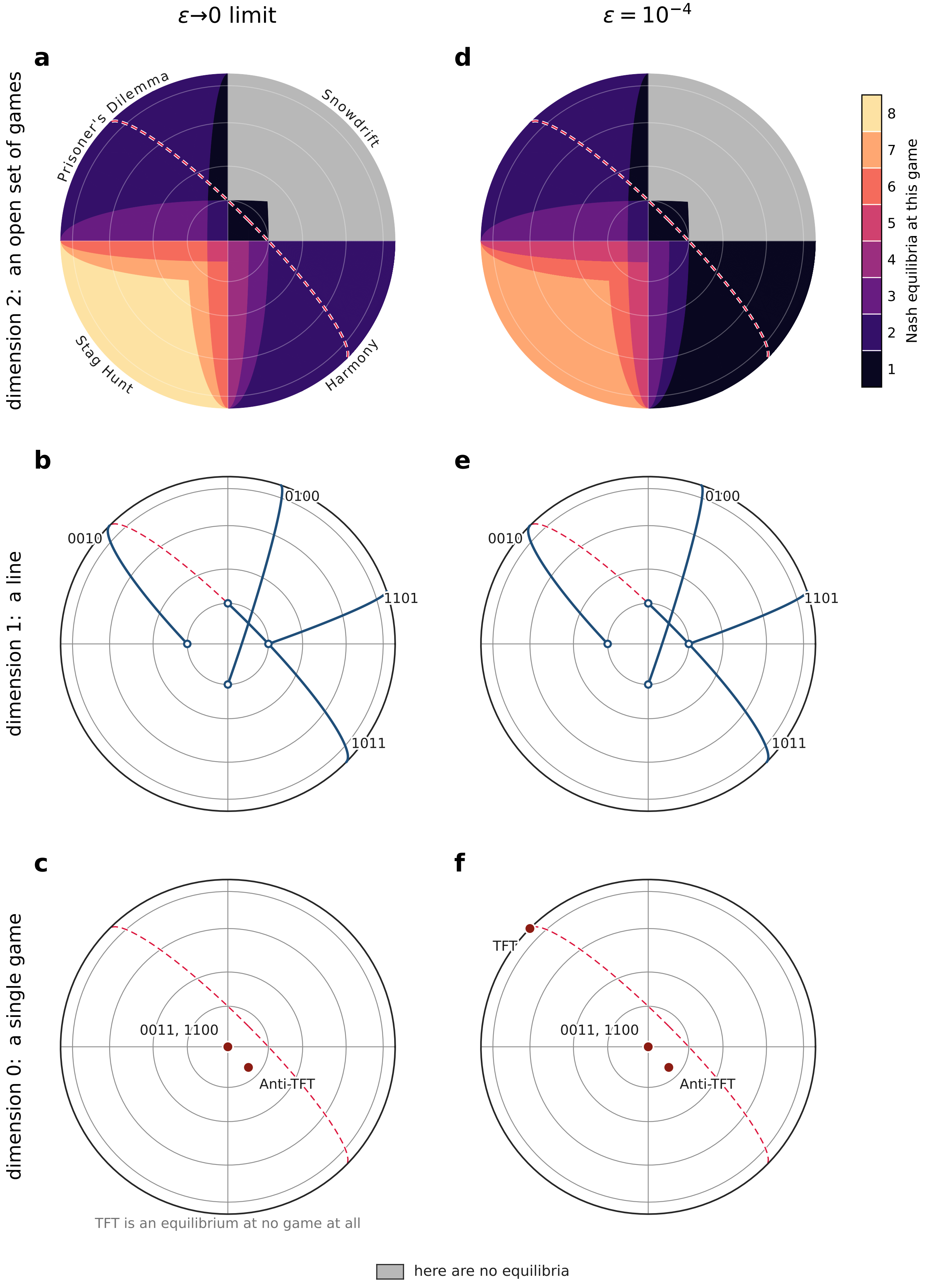

ε→0 limit
ε = 10^−4
a
d
dimension 2: an open set of games
Prisoner's Dilemma
Snowdrift
Stag Hunt
Harmony
Nash equilibria at this game
8
7
6
5
4
3
2
1
b
e
dimension 1: a line
0100
0010
1101
1011
c
f
dimension 0: a single game
TFT
0011, 1100
Anti-TFT
TFT is an equilibrium at no game at all
here are no equilibria

**SI Figure 5**

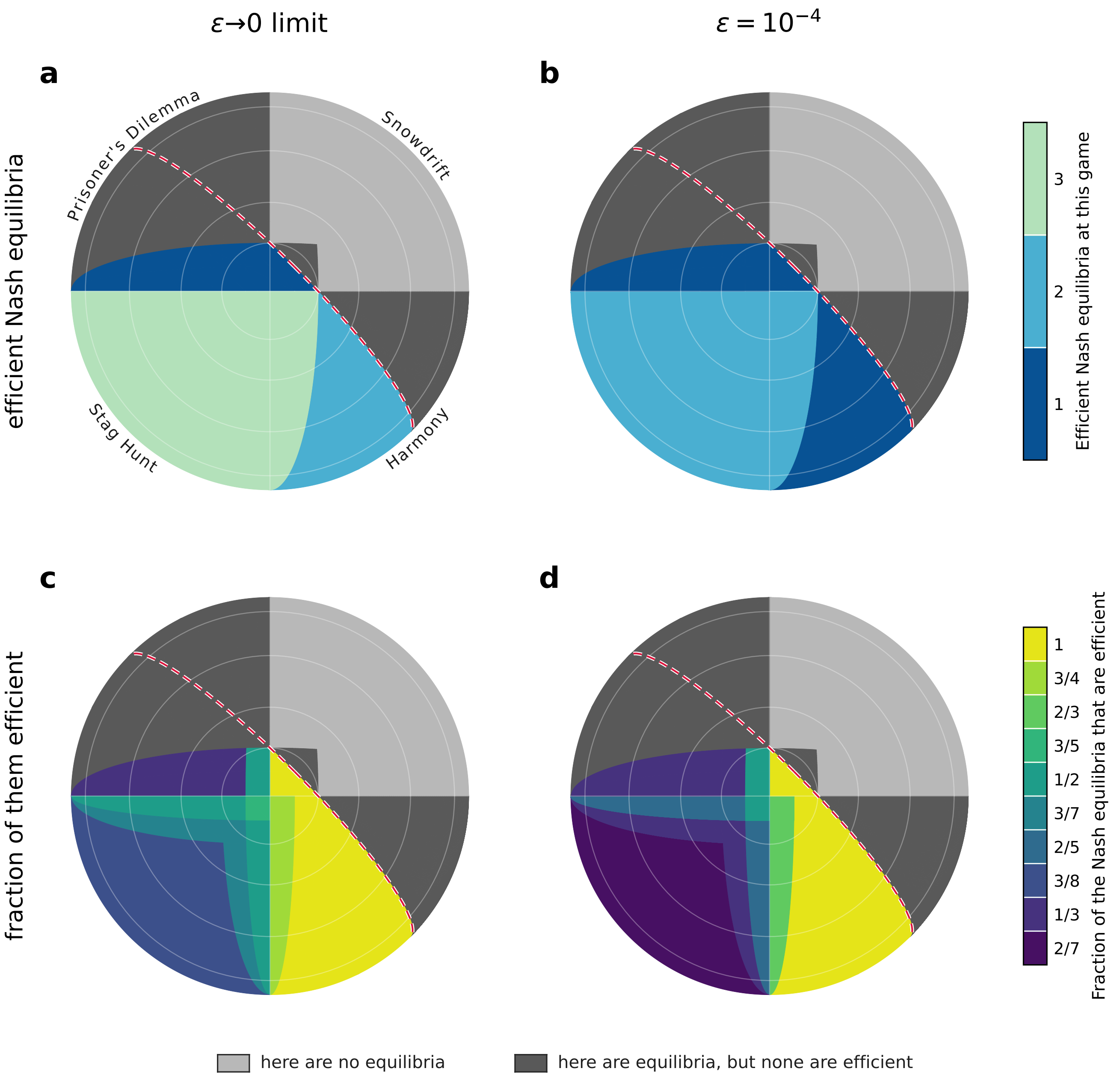

ε→0 limit
ε = 10^−4
a
b
c
d
efficient Nash equilibria
fraction of them efficient
Prisoner's Dilemma
Snowdrift
Stag Hunt
Harmony
Efficient Nash equilibria at this game
3
2
1
Fraction of the Nash equilibria that are efficient
1
3/4
2/3
3/5
1/2
3/7
2/5
3/8
1/3
2/7
here are no equilibria
here are equilibria, but none are efficient

### SI Figure 6

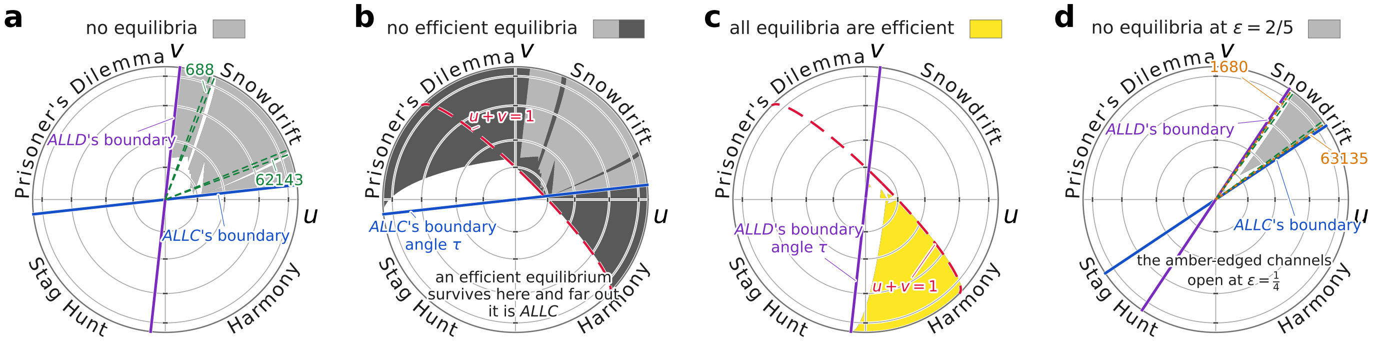


$\mu = \frac{1}{4} - (\tau + \sigma)/\pi$

$\mu = \frac{5}{8} + \tau/2\pi$

$\mu = \frac{1}{8} + \tau/2\pi$

$\mu = \frac{1}{4} - (\tau + \sigma + \varrho)/\pi$
